# Longer scans boost prediction and cut costs in brain-wide association studies

**DOI:** 10.1101/2024.02.16.580448

**Authors:** Leon Qi Rong Ooi, Csaba Orban, Shaoshi Zhang, Thomas E. Nichols, Trevor Wei Kiat Tan, Ru Kong, Scott Marek, Nico U.F. Dosenbach, Timothy Laumann, Evan M Gordon, Kwong Hsia Yap, Fang Ji, Joanna Su Xian Chong, Christopher Chen, Lijun An, Nicolai Franzmeier, Sebastian Niclas Roemer, Qingyu Hu, Jianxun Ren, Hesheng Liu, Sidhant Chopra, Carrisa V. Cocuzza, Justin T. Baker, Juan Helen Zhou, Danilo Bzdok, Simon B. Eickhoff, Avram J. Holmes, B. T. Thomas Yeo, Alzheimer’s Disease Neuroimaging Initiative

## Abstract

A pervasive dilemma in brain-wide association studies (BWAS) is whether to prioritize functional MRI (fMRI) scan time or sample size. We derive a theoretical model showing that individual-level phenotypic prediction accuracy increases with sample size and total scan duration (sample size × scan time per participant). The model explains empirical prediction accuracies extremely well across 76 phenotypes from nine resting-fMRI and task-fMRI datasets (R^2^ = 0.89), spanning a wide range of scanners, acquisitions, racial groups, disorders and ages. For scans ≤20 mins, prediction accuracy increases linearly with the logarithm of total scan duration, suggesting interchangeability of sample size and scan time. However, sample size is ultimately more important than scan time in determining prediction accuracy. Nevertheless, when accounting for overhead costs associated with each participant (e.g., recruitment costs), to boost prediction accuracy, longer scans can yield substantial cost savings over larger sample size. To achieve high prediction performance, 10-min scans are highly cost inefficient. In most scenarios, the optimal scan time is ≥20 mins. On average, 30-min scans are the most cost-effective, yielding 22% cost savings over 10-min scans. Overshooting is cheaper than undershooting the optimal scan time, so we recommend aiming for ≥30 mins. Compared with resting-state whole-brain BWAS, the most cost-effective scan time is shorter for task-fMRI and longer for subcortical-cortical BWAS. Standard power calculations maximize sample size at the expense of scan time. Our study demonstrates that optimizing both sample size and scan time can boost prediction power while cutting costs. Our empirically informed reference is available for future study planning: WEB_APPLICATION_LINK

## Introduction

A fundamental question in systems neuroscience is how individual differences in brain function are related to common variation in phenotypic traits, such as cognitive ability or physical health. Following recent work (Marek et al., 2022), we define brain wide association studies (BWAS) as studies of the associations between phenotypic traits and common inter-individual variability of the human brain. An important subclass of BWAS seeks to predict individual-level phenotypes using machine learning. Individual-level prediction is important for addressing basic neuroscience questions and critical for precision medicine (Finn et al., 2015; Gabrieli et al., 2015; Woo et al., 2017; Bzdok et al., 2019; Eickhoff et al., 2019; Varoquaux et al., 2019).

Many BWAS are underpowered, leading to low reproducibility and inflated prediction performance (Button et al., 2013; Arbabshirani et al., 2017; Bzdok et al., 2018; Kharabian Masouleh et al., 2019; Elliott et al., 2020; Poldrack et al., 2020). Larger sample sizes increase reliability of brain-behavior associations (Tian et al., 2021; Chen et al., 2023) and individual-level prediction accuracy (He et al., 2020; Schulz et al., 2020). Indeed, reliable BWAS typically requires thousands of participants (Marek et al., 2022), although certain multivariate approaches might reduce sample size requirements (Chen et al., 2023).

In parallel, other studies have emphasized the importance of longer fMRI scan time per participant during both resting and task states, which leads to improved data quality and reliability (Mumford et al., 2008; Birn et al., 2013; Nee, 2019; Noble et al., 2019; Elliott et al., 2020; Lynch et al., 2020; G. Chen et al., 2022), as well as new insights into the brain (Laumann et al., 2015; Newbold et al., 2020; Gordon et al., 2023; Lynch et al., 2024). When sample size is fixed, increasing resting-state fMRI scan time per participant improves individual-level prediction accuracy of some cognitive measures (Feng et al., 2023).

Therefore, in a world with infinite resources, fMRI-based BWAS should maximize both sample size and scan time for each participant. However, in reality, BWAS investigators have to decide between scanning more participants (for a shorter duration), or fewer participants (for a longer duration). Furthermore, there is a fundamental asymmetry between sample size and scan time per participant because of inherent overhead cost associated with each participant, which can be quite substantial, e.g., when recruiting from a rare population. Surprisingly, the exact trade-off between sample size and scan time per participant has not been comprehensively characterised. This trade-off is not only relevant for small-scale studies, but also important for large-scale data collection, given competing interests among investigators and limited participant availability.

Here, we systematically characterize the effects of sample size and scan time of fMRI on BWAS prediction accuracy, using the Adolescent Brain and Cognitive Development (ABCD) study and the Human Connectome Project (HCP). To derive a reference for future study design, we also considered the Transdiagnostic Connectome Project (TCP), Major Depressive Disorder (MDD), Alzheimer’s Disease Neuroimaging Initiative (ADNI) and the Singapore geriatric intervention study to reduce cognitive decline and physical frailty (SINGER) datasets (Table 1). We find that to increase prediction power, longer scans and larger sample size can yield substantial cost savings compared with only increasing sample size.

**Table 1.** | Sample size and amount of scan time per participant in each dataset.

| Dataset | Sample Size (N) | Scan Time (T) | Remarks |
| --- | --- | --- | --- |
| HCP-REST | 792 | 57m 36s | Two sessions; two runs per session |
| ABCD-REST | 2565 | 20m | Four fMRI runs in a single session |
| SINGER-REST | 642 | 9m 56s | One fMRI run in a single session |
| TCP-REST | 194 | 26m 2s | Four fMRI runs in a single session |
| MDD-REST | 287 | 23m 12s | Four fMRI runs in a single session |
| ADNI-REST | 586 | 9m | One fMRI run in a single session |
| ABCD-MID | 2262 | 10m 44s | Two fMRI runs in a single session |
| ABCD-NBACK | 2262 | 9m 39s | Two fMRI runs in a single session |
| ABCD-SST | 2262 | 12m 39s | Two fMRI runs in a single session |

## Results

### Larger sample size can compensate for shorter scan time & vice versa

For each participant in the HCP and ABCD datasets, we calculated a 419 × 419 resting-state functional connectivity (RSFC) matrix using the first *T* minutes of fMRI (Fischl, 2012; Schaefer et al., 2018). *T* was varied from 2 minutes to the maximum scan time in each dataset in intervals of 2 minutes. The RSFC matrices (from the first *T* minutes) served as input features to predict a range of phenotypes in each dataset using kernel ridge regression (KRR) via a nested inner-loop cross-validation procedure. The analyses were repeated with different numbers of training participants (i.e., different training sample size *N*). Within each cross-validation loop, test participants were fixed across different training set sizes, so that prediction accuracy was comparable across different training set sizes (Extended Data Fig. 1). The whole procedure was repeated multiple times and averaged. Sample size and maximum scan time of all datasets are found in Table 1.

We first considered the cognitive factor score because the cognitive factor score was predicted the best across all phenotypes (Ooi et al., 2022). Fig. 1a shows the prediction accuracy (Pearson’s correlation) of the ABCD cognitive factor score. Along a black iso-contour line, the prediction accuracy is (almost) constant even though scan time and sample size are changing. Consistent with previous literature (He et al., 2020; Schulz et al., 2023), increasing the number of training participants (when scan time per participant is fixed) improved prediction performance. Similarly, increasing scan time per participant (when sample size is fixed) also improved prediction performance (Feng et al., 2023).

**Fig. 1.**
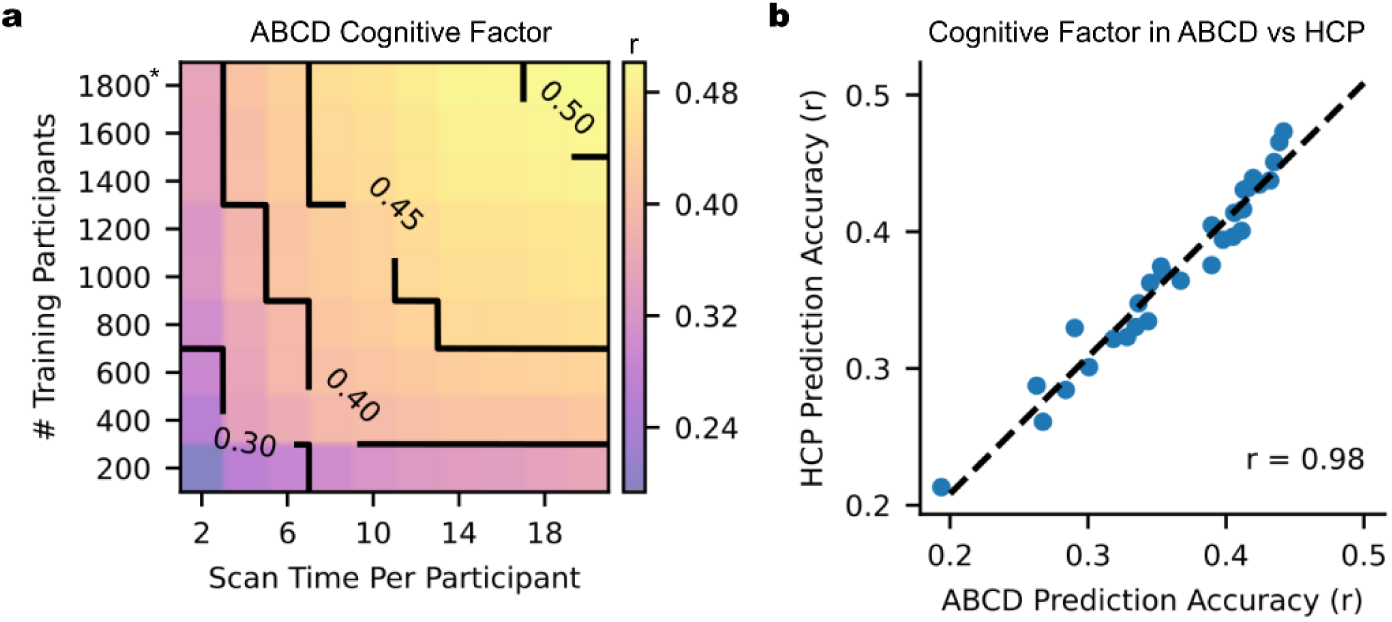
| Increasing training participants and scan time per participant lead to higher prediction accuracy of phenotypes. **a**. Contour plot of prediction accuracy (Pearson’s correlation) of the cognitive factor score as a function of the scan time *T* used to generate the functional connectivity matrix, and the number of training participants *N* used to train the predictive model in the Adolescent Brain and Cognitive Development (ABCD) dataset. Increasing training participants and scan time both improved prediction performance. The * indicates that all available participants were used, therefore the sample size will be close to, but not exactly the number shown. **b**. Scatterplot of the cognitive factor score prediction accuracy in the ABCD (x-axis) and HCP (y-axis) datasets. Each dot represents the prediction accuracy in each dataset for a particular pair of sample size and scan time per participant.

Although cognitive factor scores are not necessarily comparable across datasets (due to population and phenotypic differences), prediction accuracies were highly similar between the ABCD and HCP datasets (Fig. 1b & Supplementary Fig. 1). Similar conclusions were also obtained when we measured prediction accuracy using coefficient of determination (COD) instead of Pearson’s correlation (Supplementary Fig. 2), computed RSFC using the first *T* minutes of uncensored data (Supplementary Fig. 3), did not perform censoring of high motion frames (Supplementary Fig. 4), or utilized linear ridge regression (LRR) instead of KRR (Supplementary Figs. 5 & 6).

In the ABCD study (where maximum scan time was 20 minutes), Fig. 2a shows that the prediction accuracy of the cognitive factor increases with total scan duration (# training participants × scan time per participant), suggesting that sample size and scan time per participant are broadly interchangeable. In the HCP dataset, we observed diminishing returns of scan time relative to sample size for scan time beyond 30 minutes (Fig. 2a).

**Fig. 2.**
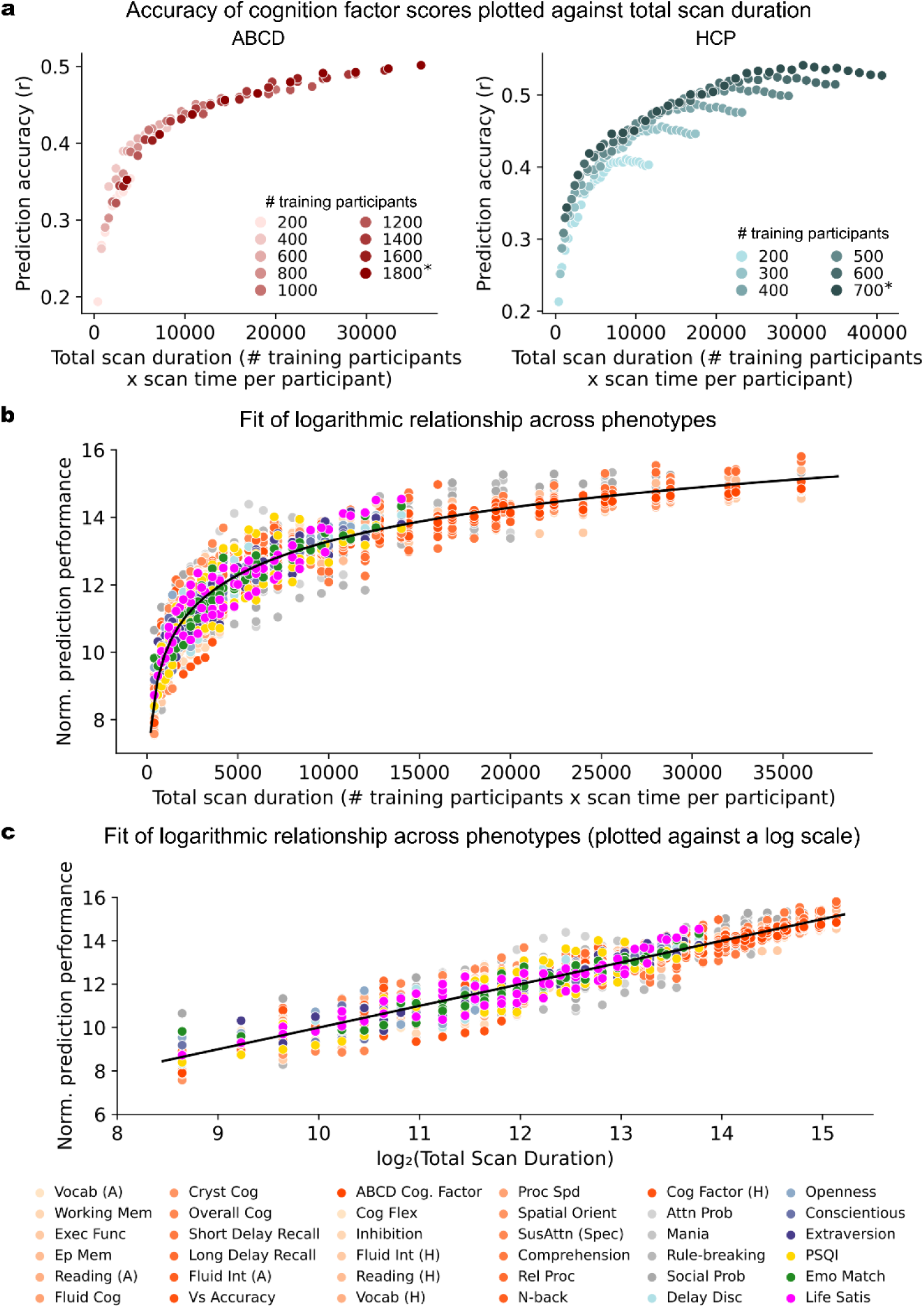
| Relationship between prediction accuracy and total scan duration (sample size × scan time per participant). a. Scatter plot showing prediction accuracy (Pearson’s correlation) of the cognitive factor as a function of total scan duration (defined as # training participants × scan time per participant). Each color shade represents different number of total participants used to train the prediction algorithm. Plots are repeated for the Adolescent Brain and Cognitive Development (ABCD) study and Human Connectome Project (HCP). The * indicates that all available participants were used, therefore the sample size will be close to, but not exactly the number shown. There was a diminishing return of scan time per participant (relative to sample size) beyond 30 minutes in the HCP dataset. In the ABCD dataset, where maximum scan time per participant was 20 minutes, the diminishing returns of scan time (relative to sample size) was not observed. b. Scatter plot showing normalized prediction accuracy of the cognitive factor scores and 34 other phenotypes versus total scan duration ignoring data beyond 20 minutes of scan time. Cognitive, mental health, personality, physicality, emotional and well-being measures are shown in shades of red, grey, blue, yellow, green and pink, respectively. The logarithmic black curve suggests that total scan duration explained prediction performance well across phenotypic domains and datasets. c. Same as panel (b), except the horizontal axis (total scan duration) is plotted on a logarithm scale. The linear black line suggests that the logarithm of total scan duration explained prediction performance well across phenotypic domains and datasets.

Beyond the cognitive factor scores, we focused on 28 (out of 59) HCP phenotypes and 23 (out of 37) ABCD phenotypes with maximum prediction accuracies of r > 0.1 (Supplementary Table 1a). We found that 89% (i.e., 25 out of 28) HCP phenotypes exhibited diminishing returns of scan time (relative to sample size) beyond 20 to 30 minutes. Diminishing returns were not observed for any of the 23 ABCD phenotypes.

A logarithmic pattern was evident in 76% (19 out of 25) HCP and 74% (17 out of 23) ABCD phenotypes (Supplementary Table 1a; Supplementary Figs. 7 & 8). To quantify the logarithmic relationship, for each of the 19 HCP and 17 ABCD phenotypes, we fitted a logarithm curve (with two free parameters) between prediction accuracy and total scan duration (ignoring data beyond 20 minutes per participant). The logarithm fit allowed phenotypic measures from both datasets to be plotted on the same normalized prediction performance scale (Figs. 2b and 2c).

The black curves (Figs. 2b and 2c) depict the logarithmic fit of the phenotypes (dots in Figs. 2b and 2c). Overall, total scan duration explained prediction accuracy across HCP and ABCD phenotypes remarkably well: coefficient of determination (COD) or R^2^ = 0.88 and 0.89 respectively.

The logarithm curve was also able to explain prediction accuracy well across different prediction algorithms (KRR and LRR) and different performance metrics (COD and r), as illustrated for the cognitive factor scores in Supplementary Fig. 9.

### Theoretical model explains diminishing returns of scan time relative to sample size

We have observed diminishing returns of scan time relative to sample size. To examine this phenomenon more closely, we considered the prediction accuracy of the HCP factor score across six combinations of sample size and scan time totalling 6000 mins of total scan duration (Fig. 3a). Prediction accuracy decreased with increasing scan time per participant, despite maintaining 6000 minutes of total scan duration (Fig. 3a). However, the accuracy reduction was modest below 30 minutes of scan time. Similar conclusions were obtained for all 19 HCP and 17 ABCD phenotypes that followed a logarithmic fit (Supplementary Fig. 10). These results indicate that while longer scan times can offset smaller sample sizes, the required increase in scan time becomes progressively larger as scan duration extends.

**Fig. 3.**
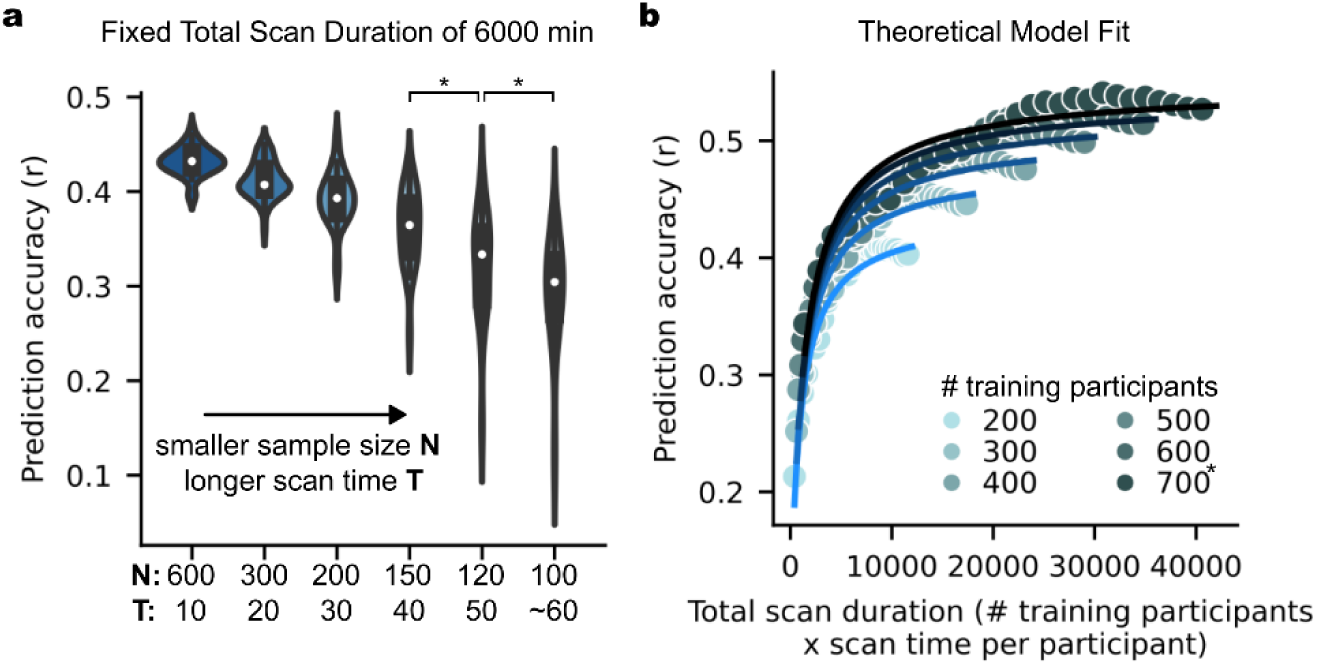
| As scan time increases, sample size eventually becomes more important than scan time. **a.** Prediction accuracy of the HCP cognition factor score when total scan duration is fixed at 6000 minutes, while varying scan time per participant from 10 to 60 minutes. Each violin plot shows the distribution of prediction accuracies across 50 random cross-validation splits. * indicates that the distributions of prediction accuracies were significantly different between adjacent pairs of sample sizes (and scan time per participant) after false discovery rate (FDR) q < 0.05 correction. *N* refers to sample size, while *T* refers to scan time per participant. **b.** Scatter plot of prediction accuracy against total scan duration for the cognitive factor score in the HCP dataset. The curves were obtained by fitting a theoretical model to the prediction accuracies of the cognitive factor score. The theoretical model explains why sample size is more important than scan time (see main text).

To gain insights into this phenomenon, we derived a closed-form mathematical relationship relating prediction accuracy (Pearson’s correlation) with scan time per participant *T* and sample size *N* (see Methods). To provide an intuition for the theoretical derivations, we note that phenotypic prediction can be theoretically decomposed into two components: one component relating to an average prediction (common to all participants) and a second component relating to a participant’s deviation from this average prediction.

The uncertainty (variance) of the first component scales as 1/*N*, like a conventional standard error of the mean. For the second component, we note that the prediction can be written as regressions coefficients × FC (for linear regression). The uncertainty (variance) of the regression coefficient estimates scales with 1/*N*. The uncertainty (variance) of the FC estimates scales with 1/*T* (i.e. reliability improves with *T*). Thus, the uncertainty of the second component scales with 1/*NT*. Overall, our theoretical derivation suggests that prediction accuracy can be expressed as a function of 1/*N* and 1/*NT* with three free parameters.

The theoretical derivations do not tell us the relative importance of the 1/*N* and 1/*NT* terms. Therefore, we fitted the theoretical model to actual prediction accuracies in the HCP and ABCD datasets. The goal was to determine (1) whether our theoretical model (despite the simplifying assumptions) would still explain the empirical results, and (2) to determine the relative importance of 1/N and 1/NT.

We found an excellent fit with actual prediction accuracies for the 19 HCP and 17 ABCD phenotypes (Figs. 3b, Supplementary Figs. 11 & 12): R^2^ = 0.89 for both datasets (Supplementary Table 1b). When *T* was small, the 1/*NT* term dominated the 1/*N* term, which explained the almost 1-to-1 interchangeability between scan time and sample size for shorter scan time. The existence of the 1/*N* term ensured that sample size was still slightly more important than scan time even for small *T*. FC reliability eventually saturated with increasing *T*. Therefore, the 1/*N* term eventually dominated the 1/*NT* term, so sample size became much more important than scan time.

For 20-min scans, the logarithmic and theoretical models performed equally well with equivalent goodness of fit (R^2^) across the 17 ABCD phenotypes (p = 0.57). For longer scan time, the theoretical model exhibited better fit than the logarithmic model (p = 0.002 across the 19 HCP phenotypes; Supplementary Fig. 13). Furthermore, prediction accuracy under the logarithmic model will exceed a correlation of one for sufficiently large N and T, which is untenable. Therefore, we will use the theoretical model in the remaining portions of the study.

### Theoretical model works better for well-predicted phenotypes

Recall that the 17 ABCD phenotypes and 19 HCP phenotypes were predicted with maximum prediction accuracies of Pearson’s r > 0.1, and the theoretical model was able to explain their prediction accuracies with an explained variance of 89% (Supplementary Table 1b). If we loosened the prediction threshold to include phenotypes whose prediction accuracies (Pearson’s r) were positive in at least 90% of all combinations of sample size *N* and scan time *T* (Supplementary Table 1a), the model fit was lower but still relatively high with average explained variance 76% and 73% in ABCD and HCP datasets respectively (Supplementary Table 1b).

More generally, phenotypes with high overall prediction accuracies adhered to the theoretical model well (example in Fig. 4a), while phenotypes with poor prediction accuracies resulted in poor adherence to the model (example in Fig. 4b). Indeed, model fit was strongly correlated with prediction accuracy across phenotypes in both datasets (Figs. 4c to 4d). These findings suggest that the imperfect fit of the theoretical model for some phenotypes may be due to their poor predictability, rather than true variation in prediction accuracy with respect to sample size and scan time.

**Fig. 4.**
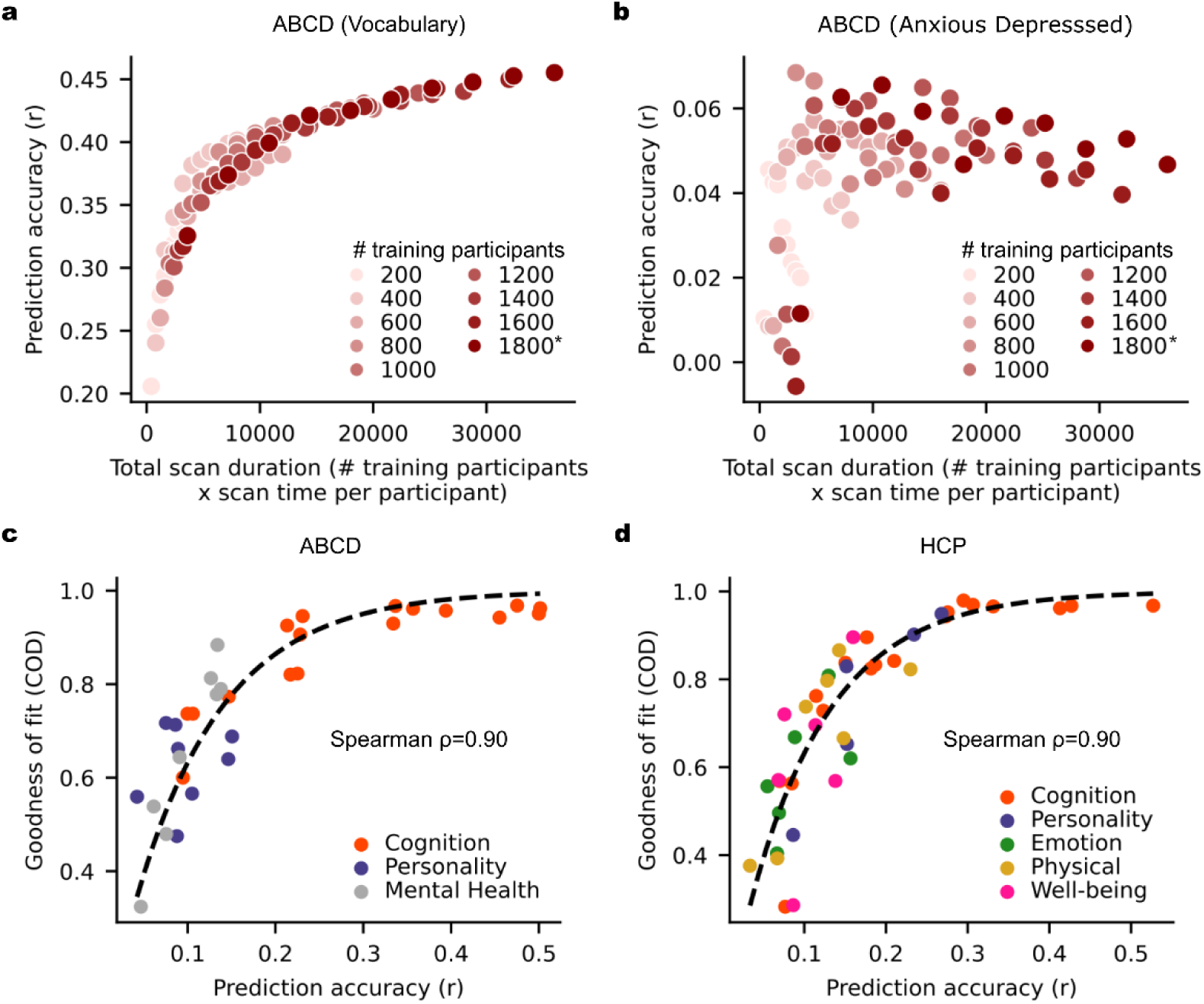
| Theoretical model works better for well-predicted phenotypes. **a.** Scatter plot of prediction accuracy against total scan duration for an exemplary phenotype with high prediction accuracy. **b.** Scatter plot of prediction accuracy against total scan duration for an exemplary phenotype with low prediction accuracy. **c.** Scatter plot of theoretical model goodness-of-fit (coefficient of determination or COD) against prediction accuracies of different ABCD phenotypes. COD (also known as R^2^) is a measure of explained variance. Here, we considered phenotypes whose prediction accuracies (Pearson’s r) were positive in at least 90% of all combinations of sample size *N* and scan time *T*, yielding 42 HCP phenotypes and 33 ABCD phenotypes. Prediction accuracy (horizontal axis) was based on maximum scan time and sample size. For visualization, we plot a dashed black line by fitting to a monotonically increasing function. **d.** Same as panel (c) but using HCP (instead of ABCD) dataset.

### Non-stationarity in fMRI-phenotype relationships weakens model adherence

As noted above, some phenotypes likely fail to match the theoretical model because of intrinsically poor predictability. However, there were also phenotypes that were reasonably well-predicted yet still exhibited a poor fit to the theoretical model. For example, “Anger: Aggression” was reasonably well-predicted in the HCP dataset, but prediction accuracy was primarily improved by sample size and not scan time (Fig. 5a). As scan time per participant increased, prediction accuracy appeared to increase, decrease and then increase again. This pattern was remarkably consistent across sample sizes (Fig. 5a).

**Fig. 5.**
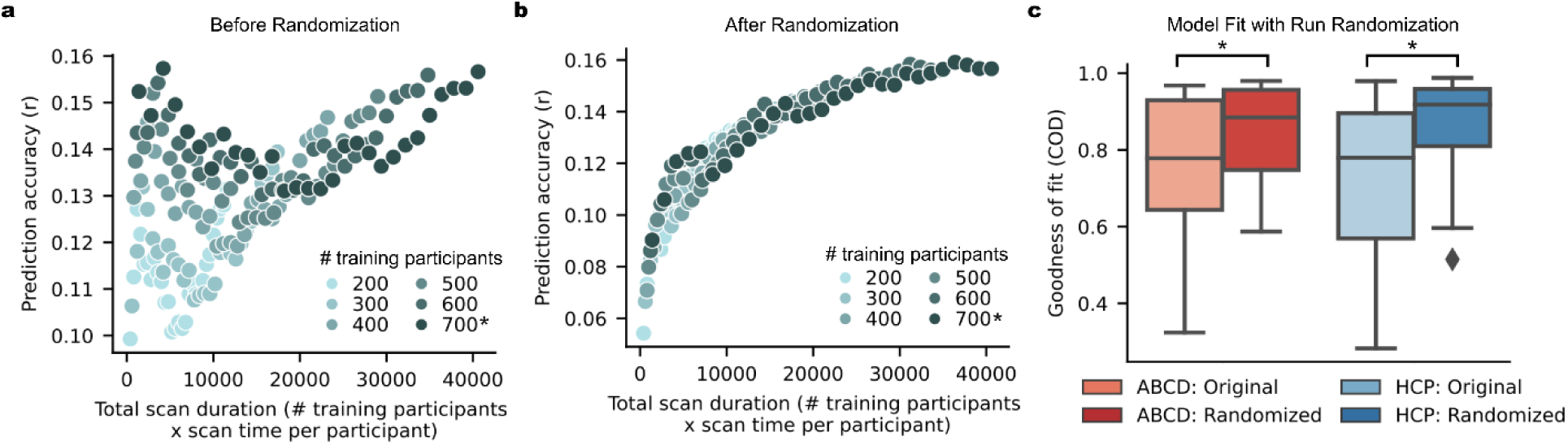
| Non-stationarity in fMRI-phenotype relationship weakens adherence to theoretical model. **a.** Scatter plot of prediction accuracy against total scan duration for the “Anger: Aggression” phenotype in the HCP dataset. Despite relatively high accuracy, the phenotype improved with larger sample size, but not scan time. As scan time per participant increases, prediction accuracy appeared to increase, decrease, then increase again. **b.** Scatter plot of prediction accuracy against total scan duration for the “Anger: Aggression” phenotype in the HCP dataset after randomizing fMRI run order for each participant. Observe that the prediction accuracy now adheres strongly to the theoretical model. **c.** Box plots showing goodness of fit to theoretical model before and after randomizing fMRI run order for 33 ABCD and 42 HCP phenotypes. * indicates that goodness-of-fits were significantly different (after FDR correction with q < 0.05). Here, we considered all phenotypes whose prediction accuracies (Pearson’s r) were positive in at least 90% of all combinations of *N* and *T*. For each boxplot, the horizontal line indicates the median across 33 ABCD phenotypes or 42 HCP phenotypes. The bottom and top edges of the box indicate the 25th and 75th percentiles, respectively. Outliers are defined as data points beyond 1.5 times the interquartile range. The whiskers extend to the most extreme data points not considered outliers.

This suggests that fMRI-phenotype relationships might be non-stationary for certain phenotypes, which violates an assumption in the theoretical model. To put this in more colloquial terms, the assumption is that the FC-phenotype relationship is the same (i.e., stationary) regardless of whether FC was computed based on five minutes of fMRI from the beginning, middle or end of the MRI session. We note that for both HCP and ABCD datasets, the fMRI data was collected over four runs. To test for non-stationarity, we randomized the fMRI run order independently for each participant and repeated the FC computation (and prediction) using the “first” T minutes of resting-state fMRI data under the randomized run order. The run randomization improved the goodness of fit of the theoretical model, suggesting the presence of non-stationarities (Figs. 5b and 5c).

Arousal changes between or during resting-state scans are well-established (Tagliazucchi et al., 2014; Wang et al., 2016; Bijsterbosch et al., 2017; Laumann et al., 2017; Orban et al., 2020), therefore we expect fMRI scans, especially longer duration scans, to be non-stationary. However, since run randomization affected some phenotypes more than others, this suggests that there is an interaction between fMRI non-stationarity and phenotypes, i.e., the fMRI-phenotype relationship is also non-stationary.

### Overhead cost per participant increases importance of scan time relative to sample size

We have shown that investigators have some flexibility of attaining a specified prediction accuracy through different combinations of sample size and scan time per participant (Fig. 1). Furthermore, the theoretical model suggests that sample size is more important than scan time (Fig. 3). However, for the purpose of study design, it is important to consider the fundamental asymmetry between sample size and scan time per participant because of inherent overhead cost associated with each participant. These overhead costs might include recruitment effort, manpower to perform neuropsychological tests, additional MRI modalities (e.g., anatomical T1, diffusion MRI), other biomarkers (e.g., position emission tomography or blood tests). Therefore, the overhead cost can often be higher than the cost of fMRI itself.

To derive a reference for future studies, we considered four additional resting-state datasets (TCP, MDD, ADNI and SINGER). 34 phenotypes exhibited good fit to the theoretical model (Supplementary Table 2; Supplementary Figs. 14 to 17). We also considered task-FC of the three ABCD tasks, and found that the number of phenotypes with good fit to the theoretical model ranged from 16 to 19 (Supplementary Table 2; Supplementary Figs. 18 to 20).

In total, we considered nine datasets: six resting-fMRI datasets and three ABCD task-fMRI datasets. We fitted the theoretical model to 76 phenotypes in the nine datasets, yielding 89% average explained variance (Supplementary Table 2). These datasets span multiple fMRI sequences (single-echo single-band, single-echo multi-band, multi-echo multi-band), coordinate systems (fsLR, fsaverage, MNI152), racial groups (Western and Asian populations), mental disorders (healthy, neurological and psychiatric) and age groups (children, young adults and elderly). More dataset characteristics are found in Supplementary Table 4.

For each phenotype, the fitted model was normalized by the phenotype’s maximum achievable accuracy (estimated by the theoretical model), yielding a fraction of maximum achievable prediction accuracy for every combination of sample size and scan time per participant. The fraction of maximum achievable prediction accuracy was then averaged across the phenotypes under a hypothetical 10-fold cross-validation scenario (Fig. 6a). We note that the correlations between Fig. 6a and Fig. 1 across corresponding sample sizes and scan durations were 0.99 for both HCP and ABCD datasets.

**Fig. 6.**
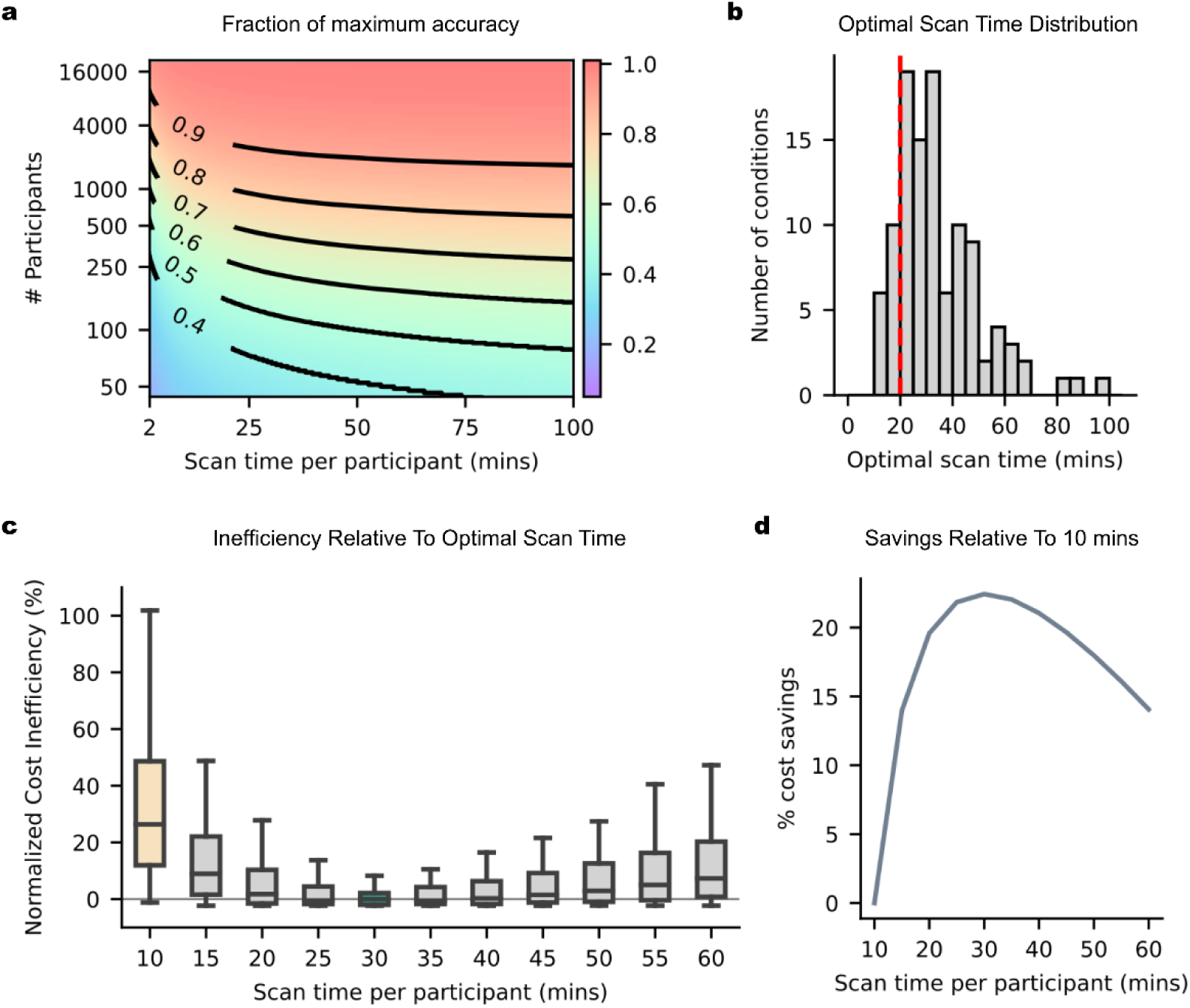
| 30-min scans yield significant cost savings over 10-min scans across nine datasets. **a.** Fraction of maximum prediction accuracy as a function of sample size and scan time per participant averaged across 76 phenotypes from nine datasets (six resting-fMRI datasets and three ABCD task-fMRI datasets). Here sample size was calculated assuming 90% of participants were used for training the model, and 10% of participants were used for model evaluation (i.e., 10-fold cross-validation). **b.** Histogram of optimal scan time (with regards to costs) across 108 scenarios. Given 3 possible accuracy targets (80%, 90% or 95% of maximum achievable accuracy), 2 possible overhead costs ($500 or $1000 per participant) and 2 possible scan costs per hour ($500 or $1000), there are 3 × 2 × 2 = 12 conditions. In total, we have 9 datasets × 12 conditions = 108 scenarios. For 85% of the scenarios, the optimal scan time (in terms of cost) was at least 20 min (red dashed line). **c.** Normalized cost inefficiency (across the 108 scenarios) as a function of a fixed scan time per participant, relative to the optimal scan time in (b). We note that in practice, the optimal scan time in (b) is not known in advance, so this plot seeks to derive a **fixed** optimal scan time generalizable to most situations. Each boxplot contains 108 data points (corresponding to the 108 scenarios). For visualization, the boxplots are normalized by subtracting the cost inefficiency of the best possible fixed scan time (30 min in this case), so that the normalized cost inefficiency of the best possible fixed scan time is centered at zero. **d.** Cost savings relative to 10 min of scan time per participant. The greatest cost saving (22%) was achieved at 30 min.

Given a scan cost per hour (e.g., $500) and overhead cost per participant ($500), we can find all pairs of sample size and scan time that fit within a particular fMRI budget (e.g., $1M). We can then use Fig. 6a to find the optimal sample size and scan time leading to the largest fraction of maximum prediction accuracy. Extended Data Fig. 2 illustrates the prediction accuracy achievable with different fMRI budgets, costs per hour of scan time and overhead cost per participant. Extended Data Table 1 shows the optimal scan time for a wider range of fMRI budgets, scan cost per hour and overhead cost per participant.

Obviously larger fMRI budgets, lower scan cost and lower overhead cost enable larger sample size and scan time, leading to greater achievable prediction accuracy (Extended Data Fig. 2). From the curves, we can determine the optimal scan time to achieve the greatest prediction accuracy within a fixed scan budget (solid circles in Extended Data Fig. 2). Optimal scan time increases with larger overhead cost, lower fMRI budget and lower scan cost. All curves rise steeply with increasing scan time per participant, and then declines slowly with increasing scan time. The asymmetry of the curves suggest that it is better to overshoot than undershoot optimal scan time.

As an example, a United States National Institutes of Health R01 grant is worth $2.5M. Assuming an fMRI budget of $1M, a scan cost of $500 per hour and an overhead cost of $500 per participant, the optimal scan time would be 34.5 min per participant. Suppose positron emission tomography was also collected, then the overhead cost might shoot up to $5000 per participant, resulting in an optimal scan time of 159.3 min per participant.

### 30-min scans are considerably cheaper than 10-min scans

Beyond optimizing scan time to maximize prediction accuracy within a fixed scan budget (previous section), the model fits shown in Fig. 6a can also be used to optimize scan time to minimize study cost to achieve a fixed accuracy target. For example, suppose we want to achieve 90% of maximum achievable accuracy, we can find all pairs of sample size and scan time per participant along the black contour line corresponding to 0.9 in Fig. 6a. For every pair of sample size and scan time, we can then compute the study cost given a particular scan cost per hour (e.g., $500) and a particular overhead cost per participant (e.g., $1000). The optimal scan time (and sample size) with the lowest study cost can then be obtained.

Here, we considered 3 possible accuracy targets (80%, 90% or 95% of maximum accuracy), 2 possible overhead costs ($500 or $1000 per participant) and 2 possible scan costs per hour ($500 or $1000). In total there were 3 × 2 × 2 = 12 conditions. Since there were nine datasets, this resulted in 12 × 9 = 108 scenarios. In the vast majority (85%) of these 108 scenarios, the optimal scan time was at least ≥20 min (Fig. 6b).

However, during study design, the optimal scan time (Fig. 6b) is not known in advance, therefore we also seek to derive a *fixed* scan time that is the most cost-effective in most situations. Fig. 6c shows the normalized cost inefficiency of various fixed scan times relative to the optimal scan time for each of 108 scenarios. Many consortia BWAS collect 10-min fMRI scans, which is highly cost inefficient. On average across resting and task states, 30-min scans were the most cost-effective, yielding 22% cost savings over 10-min scans (Fig. 6d). We again note the asymmetry in the cost curves, so it is cheaper to overshoot than undershoot the most cost-effective scan time.

### Most cost-effective scan time for task-fMRI is shorter than for resting-fMRI

Across the six resting-state datasets (Fig. 7a), the most cost-effective scan time was the longest for ABCD (60 min) and shortest for the TCP and ADNI datasets (20 min). However, a scan time of 30 minutes was still relatively cost-effective for all datasets, because of a flat cost curve near the optimum and the asymmetry of the cost curve. For example, even for the TCP dataset, which had the shortest most cost-effective scan time of 20 min, over-scanning with 30-min scans led to only 3.7% higher cost relative to 20 min, compared with 7.3% higher cost for under-scanning with 10-min scans.

**Fig. 7.**
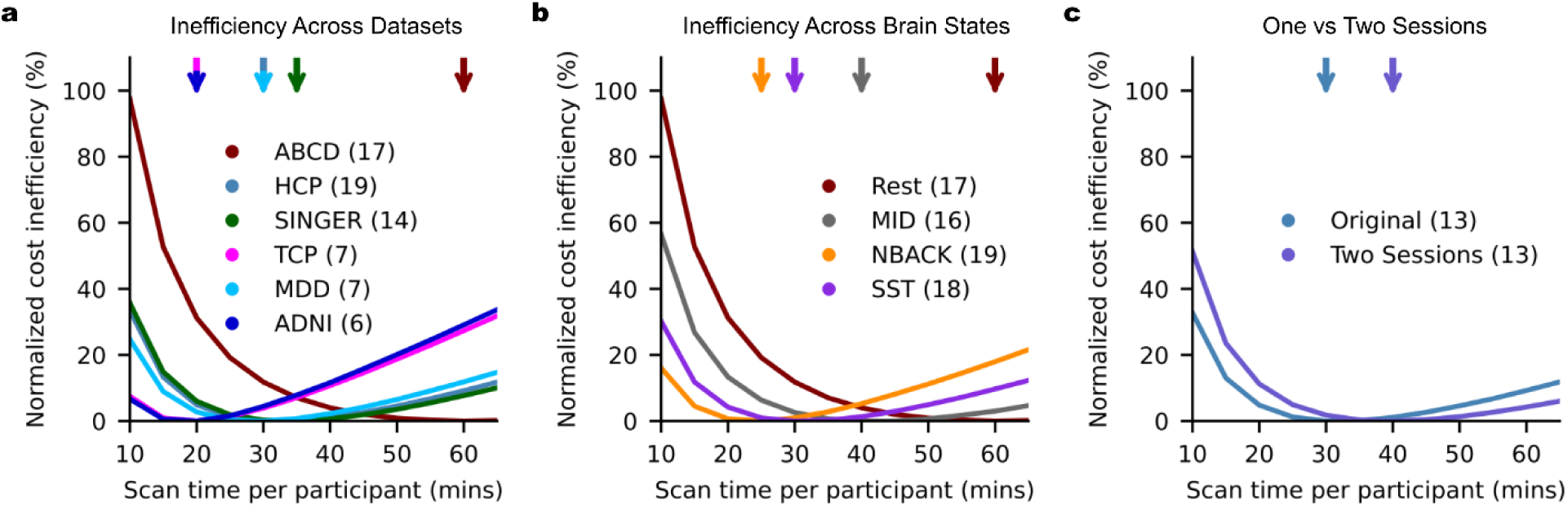
| Variation in the most cost-effective scan time across resting-state and task-state fMRI. **a.** Cost inefficiency as a function of scan time per participant for the six resting-state datasets. This plot provides the same information as Fig. 6c but shown for each dataset separately. Numbers in brackets indicate number of phenotypes. **b.** Cost inefficiency for ABCD resting-state and task-state fMRI. **c.** Cost inefficiency if data was collected in two separate sessions (based on the HCP dataset). Similar to Fig. 6c, for visualization, each curve is normalized by subtracting the cost inefficiency of the best possible fixed scan time (of each curve), so that the normalized cost inefficiency of the best possible fixed scan time is centered at zero.

Previous studies have shown that task-FC leads to better prediction performance for cognitive measures (Greene et al., 2018; J. Chen et al., 2022). Here, we extend previous results, finding that the most cost-effective scan time was significantly shorter in ABCD task-fMRI than ABCD resting-fMRI (Fig. 7b). Among the three tasks, the most cost-effective scan time was the shortest for the n-back task (25 min). However, even for the n-back task, 30-min scans led to only 0.9% higher cost (relative to 25 min), compared with 16.1% higher cost for 10-min scans.

These task results suggest that the most cost-effective scan time is sensitive to brain state manipulation. Task-based fMRI may preferentially engage cognitive and physiological mechanisms that are closely tied to the expression of specific phenotypes (e.g., processing speed), thereby enhancing the specificity of functional connectivity estimates for phenotypic prediction. Tasks may also facilitate shorter, more efficient scan durations by aligning brain states across individuals in a controlled manner, thereby reducing spurious non-stationary influences that could otherwise obscure reliable modeling of inter-individual differences. This alignment might be better achieved in tasks that present stimuli and conditions with identical timing across participants – whether using event-related or block designs.

Non-stationarity may also be potentially increased by distributing resting state fMRI runs across multiple sessions. Since the HCP dataset was collected on two different days (sessions), we were also able to directly compare the effect of a two-session versus a one-session design, controlling for total scan time. The most cost-effective scan time for the two-session design was only slightly longer than the original HCP analysis: 40 min vs 30 min (Fig. 7c).

Overall, these results suggest that state manipulation can influence the most cost-effective scan time, and that a relatively large state manipulation (e.g., task) can significantly influence cost-effectiveness.

### Variation across phenotypic domains and scan parameters

There were clear variations across phenotypes. For example, there were phenotypes that could be predicted very well and demonstrated prediction gains up to the maximum amount of data per participant (e.g., age in the ADNI dataset; Supplementary Fig. 17). However, there were also other phenotypes, which were predicted less well (e.g., BMI in the SINGER dataset; Supplementary Fig. 14), but showed prediction gains up to the maximum amount of data per participant. Since single phenotypes are not easily interpreted, we grouped the phenotypes into seven phenotypic domains to study phenotypic variation in more details.

For five of the seven phenotypic domains, the most cost-effective scan times ranged from 25 min to 40 min (Extended Data Fig. 3a). The most cost-effective scan time for emotion domain was very long, but this outlier was driven by a single phenotypic measure, so should not be over-interpreted. For the positron emission tomography (PET) phenotypic domain, our original scenarios assumed overhead costs of $500 or $1000 per participant, which was unrealistic. Assuming a more realistic overhead PET cost per participant ($5000 or $10,000) yielded 50 min as the most cost-effective scan time.

Although there was a strong relationship between phenotypic prediction accuracy and goodness-of-fit to the theoretical model (Fig. 4), we did not find an obvious relationship between phenotypic prediction accuracy and optimal scan time (Extended Data Fig. 3b). Recent studies have also demonstrated that phenotypic reliability is important for BWAS power (Nikolaidis et al., 2022; Gell et al., 2023). In our theoretical model, phenotypic reliability directly impacts overall prediction accuracy but does not directly contribute to the trade-off between sample size and scan time. Indeed, there was not an obvious relationship between phenotypic test-retest reliability and optimal scan time (Extended Data Fig. 3c).

There was also not an obvious relationship of optimal scan time with temporal resolution, voxel resolution or scan sequence (Extended Data Figs. 3d to 3f). We emphasize that we are not claiming that scan parameters do not matter, but that other variations between datasets (e.g., phenotypes, populations) might exert a greater impact than common variation in scan parameters.

Consistent with the previous sections, we note that for the vast majority of phenotypes and scan parameters, the optimal scan time was at least 20 min, and on average the most cost-effective scan time was 30 min (Fig. 6c).

### Most cost-effective scan time for subcortical BWAS is longer than whole-brain BWAS

Our main analyses involved a cortical parcellation with 400 regions and 19 subcortical regions, yielding 419 × 419 RSFC matrices. We also repeated the analyses using 19 × 419 subcortical-cortical RSFC matrices. The most cost-effective scan time for subcortical-cortical RSFC was about double that of whole-brain RSFC (Fig. 8a). This might arise because of lower signal-to-noise ratio (SNR) in subcortical regions, resulting in the need for longer scan time to achieve better estimate of subcortical FC.

**Fig. 8.**
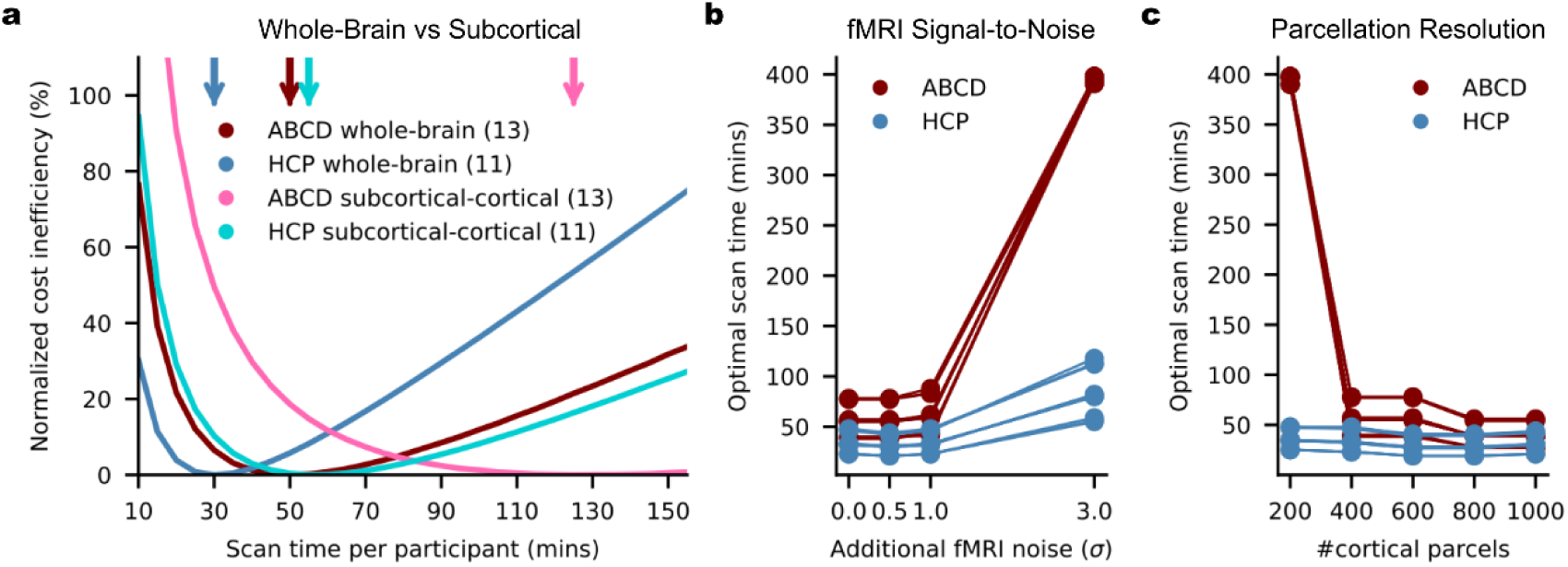
| Most cost-effective scan time for subcortical BWAS is longer than whole-brain BWAS. **a.** Cost inefficiency as a function of scan time per participant with subcortical-cortical FC versus whole-brain FC. For visualization, similar to Fig. 6c, the curves are normalized by subtracting the cost inefficiency of the best possible fixed scan time (of each curve), so that the normalized cost inefficiency of the best possible fixed scan time is centered at zero. Numbers in brackets indicate number of phenotypes in each condition. **b.** Optimal scan time for predicting the cognitive factor score as a function of simulated Gaussian noise with standard deviation σ. **c.** Optimal scan time for predicting the cognitive factor score as a function of the resolution of cortical parcellation.

To explore the effects of fMRI SNR, for each parcel time course, we z-normalized the fMRI time course, so the resulting standard deviation of the time course was equal to one. We then added zero mean Gaussian noise with standard deviation σ. Even doubling the noise (σ = 1) had very little impact on the optimal scan time (Fig. 8b). As a sanity check, we added a large quantity of noise (σ = 3), which lead to a much longer optimal scan time. (Fig. 8b).

Intuitively, this is not surprising because lower SNR means that a longer scan time is necessary to get an accurate estimate of individual-level FC. What is interesting is that a large SNR change is necessary to make a noticeable difference in optimal scan time, which might explain the robustness of optimal scan times across the common scan parameters we explored in the previous section (Extended Data Fig. 3). Therefore, even with small to moderate technological improvements in SNR, the most cost-effective scan time is unlikely to deviate from our estimate. Major increase in SNR could however shorten the most cost-effective scan time from the current estimates.

We also varied the resolution of the cortical parcellation with 200, 400, 600, 800 or 1000 parcels for predicting the cognitive factor scores in the HCP and ABCD datasets. There was a weak trend in which higher parcellation resolution led to slightly lower optimal scan time, although there was a big drop in optimal scan time from 200 parcels to 400 parcels in the ABCD dataset (Fig. 8c). Given that subcortical-cortical FC have less edges (features) than whole-brain FC, this could be another reason why subcortical-cortical FC requires longer optimal scan time.

### Accuracy versus reliability

Finally, we turn our attention to the effects of sample size and scan time per participant on the reliability of BWAS (Marek et al., 2022) using a previously established split-half procedure (Supplementary Fig. 21a; Tian et al., 2021; Chen et al., 2023). For both univariate and multivariate BWAS reliability, diminishing returns of scan time (relative to sample size) occurred beyond 10 mins per participant (Supplementary Figs. 22 to 30), instead of 20 mins for prediction accuracy (Fig. 2). We note that reliability is necessary, but not sufficient for validity (Schmidt et al., 2000; Noble et al., 2019). For example, hardware artifacts may appear reliably in measurements without having any biological relevance. Thus, reliable BWAS features do not guarantee accurate prediction of phenotypic measures. As such, we recommend that researchers prioritize prediction accuracy during study design.

### Longer scans improve prediction accuracy while reducing costs

To summarize, 30-min scans are on average the most cost-effective across resting-state and task-state whole-brain BWAS (Fig. 6c). The cost curves are also asymmetric, so it is cheaper to overshoot than undershoot the optimum. Therefore, even when the most-effective scan time is shorter than 30 min (e.g., n-back task or TCP dataset), 30-min scans only incurred a small penalty, relative to clairvoyantly knowing the true optimal scan time in advance. Furthermore, for subcortical BWAS, the most cost-effective scans are much longer than 30 mins.

Our results present a compelling case for moving beyond traditional power analyses, whose only inputs are sample size, to inform BWAS design. Such power analyses can only point towards maximizing sample size, so scan time becomes implicitly minimized under budget constraints. Our findings show that we can achieve higher prediction performance by increasing both sample size and scan time, while generating substantial cost-savings compared to only increasing sample size alone.

Our results complement recent advocation for larger sample sizes to increase BWAS reproducibility (Marek et al., 2022). Consistent with previous studies (Varoquaux, 2018), when sample size is small, there is a high degree of variability across cross-validation folds (Fig. 3a). Furthermore, large sample sizes are still necessary for high prediction accuracy. To achieve 80% of maximum prediction accuracy with 30-min scans, a sample size of ∼900 is necessary (Fig. 6a), which is much larger than typical BWAS (Marek et al., 2022). To achieve 90% of maximum prediction accuracy with 30-min scans, a sample size of ∼2500 is necessary (Fig. 6a).

In addition to increasing sample size and scan time, BWAS effect sizes can also be enhanced through innovative study designs. Recent work showed that U-shaped population sampling can enhance the strength of associations between functional connectivity and phenotypic measures (Kang et al., 2024). However, more complex screening procedures will increase overhead cost per participant, which might lengthen the optimal scan time.

The current analysis was focused on high target accuracies (80%, 90% or 95%) and relatively low overhead costs ($500 or $1000). Lower target accuracies (in smaller-scale studies) and higher overhead costs (e.g., PET, multi-site data collection) will lead to longer cost-effective scan time (Extended Data Fig. 2). In practice, scans are also more likely to be spuriously shortened (e.g., due to participant discomfort) than to be spuriously extended. Therefore, we recommend scan time to be ≥30 min.

Overall, 10-min scans are rarely cost-effective, and optimum scan time is ≥20 mins in most BWAS (Fig. 6c). Among the datasets we analyzed, four included scans ≥20 mins, providing robust evidence to support this conclusion across multiple datasets. On the other hand, we could only identify one dataset (HCP) with scans exceeding 30 minutes and a sufficiently large sample size for inclusion in our study (Table 1). This limitation underscores the importance of our findings, emphasizing the need for BWAS to prioritize longer scans.

### Non-economic considerations

Beyond economic considerations, the representativeness of the data sample and the generalizability of predictive models to subpopulations are also important factors when designing a study (Benkarim et al., 2022; Greene et al., 2022; Li et al., 2022; Kopal et al., 2023; Dhamala et al., 2024; Gell et al., 2024). One approach would be to aim for a larger sample size (potentially at the expense of scan time) to ensure sufficient sample sizes for subpopulations.

Alternatively, one could also make the participant selection criteria more stringent to maintain representativeness of a subpopulation. However, this would drive up the recruitment cost for the subpopulation, so our results suggest that it might be economically more efficient to scan harder-to-recruit subpopulations for longer. For example, instead of 20-min resting-state scans for all ABCD participants, perhaps subpopulations (e.g., Black participants) could be scanned for a longer period of time.

In other situations, the sample size is out of the investigator’s control, e.g., if the investigator wants to scan an existing cohort. In the case of the SINGER dataset, the sample size was determined by the power calculation of the actual lifestyle intervention (Xu et al., 2022) with the imaging data included to gain further insights into the intervention. As another example, in large-scale prospective studies (e.g., UK Biobank), the sample size is determined by the fact that only a small proportion of participants will develop a given condition in the future (Allen et al., 2024). In these situations, the scan time becomes constrained by the overall budget and fitting all phenotyping efforts within a small number of sessions (to avoid participant fatigue). Nevertheless, even in these situations where sample size is predetermined, Fig. 6a can still provide an empirical reference on the marginal gains in prediction accuracy as a function of scan time.

Finally, some studies may necessitate extensive scan time per participant by virtue of the scientific question. For example, when studying sleep stages, it is not easy to predict how long a participant would need to enter a particular sleep stage. Conversely, some phenomena of interest might be inherently short-lived. For example, if the goal is to characterise the effects of a fast acting drug (e.g., nitrous oxide), then it might not make sense to collect long fMRI scans. Furthermore, not all studies are interested in cross-sectional relationships between brain and non-brain-imaging phenotypes. For example, in the case of personalized brain stimulation (Cash et al., 2021; Lynch et al., 2022) or neurosurgical planning (Boutet et al., 2021), significant quantity of resting-state fMRI data might be necessary for accurate individual-level network estimation (Laumann et al., 2015; Braga et al., 2017; Gordon et al., 2017).

### A web application for optimized study design

Beyond our broad recommendation of ≥30 mins of scan time, we recognize that investigators might be interested in achieving the best optimal sample size and scan time specific to their study’s constraints. Therefore, we built a web-application to help facilitate flexible study design (WEB_APPLICATION_LINK). The web application includes additional constraints not analyzed in the current study. For example, certain demographic and patient populations might not be able to tolerate longer scans, so an additional factor will be the maximum scan time in each MRI session. Furthermore, our analysis was performed on participants whose data survived quality control. Therefore, we have also provided an option on the web application to allow researchers to specify their estimate of the percentage of participants, whose data might be lost due to poor data quality or participant drop out. Overall, our empirically established guidelines provide actionable insights for significantly reducing costs, while improving BWAS individual-level prediction performance.

## Methods

### Datasets, phenotypes and participants

Following previous studies, we considered 58 HCP phenotypes (Kong et al., 2019; Li et al., 2019) and 36 ABCD phenotypes (Chen et al., 2022; J. Chen et al., 2023). We additionally consider a cognition factor score derived from all phenotypes from each dataset (Ooi et al., 2022), yielding a total of 59 HCP and 37 ABCD phenotypes (Supplementary Table 3).

In this study, we used resting-state fMRI from the HCP WU-Minn S1200 release. We filtered participants from Li’s set of 953 participants (Li et al., 2019), excluding participants who did not have at least 40 minutes of uncensored data (censoring criteria are discussed under “Image Processing”) and did not have the full set of the 59 non-brain-imaging phenotypes (henceforth referred to as phenotypes) that we investigated. This resulted in a final set of 792 participants with demographics found in Supplementary Table 4.

We additionally considered resting-state fMRI from the ABCD 2.0.1 release. We filtered participants from Chen’s set of 5260 participants (J. Chen et al., 2023). We excluded participants who did not have at least 15 minutes of uncensored resting-fMRI data (censoring criteria are discussed under “Image Processing”) and did not have the full set of the 37 phenotypes that we investigated. This resulted in a final set of 2565 participants with demographics found in Supplementary Table 4.

We also utilized resting-state fMRI from the SINGER baseline cohort. We filtered participants from an initial set of 759 participants, excluding participants who did not have at least 10 minutes of resting-fMRI data or did not have the full set of the 19 phenotypes that we investigated (Supplementary Table 3). This resulted in a final set of 642 participants with demographics found in Supplementary Table 4.

We utilized resting-state fMRI from the TCP dataset. We filtered participants from an initial set of 241 participants, excluding participants who did not have at least 26 minutes of resting-fMRI data or did not have the full set of the 19 phenotypes that we investigated (Supplementary Table 3). This resulted in a final set of 194 participants with demographics found in Supplementary Table 4.

We utilized resting-state fMRI from the MDD dataset. We filtered participants from an initial set of 306 participants. We excluded participants who did not have at least 23 minutes of resting-fMRI data or did not have the full set of the 20 phenotypes that we investigated (Supplementary Table 3). This resulted in a final set of 287 participants with demographics found in Supplementary Table 4.

We utilized resting-state fMRI from the ADNI datasets (ADNI 2, ADNI 3 and ADNI GO). We filtered participants from an initial set of 768 participants with both fMRI and PET scans acquired within 1 year of each other. We excluded participants who did not have at least 9 minutes of resting-fMRI data or did not have the full set of the 6 phenotypes that we investigated (Supplementary Table 3). This resulted in a final set of 586 participants with demographics found in Supplementary Table 4.

In addition, we considered task-fMRI from the ABCD 2.0.1 release. We filtered participants from Chen’s set of 5260 participants (J. Chen et al., 2023). We excluded participants who did not have all 3 task-fMRI data remaining after quality control, and did not have the full set of the 37 phenotypes that we investigated. This resulted in a final set of 2262 participants with demographics found in Supplementary Table 4.

### Image processing

For the HCP dataset, the MSMAll ICA-FIX resting state scans were used (Glasser et al., 2013). Global signal regression has been shown to improve behavioral prediction (Li et al., 2019), so we further applied global signal regression (GSR) and censoring, consistent with our previous studies (Li et al., 2019; He et al., 2020; Kong et al., 2021). The censoring process entailed flagging frames with either FD > 0.2mm or DVARS > 75. The frame immediately before and after flagged frames were marked as censored. Additionally, uncensored segments of data consisting of less than 5 frames were also censored during downstream processing.

For the ABCD dataset, the minimally processed resting state scans were utilized (Hagler et al., 2019). Processing of functional data was performed in line with our previous study (Chen et al., 2022). Specifically, we additionally processed the minimally processed data with the following steps. (1) The functional images were aligned to the T1 images using boundary-based registration (Greve et al., 2009). (2) Respiratory pseudomotion motion filtering was performed by applying a bandstop filter of 0.31-0.43Hz (Fair et al., 2020). (3) Frames with FD > 0.3mm or DVARS > 50 were flagged. The flagged frame, as well as the frame immediately before and two frames immediately after the marked frame were censored. Additionally, uncensored segments of data consisting of less than 5 frames were also censored. (4) Global, white matter and ventricular signals, 6 motion parameters, and their temporal derivatives were regressed from the functional data. Regression coefficients were estimated from uncensored data. (5) Censored frames were interpolated with the Lomb-Scargle periodogram (Power et al., 2014). (6) The data underwent bandpass filtering (0.009Hz – 0.08Hz). (7) Lastly, the data was then projected onto FreeSurfer fsaverage6 surface space and smoothed using a 6 mm full-width half maximum kernel. Task-fMRI data was processed in the same way as the resting-state fMRI data.

For the SINGER dataset, we processed the functional data with the following steps. (1) Removal of the first 4 frames; (2) Slice time correction; (3) Motion correction and outlier detection: frames with FD > 0.3mm or DVARS > 60 were flagged as censored frames. 1 frame before and 2 frames after these volumes were flagged as censored frames. Uncensored segments of data lasting fewer than five contiguous frames were also labeled as censored frames. Runs with over half of the frames censored were removed; (4) Correcting for susceptibility-induced spatial distortion; (5) Multi-echo denoising (DuPre et al., 2021); (6) Alignment with structural image using boundary-based registration (Greve et al., 2009); (7) Global, white matter and ventricular signals, 6 motion parameters, and their temporal derivatives were regressed from the functional data. Regression coefficients were estimated from uncensored data.; (8) Censored frames were interpolated with the Lomb-Scargle periodogram (Power et al., 2014). (9) The data underwent bandpass filtering (0.009Hz – 0.08Hz). (10) Lastly, the data was then projected onto FreeSurfer fsaverage6 surface space and smoothed using a 6 mm full-width half maximum kernel.

For the TCP dataset, the details of data processing can be found elsewhere (Chopra et al., 2024). Briefly, the functional data was processed by following the HCP minimal processing pipeline with ICA-FIX, followed by GSR. The processed data was then projected onto MNI space.

For the MDD dataset, we processed the functional data with the following steps. (1) Slice time correction; (2) Motion correction, (3) Normalization for global mean signal intensity; (4) Alignment with structural image using boundary-based registration (Greve et al., 2009); (5) Linear detrending and bandpass filtering (0.01-0.08 Hz), and (6) Global, white matter and ventricular signals, 6 motion parameters, and their temporal derivatives were regressed from the functional data. (7) Lastly, the data was then projected onto FreeSurfer fsaverage6 surface space and smoothed using a 6 mm full-width half maximum kernel.

For the ADNI dataset, we processed the functional data with the following steps. (1) Slice time correction; (2) Motion correction; (3) Alignment with structural image using boundary-based registration (Greve et al., 2009); (4) Global, white matter and ventricular signals, 6 motion parameters, and their temporal derivatives were regressed from the functional data. (5) Lastly, the data was then projected onto FreeSurfer fsaverage6 surface space and smoothed using a 6 mm full-width half maximum kernel.

We derived a 419 × 419 RSFC matrix for each HCP and ABCD participant using the first *T* minutes of scan time. The 419 regions consisted of 400 parcels from the Schaefer parcellation (Schaefer et al., 2018), and 19 subcortical regions of interest (Fischl et al., 2002). For the HCP, ABCD and TCP datasets, *T* was varied from 2 to the maximum scan time in intervals of 2 minutes. This resulted in 29 RSFC matrices per participant in the HCP dataset (generated from using the minimum amount of 2 minutes to the maximum amount of 58 minutes in intervals of 2 minutes), 10 RSFC matrices per participant in the ABCD dataset (generated from using the minimum amount of 2 minutes to the maximum amount of 20 minutes in intervals of 2 minutes), and 13 RSFC matrices per participant in the TCP dataset (generated from using the minimum amount of 2 minutes to the maximum amount of 26 minutes in intervals of 2 minutes).

In the case of the MDD dataset, the total scan time was an odd number (23 minutes), so T was varied from 3 to the maximum of 23 minutes in intervals of 2 minutes, which resulted in 11 RSFC matrices per participant. For SINGER, ADNI and ABCD task-fMRI data, because the scans were relatively short (around 10 minutes), T was varied from 2 minutes the maximum scan time in intervals of 1 minute. This resulted in 9 RSFC matrices per participant in the SINGER datasets (generated from using the minimum amount of 2 minutes to the maximum amount of 10 minutes), 8 RSFC matrices per participant in the ADNI datasets (generated from using the minimum amount of 2 minutes to the maximum amount of 9 minutes), 9 RSFC matrices per participant in the ABCD n-back task (from using the minimum amount of 2 minutes to the maximum amount of 9.65 minutes), 11 RSFC matrices per participant in the ABCD SST task (from using the minimum amount of 2 minutes to the maximum amount of 11.65 minutes) and 10 RSFC matrices per participant in the ABCD MID task (from using the minimum amount of 2 minutes to the maximum amount of 10.74 minutes).

We note that the above preprocessed data was collated across multiple labs, and even within the same lab, datasets were processed by different individuals many years apart. This led to significant preprocessing heterogeneity across datasets. For example, raw FD was used in the HCP dataset because it was processed many years ago, while the more recently processed ABCD dataset utilized a filtered version of FD, which has been shown to more effective. Another variation is that some datasets were projected to fsaverage space, while other datasets were projected to MNI152 and fsLR space.

### Prediction workflow

The RSFC generated from the first *T* minutes were used to predict each phenotypic measures (previous section) using kernel ridge regression (KRR; He et al., 2020) within an inner-loop (nested) cross-validation procedure.

Let us illustrate the procedure using the HCP dataset (Extended Data Fig. 1). We began with the full set of participants. A 10-fold nested cross-validation procedure was used. Participants were divided in 10 folds (first row of Extended Data Fig. 1). We note that care was taken so siblings were not split across folds, so the 10 folds were not exactly the same sizes. For each of 10 iterations, one fold was reserved for testing (i.e., test set), while the remainder was used for training (i.e., training set). Since there were 792 HCP participants, the training set size was roughly 792 × 0.9 ≈ 700 participants. The KRR hyperparameter was selected via a 10-fold cross-validation of the training set. The best hyperparameter was then used to train a final KRR model in the training set and applied to the test set. Prediction accuracy was measured using Pearson’s correlation and coefficient of determination (Chen et al., 2022).

The above analysis was repeated with different training set sizes achieved by subsampling each training fold (second and third rows of Extended Data Fig. 1), while the test set remained identical across different training set sizes, so the results are comparable across different training set sizes. The training set size was subsampled from 200 to 600 (in intervals of 100). Together with the full training set size of approximately 700 participants, there were 6 different training set sizes, corresponding to 200, 300, 400, 500, 600 and 700.

The whole procedure was repeated with different values of *T*. Since there were 29 values of *T*, there were in total 29 × 6 sets of prediction accuracies for each phenotypic measure. To ensure robustness, the above procedure was repeated 50 times with different splits of the participants into 10 folds to ensure stability (Extended Data Fig. 1). The prediction accuracies were averaged across all test folds and all 50 repetitions.

The procedure for the other datasets followed the same principle as the HCP dataset. However, the ABCD (rest and task) and ADNI datasets comprised participants from multiple sites.

Therefore, following our previous studies (Chen et al., 2022; Ooi et al., 2022), we combined ABCD participants across the 22 imaging sites into 10 site-clusters and combined ADNI participants across the 71 imaging sites into 20 site-clusters (Supplementary Table 5). Each site-cluster has at least 227, 156 and 29 participants in ABCD (rest), ABCD (task) and ADNI datasets respectively.

Instead of the 10-fold inner-loop (nested) cross-validation procedure in the HCP dataset, we performed a leave-3-site-clusters-out inner-loop (nested) cross-validation (i.e., 7 site-clusters are used for training and 3 site-clusters are used for testing) in the ABCD (rest and task) dataset. The hyperparameter was again selected using a 10-fold CV within the training set. This nested cross-validation procedure was performed for every possible split of the site clusters, resulting in 120 replications. The prediction accuracies were averaged across all 120 replications.

We did not perform a leave-one-site-cluster-out procedure because the site-clusters are “fixed”, so the cross-validation procedure can only be repeated 10 times under a leave-one-site-cluster-out scenario (instead of 120 times). Similarly, we did not go for leave-two-site-clusters-out procedure because that will only yield a maximum of 45 repetitions of cross-validation. On the other hand, if we left more than 3 site clusters out (e.g., leave-5-site-clusters-out), we could achieve more cross-validation repetitions, but at the cost of reducing the maximum training set size. Therefore, we opted for the leave-3-site-clusters-out procedure, consistent with our previous study (Chen et al., 2022).

To be consistent with the ABCD dataset, for the ADNI dataset, we also performed a leave-3-site-clusters-out inner-loop (nested) cross-validation procedure. This procedure was performed for every possible split of the site clusters, resulting in 1140 replications. The prediction accuracies were averaged across all 1140 replications.

We also performed 10-fold inner-loop (nested) cross-validation procedure in the TCP, MDD and SINGER datasets. Although the data from the TCP and MDD datasets were acquired from multiple sites, the number of sites was much smaller (2 and 5 respectively) than that of the ABCD and ADNI datasets. Therefore, we were unable to use the leave-some-site-out cross-validation strategy because that would reduce the training set size by too much. Therefore, we ran a 10-fold nested cross-validation strategy (similar to the HCP). However, we regress sites from the target phenotype in the training set, which were then applied to the test set. In other words, our prediction was performed on the residuals of phenotypes after site regression. Site regression was unnecessary for the SINGER dataset as the data was only collected from a single site. The rest of the prediction workflow was the same as the HCP dataset, except for the number of repetitions. Since TCP, MDD and SINGER datasets had smaller sample size than the HCP dataset, the 10-fold cross-validation was repeated 350 times. The prediction accuracies were averaged across all test folds and all repetitions.

Similar to the HCP, the analyses were repeated with different numbers of training participants, ranging from 200 to 1600 ABCD (rest) participants (in intervals of 200). Together with the full training set size of approximately 1800 participants, there were 9 different training set sizes. The whole procedure was repeated with different values of *T*. Since there were 10 values of *T* in the ABCD (rest) dataset, there were in total 10 × 9 values of prediction accuracies for each phenotype. In the case of ABCD (task), the sample size was smaller with maximum training set size of approximately 1600 participants, so there were only 8 different training set sizes.

The ADNI and SINGER datasets had less participants than the HCP dataset, so we decided to sample the training set size more finely. More specifically, we repeated the analyses by varying the number of training participants from the minimum sample size of 100 to the maximum sample size in intervals of 100. For SINGER, the full training set size is ∼580 participants, so there were 6 different training set sizes in total (100, 200, 300, 400, 500, ∼580). For ADNI, the full training set size is ∼530, so there were also 6 different training set sizes in total (100, 200, 300, 400, 500, ∼530).

Finally, TCP and MDD datasets were the smallest, so the training set size was sampled even more finely. More specifically, we repeated the analyses by varying the number of training participants from the minimum sample size of 50 to the maximum sample size in intervals of 25. For TCP, the full training set size is ∼175, so there 6 training set sizes in total (50, 75, 100, 125, 150, 175). For MDD, the full training set size is ∼258, so there 10 training set sizes in total (50, 75, 100, 125, 150, 175, 200, 225, 250, 258).

Current best MRI practices suggest that the model hyperparameter should be optimized (Nichols et al., 2017), so in the current study, we did not consider the case where the hyperparameter was fixed. As an aside, we note that for all analyses, the best hyperparameter was selected using a 10-fold cross-validation within the training set. The best hyperparameter was then used to train the model on the full training set. Therefore, the full training set was used for hyperparameter selection and for training the model. Furthermore, we only needed to select one hyperparameter, while training the model required fitting many more parameters. Therefore, we do not expect the hyperparameter selection to be more dependent on the training set size than training the actual model itself.

We also note that our study focused on out-of-sample prediction within the same dataset, but did not explore cross-dataset prediction (Wu et al., 2023). For predictive models to be clinically useful, these models must generalize to completely new datasets. The best way to achieve this goal is by training models from multiple datasets jointly, so as to maximize the diversity of the training data (Abraham et al., 2017; P. Chen et al., 2023). However, we did not consider cross-dataset prediction in the current study because most studies are not designed with the primary aim of combining the collected data with other datasets.

A full table of prediction accuracies for every combination of sample size and scan time per participant can be found in the supplementary spreadsheet.

### Logarithmic fit of prediction accuracy with respect to total scan duration

By plotting total scan duration (number of training participants × scan duration per participant) against prediction accuracy for each phenotypic measure, we observed that for most measures, scanning beyond 20-30 minutes per participant did not improve prediction accuracy.

Furthermore, visual inspection suggests that a logarithmic curve might fit well to each phenotypic measure when scan time per participant is 30 minutes or less. To explore the universality of a logarithmic relationship between total scan duration and prediction accuracy, prediction accuracy for phenotypic measure *p*, we fitted the function *y_p_* = *z_p_log*(*t_p_*) + *k_p_*, where *y_p_* was the prediction accuracy for phenotypic measure *p*, and *t_p_* is the total scan duration. *z_p_* and *k_p_* were estimated from data by minimizing the square error, yielding *ẑ* and *k̂_p_*.

In addition to fitting the logarithmic curve to different phenotypic measures, the fitting can also be performed with different prediction accuracy measures (Pearson’s correlation or coefficient of determination) and different predictive models (kernel ridge regression and linear ridge accuracies (*y_p_* - *k̂_p_*)/*ẑ_p_* should follow a standard u curve across phenotypic measures, prediction accuracies, predictive models, and datasets. As an example, Fig. 2b shows the normalized prediction performance of the cognitive factors for different prediction accuracy measures (Pearson’s correlation or coefficient of determination) and different predictive models (kernel ridge regression and linear ridge regression) across HCP and ABCD datasets.

Here we have chosen to use kernel ridge regression and linear regression because previous studies have shown that they have comparable prediction performance, and also exhibited similar prediction accuracies as several deep neural networks (He et al., 2020; Chen et al., 2022). Indeed, a recent study suggested that linear dynamical models provide better fit to resting-state brain dynamics (as measured by fMRI and intracranial electroencephalogram) than nonlinear models, suggesting that due to the challenges of in-vivo recordings, linear models might be sufficiently powerful to explain macroscopic brain measurements. However, we note that in the current study, we are not making a similar claim. Instead, our results suggest that the trade-off between scan time and sample size are similar for different regression models, and phenotypic domains, scanners, acquisition protocols, racial groups, mental disorders, age groups, as well as resting-state and task-state functional connectivity.

### Fit of theoretically-motivated model of prediction accuracy, sample size and scan time

We observed that sample size and scan time per participant did not contribute equally to prediction accuracy, with sample size playing a slightly more important role than scan time. To explain this observation, we derived a mathematical relationship relating the expected Pearson’s correlation between noisy brain measurements and non-brain-imaging phenotype with scan time and sample size.

Based on a linear regression model with no regularization and assumptions including (1) stationarity of fMRI (i.e., autocorrelation in fMRI is the same at all timepoints), and (2) prediction errors are uncorrelated with errors in brain measurements, we found that

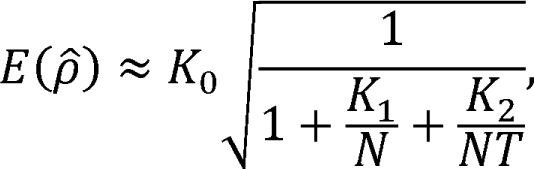

where *E*(ρ̂) is the expected correlation between regression weights estimated from noisy brain measurements and the observed phenotype. *K*_0_ is related to the ideal association between brain measurements and phenotypes, attenuated by phenotype reliability. K_1_ is related to the true association between brain measurements and phenotype, *K*_2_ is related to brain-phenotype prediction errors due to brain measurement inaccuracies. Full derivations can be found in Supplementary Methods Sections 1.1 and 1.2.

Based on the above equation, we fitted the following function 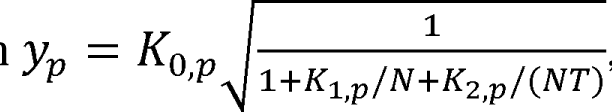, where *y_p_* was the prediction accuracy for phenotypic measure *p*, *N* was the sample size and *T* was the scan time per participant. *K*_0_,*p*,*K*_1_,*p* and *K*_2_,*p* were estimated by minimizing the mean squared error between the above functional form and actual observation of *y_p_* using gradient descent.

### Analysis of non-stationarity

In the original analysis, FC matrices were generated with increasing time *T* based on the original run order. To account for the possibility of state effects, we randomized the order in which the runs were considered for each participant. Since both HCP and ABCD datasets contained 4 runs of resting-fMRI, we generated FC matrices from all 24 possible permutations of run order. For each cross-validation split, the FC matrix for a given participant was randomly sampled from one of the 24 possible permutations. We note that the randomization was independently performed for each participant.

To elaborate further, let us consider an ABCD participant with the original run order (run 1, run 2, run 3, run 4). Each run was 5 minutes long. In the original analysis, if scan time *T* was 5 minutes, then we used all the data from run 1 to compute FC. If scan time *T* was 10 minutes, then we used run 1 and run 2 to compute FC. If scan time *T* was 15 minutes, then we used runs 1, 2 and 3 to compute FC. Finally, if scan time *T* was 20 minutes, we used all 4 runs to compute FC.

On the other hand, after run randomization, for the purpose of this exposition, let us assume this specific participant’s run order had become run 3, run 2, run 4, run 1. In this situation, if scan time *T* was 5 minutes, then we used all data from run 3 to compute FC. If scan time *T* was 10 minutes, then we used run 3 and run 2 to compute FC. If scan time *T* was 15 minutes, then we used runs 3, 2 and 4 to compute FC. Finally, if T was 20 minutes, we used all 4 runs to compute FC.

### Optimizing sample size and scan time within a fixed fMRI budget

To generate Extended Data Fig. 2, we note that given a particular scan cost per hour S and overhead cost per participant O, the total budget for scanning N participants with T min per participant is given by (T/60*S + O) × N. Therefore, given a fixed fMRI budget (e.g., $1M), scan cost per hour (e.g., $500) and overhead cost per participant (e.g., $500), we increase scan time T in one minute interval from 1 to 200, and for each value of T, we can find the largest sample size N, such that the scan costs stayed within the fMRI budget. For each pair of sample size N and scan time T, we can then compute the fraction of maximum accuracy based on Fig. 6a.

### Optimizing sample size and scan time to achieve a fixed prediction accuracy

To generate Fig. 6b, 6c, 7 and 8a, suppose we want to achieve 90% of maximum achievable accuracy, we can find all pairs of sample size and scan time per participant along the 0.9 black contour line in Fig. 6a. For every pair of sample size and scan time, we can then compute the study cost given a particular scan cost per hour (e.g., $500) and a particular overhead cost per participant (e.g., $1000). The optimal scan time (and sample size) with the lowest study cost can then be obtained.

### Brain-wide association reliability workflow

To explore the reliability of univariate brain-wide association analyses (BWAS; Marek et al., 2022), we followed a previously established split-half procedure (Tian et al., 2021; J. Chen et al., 2023).

Let us illustrate the procedure using the HCP dataset (Supplementary Fig. 21a). We began with the full set of participants, which were then divided into 10 folds (first row of Supplementary Fig. 22a). We note that care was taken so siblings were not split across folds, so the 10 folds were not exactly the same sizes. The 10 folds were divided into two non-overlapping sets of 5 folds. For each set of 5 folds and each phenotype, we computed Pearson’s correlation between each RSFC edge and phenotype across participants, yielding a 419 × 419 correlation matrix, which was then converted into a 419 × 419 t-statistic matrix. Split-half reliability between the (lower triangular portions of the symmetric) t-statistic matrices from the two sets of 5 folds was then computed using the intra-class correlation formula (Tian et al., 2021; J. Chen et al., 2023).

The above analysis was repeated with different sample sizes achieved by subsampling each fold (second and third rows of Supplementary Fig. 21a). The split-half sample sizes were subsampled from 150 to 350 (in intervals of 50). Together with the full sample size of approximately 800 participants (corresponding to a split-half sample size of around 400), there were 6 split-half sample sizes corresponding to 150, 200, 250, 300, 350 and 400 participants.

The whole procedure was also repeated with different values of *T*. Since there were 29 values of *T*, there were in total 29 × 6 univariate BWAS split-half reliability values for each phenotype. To ensure robustness, the above procedure was repeated 50 times with different split of the participants into 10 folds to ensure stability (Supplementary Fig. 21a). The reliability values were averaged across all 50 repetitions.

The same procedure was followed in the case of the ABCD dataset, except as previously explained, the ABCD participants were divided into 10 site-clusters. Therefore, the split-half reliability was performed between two sets of 5 non-overlapping site-clusters. In total, this procedure was repeated 126 times since there were 126 ways to divide 10 site-clusters into two sets of 5 non-overlapping site-clusters.

Similar to the HCP, the analyses were repeated with different numbers of split-half participants, ranging from 200 to 1000 ABCD participants (in intervals of 200). Together with the full training set size of approximately 2400 participants (corresponding to a split-half sample size of approximately 1200 participants, there were 6 split-half sample sizes, corresponding to 200, 400, 600, 800, 1000, 1200.

The whole procedure was also repeated with different values of *T*. Since there were 10 values of *T* in the ABCD dataset, there were in total 10 × 6 values univariate BWAS split-half reliability values for each phenotype.

Previous studies have suggested the Haufe-transformed coefficients from multivariate prediction are significantly more reliable than univariate BWAS (Tian et al., 2021; J. Chen et al., 2023).

Therefore, we repeated the above analyses by replacing BWAS with the multivariate Haufe-transform.

A full table of split-half BWAS reliability for each given combination of sample size and scan time per participant can be found in the supplementary spreadsheet.

### Data and Code Availability

The prediction accuracies for each phenotype, sample size N, and scan time T in all six datasets are publicly available (LINK_TO_BE_UPDATED). The raw data for HCP (https://www.humanconnectome.org/), ABCD (https://abcdstudy.org/), TCP (https://openneuro.org/datasets/ds005237 and https://nda.nih.gov/edit_collection.html?id=3552) and ADNI (https://ida.loni.usc.edu/) are publicly available. ABCD parcellated time courses can be found on NDA (LINK_TO_BE_UPDATED). HCP and TCP parcellated time courses can be found on GitHub (LINK_TO_BE_UPDATED). The ADNI user agreement does not allow us to share the ADNI derivatives. The SINGER dataset can be obtained via a data-transfer agreement (https://medicine.nus.edu.sg/macc/projects/singer/). The MDD dataset is available upon request to co-author HL.

Code for this study is publicly available in the GitHub repository maintained by the Computational Brain Imaging Group (https://github.com/ThomasYeoLab/CBIG). Processing pipelines of the fMRI data can be found here (https://github.com/ThomasYeoLab/CBIG/tree/master/stable_projects/preprocessing/CBIG_fMR <u>I_Preproc2016</u>).

Code specific to the analyses in this study can be found here (LINK_TO_BE_UPDATED). Code related to this study was reviewed by co-author TWKT to reduce the chance of coding errors.

## Acknowledgements

Our research is supported by the NUS Yong Loo Lin School of Medicine (NUHSRO/2020/124/TMR/LOA), the Singapore National Medical Research Council (NMRC) LCG (OFLCG19May-0035), NMRC CTG-IIT (CTGIIT23jan-0001), NMRC OF-IRG (OFIRG24jan-0006), NMRC STaR (STaR20nov-0003), Singapore Ministry of Health (MOH) Centre Grant (CG21APR1009), the Temasek Foundation (TF2223-IMH-01), and the United States National Institutes of Health (R01MH120080 & R01MH133334). Any opinions, findings and conclusions or recommendations expressed in this material are those of the authors and do not reflect the views of the Singapore NMRC, MOH or Temasek Foundation.

Data were provided [in part] by the Human Connectome Project, WU-Minn Consortium (Principal Investigators: David Van Essen and Kamil Ugurbil; 1U54MH091657) funded by the 16 NIH Institutes and Centers that support the NIH Blueprint for Neuroscience Research; and by the McDonnell Center for Systems Neuroscience at Washington University.

Data used in the preparation of this article were obtained from the Adolescent Brain Cognitive DevelopmentSM (ABCD) Study (https://abcdstudy.org), held in the NIMH Data Archive (NDA). This is a multisite, longitudinal study designed to recruit more than 10,000 children age 9-10 and follow them over 10 years into early adulthood. The ABCD Study® is supported by the National Institutes of Health and additional federal partners under award numbers U01DA041048, U01DA050989, U01DA051016, U01DA041022, U01DA051018, U01DA051037, U01DA050987, U01DA041174, U01DA041106, U01DA041117, U01DA041028, U01DA041134, U01DA050988, U01DA051039, U01DA041156, U01DA041025, U01DA041120, U01DA051038, U01DA041148, U01DA041093, U01DA041089, U24DA041123, U24DA041147. A full list of supporters is available at https://abcdstudy.org/federal-partners.html. A listing of participating sites and a complete listing of the study investigators can be found at https://abcdstudy.org/consortium_members/. ABCD consortium investigators designed and implemented the study and/or provided data but did not necessarily participate in the analysis or writing of this report. This manuscript reflects the views of the authors and may not reflect the opinions or views of the NIH or ABCD consortium investigators. The ABCD data repository grows and changes over time. The ABCD data used in this report came from http://dx.doi.org/10.15154/1504041.

Data collection and sharing for the Alzheimer’s Disease Neuroimaging Initiative (ADNI) is funded by the National Institute on Aging (National Institutes of Health Grant U19AG024904). The grantee organization is the Northern California Institute for Research and Education. In the past, ADNI has also received funding from the National Institute of Biomedical Imaging and Bioengineering, the Canadian Institutes of Health Research, and private sector contributions through the Foundation for the National Institutes of Health (FNIH) including generous contributions from the following: AbbVie, Alzheimer’s Association; Alzheimer’s Drug Discovery Foundation; Araclon Biotech; BioClinica, Inc.; Biogen; BristolMyers Squibb Company; CereSpir, Inc.; Cogstate; Eisai Inc.; Elan Pharmaceuticals, Inc.; Eli Lilly and Company; EuroImmun; F. Hoffmann-La Roche Ltd and its affiliated company Genentech, Inc.; Fujirebio; GE Healthcare; IXICO Ltd.; Janssen Alzheimer Immunotherapy Research & Development, LLC.; Johnson & Johnson Pharmaceutical Research & Development LLC.; Lumosity; Lundbeck; Merck & Co., Inc.; Meso Scale Diagnostics, LLC.; NeuroRx Research; Neurotrack Technologies; Novartis Pharmaceuticals Corporation; Pfizer Inc.; Piramal Imaging; Servier; Takeda Pharmaceutical Company; and Transition Therapeutics. Data used in preparation of this article were obtained from the Alzheimer’s Disease Neuroimaging Initiative (ADNI) database (adni.loni.usc.edu). As such, the investigators within the ADNI contributed to the design and implementation of ADNI and/or provided data but did not participate in the analysis or writing of this report. A complete listing of ADNI investigators can be found at: http://adni.loni.usc.edu/wp-content/uploads/how_to_apply/ADNI_Acknowledgement_List.pdf.

## Author Contributions

LQRO, CO, RK and BTTY conceptualized the study and designed the methodology. LQRO, RK preprocessed the HCP and ABCD datasets. KHY, FJ and JSXC preprocessed the SINGER dataset. SC and CC preprocessed the TCP dataset. QH, JR and HL preprocessed the MDD dataset. NF and SNR preprocessed the ADNI dataset. LQRO carried out the analysis in the HCP and ABCD datasets. SZ carried out the analysis in the SINGER, TCP and ADNI datasets. QH carried out the analysis in the MDD dataset. TEN derived the theoretical models in the study.

TWKT and RK reviewed the code utilized in the study. LQRO, CO and BTTY wrote the original draft. All authors reviewed and edited the final manuscript.

