## Extended Data for "Longer scans boost prediction and cut costs in brain-wide association studies"

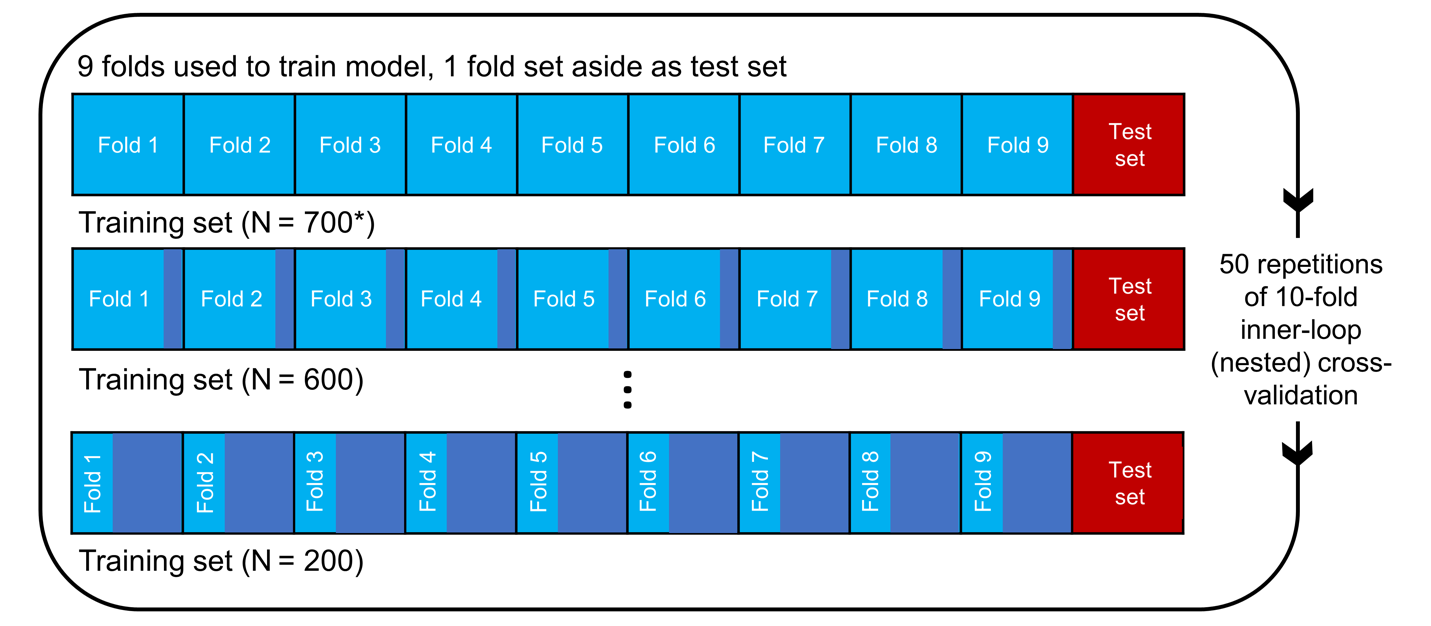

**Extended Data Fig. 1 | Prediction workflow for the HCP dataset.** The participants were split into 10 folds. One fold was set aside to be the test set. The remaining folds comprised the training set. Cross-validation was performed on the training set to select the best hyperparameter. The best hyperparameter was then used to fit a final model from the full training set, which was then used to predict phenotypes in the test set. To vary training set size, each training fold was subsampled and the whole inner-loop nested cross-validation procedure was repeated with the resulting smaller training set. As shown in the panel, the test set remained the same across different training set sizes, so that prediction accuracies were comparable across different sample sizes. Each fold took a turn to be the test set (i.e., 10-fold inner-loop nested cross-validation) and the procedure was repeated with different amounts of fMRI data per participant T (not shown in panel). For stability, the entire procedure was repeated 50 times and averaged. A similar workflow was used in the ABCD dataset (see Methods). We note that in the case of HCP, care was taken so siblings were not split across folds, while in the case of ABCD, participants from the same site were not split across folds.

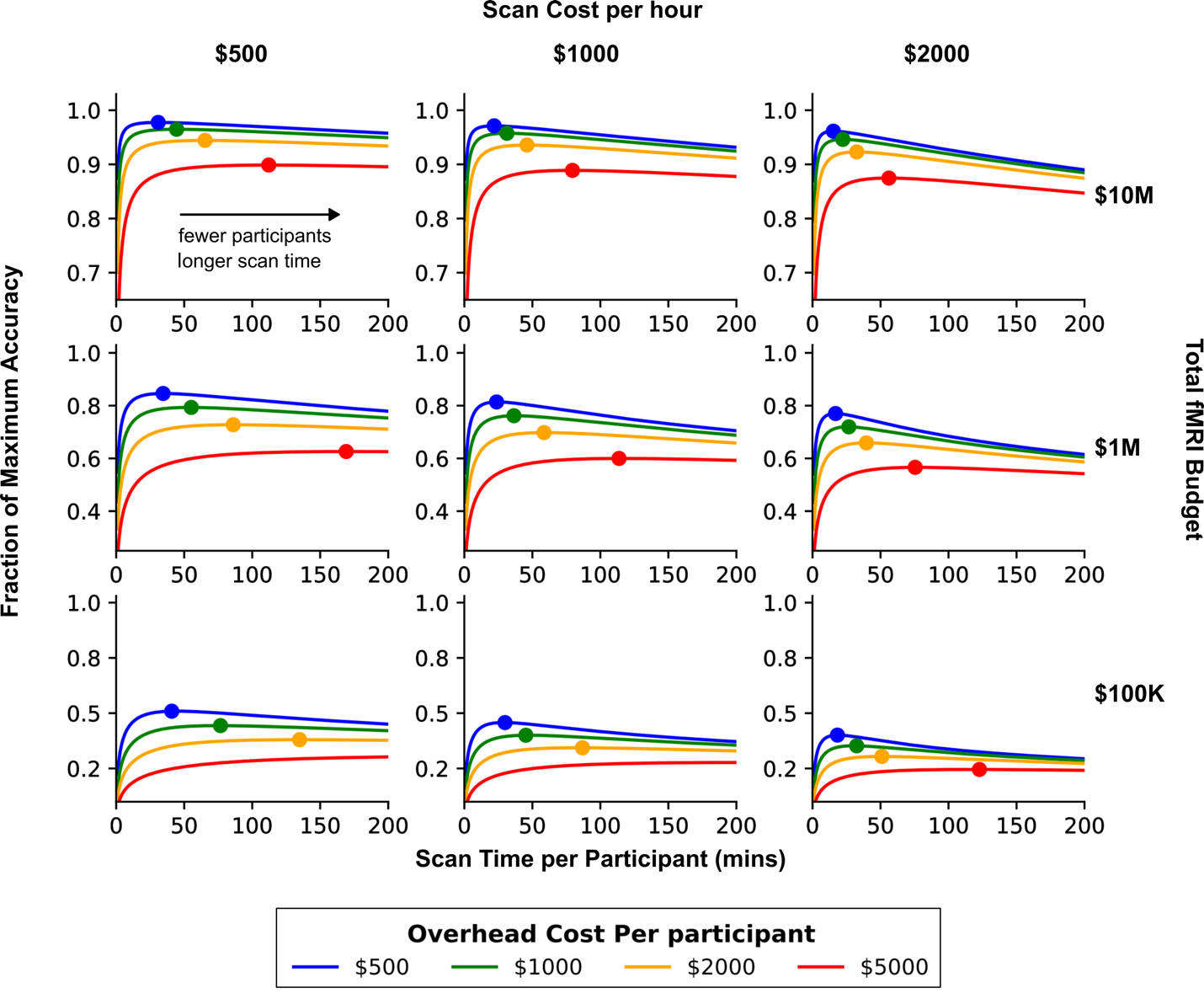

### **Extended Data Fig. 2 |** **Fraction of maximum achievable prediction accuracy as a function of total fMRI budget, scan cost per hour and overhead cost per participant.** The three columns correspond to scan cost per hour of $500, $1000 and $2000 respectively. The three rows correspond to total fMRI budget of $10M, $1M and $100K respectively. The different colored lines correspond to different overhead cost per participant. Each curve shows the fraction of maximum prediction accuracy as a function of scan time per participant for a given overhead cost per participant and scan cost per hour, while keeping to within the total fMRI budget. On each curve, the solid circle indicates location of the maximum prediction accuracy. Circles are not shown if optimal scan time was beyond the edge of the graph (i.e., more than 200 minutes of scan time).

### **Extended Data Table 1 |** This table expands Extended Data Fig. 2 for a wider range of fMRI budgets, scan cost per hour and overhead cost per participant. More specifically, for a fixed fMRI budget, scan cost per hour and overhead cost per participant, the goal is to find the optimal scan time and sample size in order to maximize the prediction accuracy. Entries in the table shows the optimal scan time in minutes.

| Overhead cost per participant | Scan cost per hour | Total fMRI budget | | | | | | |
| --- | --- | --- | --- | --- | --- | --- | --- | --- |
|  |  | 100K | 250K | 1M | 2.5M | 10M | 25M | 100M |
| 500 | 500 | 40.0 | 40.0 | 34.5 | 32.5 | 31.5 | 30.5 | 30.5 |
|  | 1000 | 30.0 | 25.5 | 24.5 | 22.5 | 22.0 | 21.5 | 21.5 |
|  | 1500 | 20.0 | 20.0 | 20.0 | 18.5 | 18.0 | 17.5 | 17.5 |
|  | 2000 | 19.5 | 18.0 | 16.0 | 16.0 | 15.5 | 15.5 | 15.0 |
|  | 2500 | 18.0 | 15.5 | 15.0 | 14.5 | 14.0 | 13.5 | 13.5 |
| 1000 | 500 | 80.0 | 67.4 | 56.4 | 50.5 | 44.5 | 44.0 | 43.0 |
|  | 1000 | 40.0 | 40.0 | 38.5 | 35.0 | 31.5 | 31.0 | 30.5 |
|  | 1500 | 40.0 | 33.0 | 30.5 | 28.5 | 25.5 | 25.0 | 25.0 |
|  | 2000 | 30.0 | 28.0 | 25.5 | 24.0 | 22.5 | 22.0 | 21.5 |
|  | 2500 | 24.0 | 26.0 | 24.0 | 21.5 | 20.0 | 19.5 | 19.0 |
| 2000 | 500 | 159.8 | 100.9 | 86.9 | 76.4 | 66.4 | 62.4 | 60.9 |
|  | 1000 | 79.9 | 67.4 | 56.4 | 50.5 | 45.5 | 44.5 | 43.0 |
|  | 1500 | 52.9 | 62.4 | 49.0 | 40.5 | 37.5 | 36.5 | 35.5 |
|  | 2000 | 40.0 | 47.0 | 40.0 | 37.5 | 32.5 | 31.5 | 30.5 |
|  | 2500 | 37.5 | 37.5 | 35.0 | 32.0 | 29.0 | 28.0 | 27.5 |
| 5000 | 500 | 322.7 | 233.3 | 159.3 | 133.4 | 110.9 | 102.9 | 96.9 |
|  | 1000 | 299.7 | 199.8 | 128.4 | 94.4 | 79.9 | 74.4 | 69.9 |
|  | 1500 | 199.8 | 132.9 | 85.4 | 78.4 | 63.4 | 60.0 | 55.9 |
|  | 2000 | 149.9 | 99.9 | 80.4 | 65.4 | 55.3 | 51.4 | 49.0 |
|  | 2500 | 119.9 | 79.9 | 64.4 | 56.4 | 50.5 | 45.5 | 44.0 |
| 10000 | 500 | 500.0 | 466.5 | 299.7 | 228.3 | 180.8 | 151.3 | 140.9 |
|  | 1000 | 399.6 | 282.2 | 189.3 | 149.9 | 112.9 | 110.9 | 100.9 |
|  | 1500 | 266.2 | 187.8 | 171.3 | 125.9 | 99.9 | 90.4 | 81.9 |
|  | 2000 | 199.8 | 168.3 | 128.4 | 116.4 | 84.4 | 77.4 | 69.9 |
|  | 2500 | 159.8 | 134.9 | 112.9 | 94.9 | 80.4 | 69.4 | 63.4 |

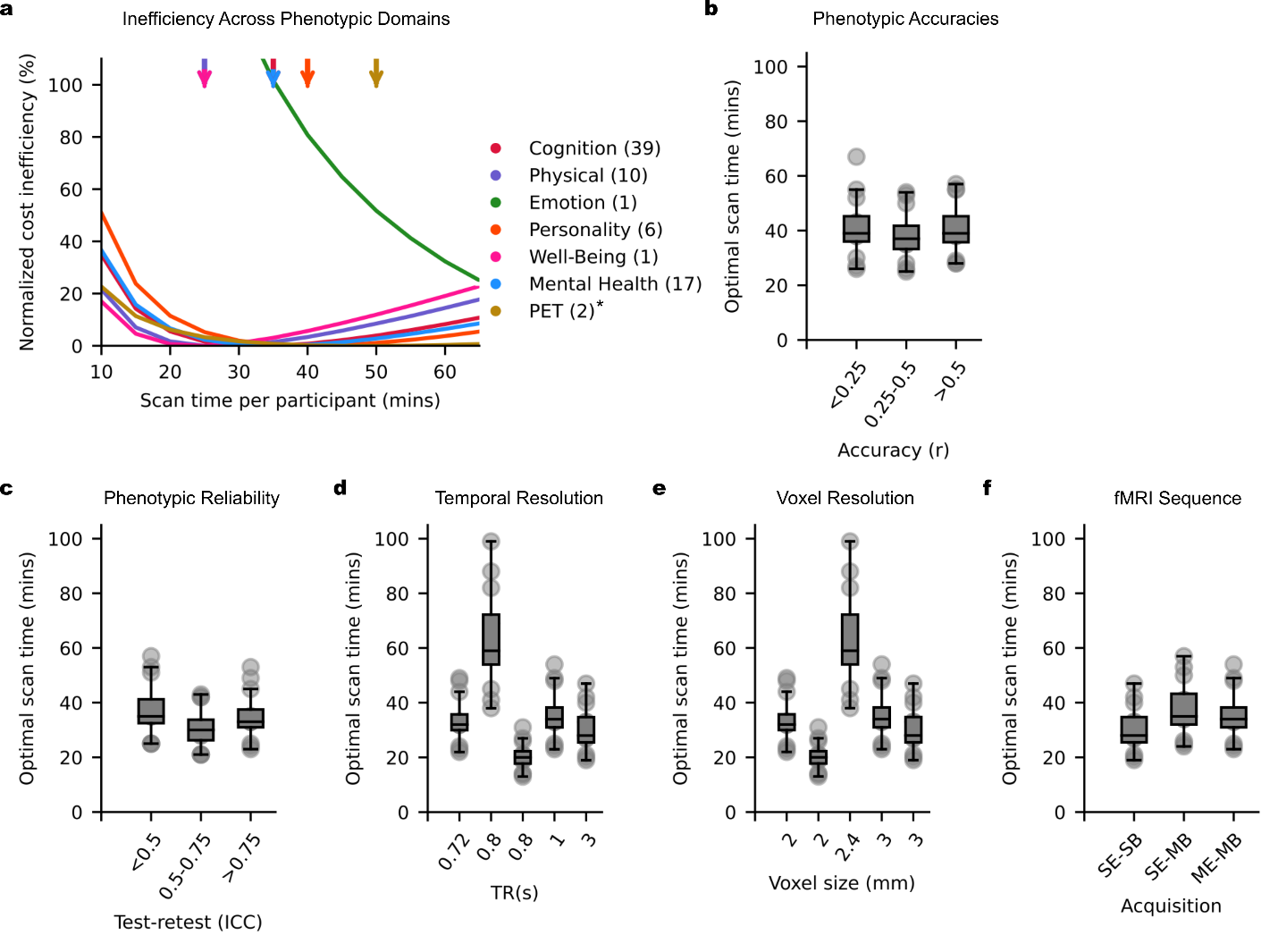

### **Extended Data Fig. 3 | Variation in cost inefficiency and optimal scan time across phenotypic domains and common scan parameters. a.** Cost inefficiency as a function of scan time for various phenotypic domains across nine resting-fMRI and task-fMRI datasets. For the purpose of this plot, the positron emission tomography (PET) curve was based on a more realistic overhead cost of $5000 or $10000 per participant (instead of $500 or $1000) that was used for other phenotypic measures. Arrows indicate scan times with the lowest budgets. **b.** Optimal scan time as a function of phenotypic prediction accuracies. This analysis was obtained by sorting the maximum prediction accuracies (based on resting-state FC) of 19 HCP and 17 ABCD phenotypes into three bins. **c.** Optimal scan time as a function of phenotypic test-retest reliability. This analysis was obtained by considering 41 participants from the HCP, where the same phenotypic measures were collected twice (several months apart), allowing us to compute the test-retest reliability of the HCP phenotypes. We sorted the phenotypic test-retest reliability of the 19 HCP phenotypes into three bins. **d.** Optimal scan time as a function of repetition time (TR). **e.** Optimal scan time as a function of voxel size. **f.** Optimal scan time as a function of MRI acquisition. SE-SB: single-echo single-band; SE-MB: single-echo multi-band; ME-MB: multi-echo multi-band. We note that panels (b) to (f) only considered resting-state FC. Furthermore, the ADNI dataset was excluded from the analysis in panels (d), (e) and (f) because it included both single-band and multi-band data with different TRs and voxel sizes. Similar to Fig. 6c, for visualization, the curves in panel (a) are normalized by subtracting the cost inefficiency of the best possible fixed scan time (of each curve), so that the best possible fixed scan time is centered at zero.
