## Supplemental Material for "Longer scans boost prediction and cut costs in brain-wide association studies"

### **Supplementary methods**

This section provides approximations that justify the form of key outcomes as a function of sample size $N$ and scan time $T$. We consider two types of outcomes, the correlation between a linear prediction of non-brain-imaging phenotype (henceforth referred to as phenotype) using functional connectivity (FC), and the reliability of edge-wise FC-phenotypic correlation after participant-wise data splitting.

It is important to note that the following derivations are general and not limited to functional connectivity, and are in fact applicable to the relationship between any phenotype with measurements from any sensor (not necessarily MRI).

#### 1.1 **Preliminaries: 1-edge results**

Define the target phenotype variable for participant $i$ as

$Y_{i}=\psi_{i}+\xi_{i}$	(S1)

for $i=1,\ldots,N$, where $\psi_{i}$ is the noise-free, latent feature that $Y$ attempts to measure and $\xi_{i}$ is the random error. For a typical phenotypic trait, $\psi_{i}$ could be thought as a measurement obtainable if you had multiple days and endless tests to acquire each participant; $\xi_{i}$ is the divergence between that ideal value and $Y_{i}$. Let the variance of the true measure be $\sigma_{\psi}^{2}$, and for the measurement error $\sigma_{\xi}^{2}$; the intraclass correlation reliability of phenotype is then $R\left( Y \right)=\frac{\sigma_{\psi}^{2}}{(\sigma_{\psi}^{2}+\sigma_{\xi}^{2})}$.

Let the FC measure for participant $i$, edge $j$ based on $T$ scans be $x_{Tij}$, formed into a length-$J$ row vector $x_{Ti}$. The observable FC is also a noisy measure of the ideal measure; for scan length $T$, participant $i$ and edge $j$ this is

$x_{Tij}=\theta_{ij}+\epsilon_{\mathrm{Tij}}$	(S2)

where $\theta_{ij}$ is the noise-free FC measure. The values $\theta_{ij}$ can be considered as the FC value you’d obtain if you could leave the participant in the scanner so long that measurement error vanishes (or is negligible). The measurement error $\epsilon_{Tij}$is specific to the acquisition time $T$, as precision will increase with longer scan time. Let the variance of true FC be $\sigma_{\theta_{j}}^{2}$ (i.e. the participant-to-participant variability in true FC at edge $j$); the measurement error for FC is more involved.

If fMRI time series had no temporal autocorrelation, the sampling variance of Pearson’s correlation would be well approximated by

$\frac{1}{T}\left( 1-\theta_{ij}^{2} \right)^{2}$	(S3)

However, as covered in detail in Afyouni et al., 2019, the distinct temporal autocorrelation in each node and the cross-correlation (at all lags) influence the sampling variance of Pearson’s correlation in complex fashion. For simplicity, we just assume that there is some normalized variance $\tau_{j}^{2}$ such that $V\left( \epsilon_{Tij} \right)=\frac{\tau_{j}^{2}}{T}$ (strictly, we should keep track of participant-specific variance $\frac{\tau_{ij}^{2}}{T}$ since it depends on each participant’s correlation $\theta_{ij}$ and autocorrelation; however, this is the first of many simplification we make to obtain tractable results). Thus the intraclass correlation reliability of FC is then $R\left( x_{Tj} \right)=\frac{\sigma_{\theta_{j}}^{2}}{\sigma_{\theta_{j}}^{2}+\frac{\tau_{j}^{2}}{T}}$. The ideal FC-phenotype correlation for edge $j$ is

$\rho_{j}=\text{corr}(\theta_{ij},\psi_{i})$	(S4)

However, we cannot directly observe these noise-free measurements, as each is corrupted by measurement noise. Due to a classic result by Spearman, 1904, we know that when corrupted measures are used to compute $\hat{\rho}_{j}=\hat{\text{corr}}(x_{Tij},Y_{i})$ the result is biased, with

$E\left( \hat{\rho}_{j} \right) = \rho_{j}\sqrt{R(X_{Tj})}\sqrt{R(Y)}$	(S5)

$= \rho_{j}\sqrt{\frac{1}{1+\frac{(\tau_{j}^{2}/\sigma_{\theta_{j}}^{2})}{T}}}\sqrt{\frac{1}{1+\frac{\sigma_{\xi}^{2}}{\sigma_{\psi}^{2}}}}$	(S6)

This shows the dependence of edge-wise FC-phenotype association on scan length T. If no information is available on the reliability of the phenotype, we can simply act as if there is no measurement error $(\sigma_{\xi}^{2}=0)$ and then there is no dependence on phenotype variability $\sigma_{\psi}^{2}$.

Below we will also need the variance of $\hat{\rho}_{j}$ , which is simply the variance for Pearson’s correlation for variables where the true correlation is $\rho_{j}\sqrt{R\left( x_{Tj} \right)R(Y)}$,

$V\left( \hat{\rho}_{j} \right)=\frac{1}{N}\left( 1-\rho_{j}^{2}R\left( X_{Tj} \right)R\left( Y \right) \right)^{2}$	(S7)

which is the same as (S3) except applied over participants.

Note that it will be useful to approximate this with a 2^nd^ order Taylor series approximation for $f\left( t \right)=\left( 1-t^{2} \right)^{2}$ about $t=0$, $1-2t^{2}$, here

$V\left( \hat{\rho}_{j} \right)\approx\frac{1}{N}\left( 1-2\rho_{j}^{2}R\left( X_{Tj} \right)R\left( Y \right) \right)$	(S8)

which we find to be fairly accurate up through $t=\rho_{j}\sqrt{R\left( X_{Tj} \right)R(Y)}=0.5$, where as a reminder, $\rho_{j}\sqrt{R\left( X_{Tj} \right)R(Y)}$ is the true correlation between FC edge $j$ and a phenotype for a fMRI acquisition of length $T$. We will use this approximation for the reliability analysis (Supplementary Methods S1.3). Obviously, we do not know the true correlation between FC edge $j$ and a phenotype. However, we can compute the actual correlation between FC and a phenotype to check the quality of the approximation. The phenotype with the strongest correlation with FC is the cognitive factor score. In the case of the HCP dataset, across all edges, the largest absolute correlation between FC and the cognitive factor was 0.27, while in the case of the ABCD dataset, across all edges, the largest absolute correlation between FC and the cognitive factor was 0.22. As the strongest correlation is much smaller than 0.5, we believe that our approximation is good.

#### 1.2 **FC-phenotype prediction accuracy**

While the body of the paper presents results for kernel ridge regression, we found that linear ridge regression gave very similar results. The analysis here is only for linear regression and is provided to motivate the role that $N$ and $T$ might play in the phenotype of FC-phenotype prediction accuracy measured with correlation.

First, for subject $i$ write the linear predictor of the noise-free phenotype $\psi_{i}$ using noise-free FC $\theta_{i}$ as $\theta_{i}\beta_{i}$, where $\beta_{i}$ is the ideal length-$J$ vector of regression weights. Define the true, ideal correlation as

$\rho=\text{corr}(\theta_{i}\beta_{i},\psi_{i})$.	(S9)

In practice we can at most compute $\hat{\rho}=\hat{\text{corr}}(x_{Ti}\hat{\beta},Y_{i})$, for which previously stated results give us

$E\left( \hat{\rho} \right)=\rho\sqrt{R\left( x_{T}\hat{\beta} \right)}\sqrt{R\left( Y \right)}$	(S10)

With slight abuse of notation, here we use $\rho$ for the regression-based noise free FC-phenotype association over edges, and $\hat{\rho}$ as the noise-corrupted association; these are distinct from the previous $\rho_{j}$ and $\hat{\rho}_{j}$, the edgewise noise-free and noise-corrupted FC-phenotype associations respectively.

Calculating the sample variance of $x_{Ti}\hat{\beta}$ to compute $R(X_{T}\hat{\beta})$ is challenging: standard regression results don’t apply since they neglect using the noisy FC $x_{Ti}$ instead of the ideal $\theta_{i}$, in what is known as an “Errors-in-Variables” problem.

To find the sample variance of $x_{Ti}\hat{\beta}$ when OLS is used to estimate a errors-in-variables model we rely on the results from Gleser et al., 1987 (GCG). Following GCG, the regression is partitioned into known (error-free) variables $F_{1}(N\times P)$ and unobserved variables $F_{2}(N\times J)$, for which we observe a noisy version $X$:

$Y=F_{1}\beta_{1}+F_{2}\beta_{2}+e$	(S11)

$X=F_{2}+U$	(S12)

All useful results require normality of the errors $e$ and the corrupting noise $U$, with the joint distribution of these stochastic components having $\left( 1+J \right)\times(1+J)$ covariance

$\text{Cov}\left( \left[ e_{i},u_{i} \right] \right)=\Sigma= \left[ \begin{matrix} \sigma_{11}^{2} & \sigma_{12} \\ \sigma_{12}^{\top} & \Sigma_{22} \end{matrix} \right]$	(S13)

where $u_{i}$ is the $J$-vector of covariate errors, $\sigma_{22}^{2}$ is the variance of the residual error (in the ideal model with $F_{1}$ and $F_{2}$), $\sigma_{12}$ the $1\times J$ covariance between the residual error and the $F_{2}$ corrupting noise, and $\Sigma_{22}$ the $J\times J$ covariance of the corrupting noise.

GCG gives the properties of the OLS regression with $\beta^{\top}=(\beta_{1}^{\top},\beta_{2}^{\top})$ when the design matrix $[F_{1} X]$ is used. The results depend on the limiting mean-squares of the covariates

$\lim_{N\to\infty} \frac{1}{N}\left[ \begin{matrix} F_{1}^{\top}F_{1} & F_{1}^{\top}F_{2} \\ F_{2}^{\top}F_{1} & F_{2}^{\top}F_{2} \end{matrix} \right]= \left[ \begin{matrix} \Delta_{11} & \Delta_{12} \\ \Delta_{12}^{\top} & \Delta_{22} \end{matrix} \right]$	(S14)

With errors-in-variables OLS is biased, with $\hat{\beta}$ having asymptomatic expectation

$\beta+\left[ \begin{matrix} \Delta_{11} & \Delta_{12} \\ \Delta_{12}^{\top} & \Delta_{22} \end{matrix} \right]^{-1}\left( \begin{matrix} 0 \\ \gamma\end{matrix} \right)=\beta+\left[ \begin{matrix} -\Delta_{11}^{-1}\Delta_{12}\left( \Delta_{22}-\Delta_{12}^{\top}\Delta_{11}^{-1}\Delta_{12} \right)^{-1}\gamma\\ \left( \Delta_{22}-\Delta_{12}^{\top}\Delta_{11}^{-1}\Delta_{12} \right)^{-1}\gamma\end{matrix} \right]$	(S15)

where

$\gamma=\sigma_{12}^{\top}-\Sigma_{22}\beta_{2}$	(S16)

We use GCG’s Theorem 2 that unfortunately is narrowly stated for contrasts $C\hat{\beta}$ where OLS is unbiased, i.e. where $C$ has a certain form, $C=[I_{P},\Delta_{11}^{-1}\Delta_{12}]$. Theorem 2 states that the asymptotic variance of $C\hat{\beta}$ is

$\text{Cov}\left( C\hat{\beta} \right)=\frac{1}{N}\eta^{\top}\Sigma\eta\Delta_{11}^{-1}$	(S17)

where

$\eta= \left[ \begin{matrix} 1 \\ -(\beta_{2}+Q\gamma) \end{matrix} \right]$	(S18)

$Q = \left( \Delta_{22.1}+\Sigma_{22} \right)^{-1}$	(S19)

$\Delta_{22.1} = \Delta_{22}-\Delta_{12}^{\top}\Delta_{11}^{-1}\Delta_{12}$	(S20)

$\gamma= \sigma_{12}^{\top}-\Sigma_{22}\beta_{2}$	(S21)

In our setting, we have $P=1$ error-free predictor, with $F_{1}$ being a column of $1$’s for the intercept, and thus $\Delta_{11}=1$ (a scalar) and $\Delta_{12}=\bar{\theta}$ is the $1\times J$ vector of means of the noiseless FC edges over $N$ participants; the $J$ noise-corrupted FC measurements make up $X$, and the elements of $U$ are exactly the measurement errors $\epsilon_{Tij}$. A common assumption is that the model error $e$ is uncorrelated with the measurement noise, and hence $\sigma_{12}=0$. Further, we can capture the $T$-dependence of the measurement error by assuming,

$\Sigma_{22}=\frac{1}{T}\Sigma_{22}^{*}$	(S22)

that is, that $\Sigma_{22}^{*}$ is a normalized measurement error covariance that scales by $\frac{1}{T}$ to give the actual measurement error. As an aside, under these settings, OLS $\hat{\beta}$ has mean

$\beta+\frac{1}{T}\left[ \begin{matrix} -\bar{\theta}\Delta_{22.1}^{-1}\Sigma_{22}^{*}\beta_{2} \\ \Delta_{22.1}^{-1}\Sigma_{22}^{*}\beta_{2} \end{matrix} \right]$	(S23)

Note in our setting $\Delta_{22.1}$ is the limiting covariance of the (unobserved, noise-free) FC design matrix $F_{2}$. The reason is that in our case $\Delta_{11}^{-1}=1$, so $\Delta_{22.1}=\Delta_{22}-\Delta_{12}^{\top}\Delta_{11}^{-1}\Delta_{12}=E\left( F_{2}^{T}F_{2} \right)-E^{T}\left( F \right)E(F)$, which is the standard variance formula.

In our setting $C=\left[ 1,\bar{\theta} \right]$, which is the intercept plus (true) mean FC. Instead we want, for each participant $i$, $\hat{Y}_{i}=\left[ 1,X_{i} \right]\hat{\beta}$, the linear combination of $\hat{\beta}$ that is the intercept plus the J elements of $\hat{\beta}$ weighted according to that participant’s FC measurements. Hence, strictly, the Theorem 2 result is relevant to the prediction for the average participant (using the noise-free FC), however we use this to gauge the properties of the sampling variance of the prediction.

To simplify the main result, let $\beta_{2}^{*}=\beta_{2}-\left( \Delta_{22.1}+\frac{1}{T}\Sigma_{22}^{*} \right)^{-1}\frac{1}{T}\Sigma_{22}^{*}\beta_{2}$, which note converges to $\beta_{2}$ as $T$ grows. Then in our application, the asymptotic variance is

$\text{Var}\left( C\hat{\beta} \right)=\frac{1}{N}\left[ 1,-{\beta_{2}^{*}}^{\top} \right]\left[ \begin{matrix} \sigma_{11}^{2} & 0^{\top} \\ 0 & \frac{\Sigma_{22}^{*}}{T} \end{matrix} \right]\left[ 1,-{\beta_{2}^{*}}^{\top} \right]^{\top}$	(S24)

$=\frac{1}{N}\sigma_{11}^{2}+\frac{1}{TN}{\beta_{2}^{*}}^{\top}\Sigma_{22}^{*}\beta_{2}^{*}$	(S25)

We can see that this variance has 2 terms: the first is the residual variance in $Y$ not explained by (noiseless) FC; the second contains the contribution of FC measurement error, which we have expressed relative to the normalized $\Sigma_{22}^{*}$, showing the dependence on $\frac{1}{TN}$.

Finally, to compute reliability, we have

$R\left( X_{T}\hat{\beta} \right)=\frac{S_{\theta\beta}^{2}}{S_{\theta\beta}^{2}+\frac{1}{N}\sigma_{11}^{2}+\frac{1}{TN}{\beta_{2}^{*}}^{\top}\Sigma_{22}^{*}\beta_{22}^{*}}$	(S26)

where $S_{\theta\beta}^{2}$ is the “true” variation of interest, the inter-participant variation in the predictions using noise-free FC $\theta$ and ideal regression coefficients $\beta$. Then the expected regression FC-phenotype correlation is

$E\left( \hat{\rho} \right)\approx\rho\sqrt{\frac{S_{\theta\beta}^{2}}{S_{\theta\beta}^{2}+\frac{1}{N}\sigma_{11}^{2}+\frac{1}{TN}{\beta_{2}^{*}}^{\top}\Sigma_{22}^{*}\beta_{22}^{*}}}\sqrt{\frac{\sigma_{\psi}^{2}}{\sigma_{\psi}^{2}+\sigma_{\xi}^{2}}}$	(S27)

#### 1.2.1 **Empirical curve fitting for prediction accuracy**

When we estimate a function of observed correlations as a function of $N$ and $T$

$K_{0}\sqrt{\frac{1}{1+\frac{1}{N}K_{1}+\frac{1}{TN}K_{2}}}$	(S28)

we can interpret $K_{0}=\rho R\left( Y \right)$ as the ideal association attenuated by phenotype reliability. Noting that $\rho^{2}=\frac{S_{\theta\beta}^{2}}{\sigma_{\psi}^{2}}$, the proportion of noise-free phenotype explained by the ideal prediction, and $\sigma_{11}^{2}=\left( 1-\rho^{2} \right)\sigma_{\psi}^{2}$ is the noise-free phenotype variance not explained, then

$K_{1}=\frac{\sigma_{11}^{2}}{S_{\theta\beta}^{2}}=\frac{\left( 1-\rho^{2} \right)\sigma_{\psi}^{2}}{\rho^{2}\sigma_{\psi}^{2}}=\frac{{1-\rho}^{2}}{\rho^{2}}$	(S29)

as the inverse Cohen’s $f^{2}$ of the (noise-free, ideal) prediction ($f^{2}$ is an effect size often similar to $R^{2}$), and

$K_{2}=\frac{{\beta_{2}^{*}}^{\top}\Sigma_{22}^{*}\beta_{2}^{*}}{S_{\theta\beta}^{2}}=\frac{{\beta_{2}^{*}}^{\top}\Sigma_{22}^{*}\beta_{2}^{*}}{\rho^{2}\sigma_{\psi}^{2}}$	(S30)

as the measurement error relative to the variance explained with noise-free ideal prediction.

These interpretations, however, should be tempered by the many assumptions leading up to this result. The principle critical assumptions are $\sigma_{12}=0$, in that ideal prediction errors are uncorrelated with measurement errors, and that we’re using a variance result for the prediction of a participant with the (true) average FC value for each edge, which may not be representative over all.

#### 1.2.2 **Intuition under restrictive independence assumptions**

Returning to $R(x_{T}\hat{\beta})$, note that if we assume independent measurement errors along the $J$ edges, then $\Sigma_{22}^{*}=\text{diag}\left( \left\{ \tau_{j}^{2} \right\} \right)$ and ${\beta_{2}^{*}}^{\top}\Sigma_{22}^{*}\beta_{22}^{*}=\Sigma_{j}{\beta_{2j}^{*}}^{2}\tau_{j}^{2}$. Further, if we can assume that noise-free FC is (1) normalized to unit variance and (2) independent, then $\Delta_{22.1}=I$ and the measurement error contribution ${\beta_{2}^{*}}^{\top}\Sigma_{22}^{*}\beta_{2}^{*}$ further simplifies to $\Sigma_{j}\beta_{j}^{2}\tau_{j}^{2}\left( 1+\frac{\tau_{j}^{2}}{T} \right)^{-2}$. However, we realize that we can only ever normalize $X$ by empirical variance which is an over-estimate of noise-free FC, and that FC edges are highly structured and could never be anywhere near independent unless some very careful thinning was done. However, using $\rho^{2}$ and these restrictive assumptions, we obtain an alternate form of $R(x_{T}\hat{\beta})$ of

$\frac{\rho^{2}}{\rho^{2}+\frac{1}{N}\left( 1-\rho^{2} \right)+\frac{J}{TN}\left\langle\tau_{j}^{2}\left( \frac{1}{1+\frac{\tau_{j}^{2}}{T}} \right)^{2}\left( \frac{\beta_{j}}{\sigma_{\psi}} \right)^{2} \right\rangle}$	(S31)

which clearly shows the contributions of noise-free FC-phenotype association $\left( \rho^{2} \right)$, a $\left( \frac{1}{N} \right)$-weighted contribution of unexplained variation $\left( 1-\rho^{2} \right)$, and a $\left( \frac{1}{TN} \right)$-weighted contribution of measurement error and normalized (noise-free) regression coefficients, where the $\left\langle\cdot\right\rangle$ notation indicates average over edges. As any of $\rho^{2}$, $N$ or $T$ increases, reliability $R(x_{T\hat{\beta}})$ grows to an asymptote of 1; increases in measurement noise decrease reliability through a complex weighted fashion depending on the regression coefficients.

#### **1.3 Edgewise reliability of FC-phenotype association**

We measure edgewise reliability of FC-phenotype association by splitting participants into two groups, computing FC-phenotype association at each of $J$ edges, and then correlating the associations over edges. Consider participants split into groups $A$ and $B$, $\frac{N}{2}$ in each, computing $\hat{\rho}_{Aj}$ and $\hat{\rho}_{Bj}$ for each edge $j$, with a model for these noisy correlations at edge $j$ of

$\hat{\rho}_{Aj}=\rho_{j}+\epsilon_{Aj}$	(S32)

$\hat{\rho}_{Bj}=\rho_{j}+\epsilon_{Bj}$	(S33)

where $\epsilon_{Aj}$ and $\epsilon_{Bj}$ are the random measurement error from the true association value $\rho_{j}$; note that since participants are split, these two errors are independent and from (S3) above, we know the variance is

$V\left( \hat{\rho}_{Aj} \right)=V\left( \hat{\rho}_{Bj} \right)\approx\frac{1}{N/2}\left( 1-\rho_{j}^{2}R\left( X_{Tj} \right)R\left( Y \right) \right)^{2}\approx\frac{1}{N/2}(1-2\rho_{j}^{2}R\left( X_{Tj} \right)R\left( Y \right))$	(S34)

using the (S8) to obtain a simplified form. If we use the true means instead of sample means, the correlation coefficient computed between $\hat{\rho}_{Aj}$ and $\hat{\rho}_{Bj}$ over edges is

$\frac{\sum_{j} \left( \hat{\rho}_{Aj}-\bar{\rho} \right)\left( \hat{\rho}_{Bj}-\bar{\rho} \right)}{\sqrt{\sum_{j} \left( \hat{\rho}_{Aj}-\bar{\rho} \right)^{2}}\sqrt{\sum_{j} \left( \hat{\rho}_{Bj}-\bar{\rho} \right)^{2}}}$	(S35)

where $\bar{\rho}=\frac{1}{J}\sum_{j} \rho_{j}$, the true FC-phenotype association averaged over edges. We proceed by approximating the expectation of this ratio as a ratio of expectations.

The expected value of the numerator is

$E\left( \sum_{j} \left( \hat{\rho}_{Aj}-\bar{\rho} \right)\left( \hat{\rho}_{Bj}-\bar{\rho} \right) \right)=\sum_{j} E((\rho_{j}-\bar{\rho}+\epsilon_{Aj})(\rho_{j}-\bar{\rho}+\epsilon_{Aj}))$	(S36)

$= \sum_{j} \left( E\left( \rho_{j}-\bar{\rho} \right)^{2}+E\left( \left( \rho_{j}-\bar{\rho} \right)\epsilon_{Bj} \right)+E\left( \left( \rho_{j}-\bar{\rho} \right)\epsilon_{Aj} \right)+ E\left( \epsilon_{Aj}\epsilon_{Bj} \right) \right)$	(S37)

$=JS_{\rho}^{2}$	(S38)

where $S_{\rho}^{2}$ is the inter-edge variance of the true FC-phenotype association, and the other terms are zero because the errors are uncorrelated with FC and between participant groups $A$ and $B$.

For the denominator term for $A$ (identical to that for $B$),

$E\left( \sum_{j} \left( \hat{\rho}_{Aj}-\bar{\rho} \right)^{2} \right)=\sum_{j} E\left( \left( \rho_{j}-\bar{\rho}+\epsilon_{Aj} \right)^{2} \right)$	(S39)

$=\sum_{j} \left( E\left( \left( \rho_{j}-\bar{\rho} \right)^{2} \right)+2E\left( \left( \rho_{j}-\bar{\rho} \right)\epsilon_{Aj} \right)+E\left( \epsilon_{Aj}^{2} \right) \right)$	(S40)	$=JS_{\rho}^{2}+\frac{1}{N/2}\sum_{j} \left( 1-\rho_{j}^{2}R\left( X_{Tij} \right)R\left( Y \right) \right)^{2}$	(S41)

$\approx JS_{\rho}^{2}+\frac{J}{N/2}\left( 1-2\langle\rho_{j}^{2}R\left( X_{Tij} \right)\rangle R\left( Y \right) \right)$	(S42)

$=JS_{\rho}^{2}+\frac{J}{N/2}\left( 1-2\left\langle\rho_{j}^{2}\frac{1}{1+\frac{{\tau_{j}^{2}}/{\sigma_{\theta j}^{2}}}{T}} \right\rangle R\left( Y \right) \right)$	(S43)

again using (S8) to facilitate writing the sum of measurement error variance as an average over edges. If we then approximate an expectation of a product with a product of expectations, and expectation of a square root with square root of the expectation, we can use these to obtain the final result, an approximation to the expected value of the split-participants inter-edge correlation of FC-phenotype correlation:

$\frac{S_{\rho}^{2}}{S_{\rho}^{2}+\frac{1}{N/2}\left( 1-2\left\langle\rho_{j}^{2}\frac{1}{1+\frac{{\tau_{j}^{2}}/{\sigma_{\theta j}^{2}}}{T}} \right\rangle R(Y) \right)}$	(S44)

This can be interpreted as follows: This reliability measure has 3 fundamental inputs: (1) Inter-edge variability of true association, $S_{\rho}^{2}$, (2) reliability of FC-phenotype correlation depending on $\rho_{j}^{2}$, $\tau_{j}^{2}$, and $\sigma_{\theta j}^{2}$, and (3) reliability of phenotype $R(Y)$. These last 2 are jointly scaled by $\frac{1}{N/2}$, but the FC-phenotype correlation reliability also has a $\frac{1}{T}$ dependence embedded in an inter-edge average such that, all else equal, increased $T$ increases reliability. While the appearance of a negative term may be unexpected, it directly follows from the variance of Pearson’s correlation (S3) and as approximated by (S8): variance of Pearson’s correlation is maximal when true correlation is 0, and decreases with the absolute value of correlation; the reliability terms attenuate the correlation, and thus increase the sample variance. Note also that further simplification would be possible as $\left\langle\rho_{j}^{2} \right\rangle=S_{\rho}^{2}+\bar{\rho}^{2}$, i.e. the FC variability is also captured by the averaged squared true correlation, but this is complicated by the edgewise weighting involving ${\tau_{j}^{2}}/{\sigma_{\theta j}^{2}}$.

#### 1.3.1 **Empirical curve fitting for edgewise reliability**

The reliability result (S44) doesn’t make a simple prediction for the interplay of the $T$ and $N$ terms, as the influence of $T$ is embedded within an average over edges. However, if it is the case that $\left( 1+\frac{\tau_{j}^{2}}{\sigma_{\theta j}^{2}} \right)^{-1}$ varies little relative to $\rho_{j}^{2}$, then we might be able to fit observed correlations as a function of $N$ and $T$ like

$\frac{K_{0}}{K_{0}+\frac{1}{N/2}\left( 1-2K_{1}\frac{1}{1+{K_{2}}/T} \right)}$	(S45)

where we can interpret $K_{0}$ as the inter-edge variability of true association $S_{\rho}^{2}$, $K_{1}$ as the joint influence of FC-phenotype association variability and phenotype reliability $R\left( Y \right)$, and $K_{2}$ as the normalized FC measurement error relative to true FC variance.

### **Supplemental material**

#### **Supplementary Tables**

**Supplementary Table 1.** Summary of prediction accuracy analyses in the ABCD and HCP datasets. Table S1a shows the number of phenotypes in the prediction accuracy analyses across phenotypic domains. The “loose accuracy threshold” column shows the number of phenotypes whose prediction accuracies (Pearson’s r) were positive in at least 90% of all combinations of sample size *N* and scan time *T*. The “strict accuracy threshold” column shows the number of phenotypes whose prediction accuracies (Pearson’s r) were more than 0.1 when the full dataset was used (maximum *N* and *T*). The “Diminishing returns” column shows the number of phenotypes (among the subset of scores than passed the strict threshold) that exhibited diminishing returns in prediction accuracy with more than 20 minutes of scan time (relative to sample size). The “Exhibit log relationship” column shows the number of phenotypes (among the diminishing returns phenotypes) that displayed a possible logarithmic relationship between prediction accuracy and total scan duration. Table S1b shows the goodness-of-fit for the theoretical prediction accuracy models before and after randomizing run order. Goodness-of-fit was measured using the coefficient of determination (COD), which ranges from 0 to 1, which can be thought of fraction of variance explained. The COD was averaged across all phenotypes in the “loose accuracy threshold”, “strict accuracy threshold” and “exhibit log relationship”.

| **S1a. Number of phenotypes in the prediction accuracy analyses** | | | | | | | | | |
| --- | --- | --- | --- | --- | --- | --- | --- | --- | --- |
| **Dataset** | **Phenotypic Domain** | **Total** | **Loose threshold (at least 90% with r > 0)** | **Strict threshold (max r > 0.1)** | | **Diminishing returns at 20 mins** | | **Exhibit log relationship** | |
| ABCD | All | 37 | 33 (out of 37) | 23 (out of 37) | | 23 (out of 23) | | 17 (out of 23) | |
|  | Cognition | 17 | 17 (out of 17) | 15 (out of 17) | | 15 (out of 15) | | 13 (out of 15) | |
|  | Personality | 9 | 6 (out of 9) | 3 (out of 9) | | 3 (out of 3) | | 0 (out of 3) | |
|  | Mental Health | 11 | 10 (out of 11) | 5 (out of 11) | | 5 (out of 5) | | 4 (out of 5) | |
| HCP | All | 59 | 42 (out of 59) | 28 (out of 59) | | 25 (out of 28) | | 19 (out of 25) | |
|  | Cognition | 20 | 18 (out of 20) | 15 (out of 20) | | 14 (out of 15) | | 12 (out of 14) | |
|  | Personality | 6 | 5 (out of 6) | 4 (out of 6) | | 4 (out of 4) | | 4 (out of 4) | |
|  | Emotion | 13 | 6 (out of 13) | 1 (out of 13) | | 0 (out of 1) | | 1 (out of 1) | |
|  | Physical | 9 | 7 (out of 9) | 5 (out of 9) | | 5 (out of 5) | | 1 (out of 5) | |
|  | Well-being | 11 | 6 (out of 11) | 3 (out of 11) | | 2 (out of 3) | | 1 (out of 2) | |
| **S1b. Average goodness-of-fit (COD or R^2^) over each set of phenotypes for prediction models before and after randomizing run orders** | | | | | | | | | |
|  | | | | **20 min** | **58 min** | | **20 min (random)** | | **58 min (random)** |
| ABCD (Loose accuracy threshold: 33 phenotypes) | | | |  |  | |  | |  |
| Theoretical | | | | 0.763 | N.A | | 0.844 | | N.A |
| ABCD (Strict accuracy threshold: 23 phenotypes) | | | |  |  | |  | |  |
| Theoretical | | | | 0.834 | N.A | | 0.885 | | N.A |
| ABCD (Exhibit log relationship: 17 phenotypes) | | | |  |  | |  | |  |
| Theoretical | | | | 0.894 | N.A | | 0.940 | | N.A |
| HCP (Loose accuracy threshold: 42 phenotypes) | | | |  |  | |  | |  |
| Theoretical | | | | 0.724 | 0.728 | | 0.854 | | 0.880 |
| HCP (Strict accuracy threshold: 28 phenotypes) | | | |  |  | |  | |  |
| Theoretical | | | | 0.818 | 0.836 | | 0.918 | | 0.926 |
| HCP (Exhibit log relationship: 19 phenotypes) | | | |  |  | |  | |  |
| Theoretical | | | | 0.890 | 0.888 | | 0.945 | | 0.947 |

##### **Supplementary Table 2.** Summary of prediction accuracy analyses for all datasets. The “strict accuracy threshold” column shows the number of phenotypes whose prediction accuracies (Pearson’s r) were more than 0.1 when the full dataset was used (maximum *N* and *T*). The “Adherence to theoretical model” column shows the number of phenotypes which passed the strict accuracy threshold and showed a good fit to the theoretical models after visual assessment. “Average COD of model fit” refers to the goodness-of-fit of the theoretical model averaged over the phenotypes that showed adherence to the theoretical model. We note that many phenotypes overlap between ABCD (rest), ABCD (MID), ABCD (NBACK), ABCD (SST), so in the case of the ABCD, there were in total 23 unique phenotypes that showed adherence to the theoretical model, yielding 23 + 19 + 14 + 7 + 7 + 6 = 76 unique phenotypes used for generating Fig. 6a.

| **Dataset** | **Total number of phenotypes** | **Strict accuracy** **threshold (max r > 0.1)** | **Adherence to theoretical model** | **Average COD of model fit** |
| --- | --- | --- | --- | --- |
| ABCD (rest) | 37 | 23 (out of 37) | 17 (out of 23) | 0.894 |
| HCP | 59 | 28 (out of 59) | 19 (out of 28) | 0.888 |
| SINGER | 19 | 15 (out of 19) | 14 (out of 15) | 0.926 |
| TCP | 19 | 10 (out of 19) | 7 (out of 10) | 0.818 |
| MDD | 20 | 11 (out of 20) | 7 (out of 11) | 0.844 |
| ADNI | 6 | 6 (out of 6) | 6 (out of 6) | 0.920 |
| ABCD (MID) | 37 | 21 (out of 37) | 16 (out of 21) | 0.921 |
| ABCD (NBACK) | 37 | 22 (out of 37) | 19 (out of 22) | 0.884 |
| ABCD (SST) | 37 | 22 (out of 37) | 18 (out of 22) | 0.872 |
| **Control Analyses** |  |  |  |  |
| ABCD (subcortical) | 37 | 18 (out of 37) | 14 (out of 18) | 0.868 |
| HCP (subcortical) | 59 | 21 (out of 59) | 13 (out of 21) | 0.860 |
| ABCD (1000 parcels) | 37 | 24 (out of 37) | 18 (out of 24) | 0.890 |
| HCP (1000 parcels) | 59 | 30 (out of 59) | 24 (out of 30) | 0.858 |
| HCP (mix days) | 59 | 21 (out of 59) | 16 (out of 21) | 0.847 |

**Supplementary Table 3.1** Phenotypic measures in the ABCD dataset.

|  | **Description** | **ABCD field** | **ABCD file** | **Category** |
| --- | --- | --- | --- | --- |
| 1 | Anxious Depressed | cbcl_scr_syn_anxdep_r | abcd_cbcls01.txt | Mental Health |
| 2 | Withdrawn Depressed | cbcl_scr_syn_withdep_r | abcd_cbcls01.txt | Mental Health |
| 3 | Somatic Complaints | cbcl_scr_syn_somatic_r | abcd_cbcls01.txt | Mental Health |
| 4 | Social Problems | cbcl_scr_syn_social_r | abcd_cbcls01.txt | Mental Health |
| 5 | Thought Problems | cbcl_scr_syn_thought_r | abcd_cbcls01.txt | Mental Health |
| 6 | Attention Problems | cbcl_scr_syn_attention_r | abcd_cbcls01.txt | Mental Health |
| 7 | Rule-breaking Behavior | cbcl_scr_syn_rulebreak_r | abcd_cbcls01.txt | Mental Health |
| 8 | Aggressive Behavior | cbcl_scr_syn_aggressive_r | abcd_cbcls01.txt | Mental Health |
| 9 | Vocabulary | nihtbx_picvocab_uncorrected | abcd_tbss01.txt | Cognition |
| 10 | Attention | nihtbx_flanker_uncorrected | abcd_tbss01.txt | Cognition |
| 11 | Working Memory | nihtbx_list_uncorrected | abcd_tbss01.txt | Cognition |
| 12 | Executive Function | nihtbx_cardsort_uncorrected | abcd_tbss01.txt | Cognition |
| 13 | Processing Speed | nihtbx_pattern_uncorrected | abcd_tbss01.txt | Cognition |
| 14 | Episodic Memory | nihtbx_picture_uncorrected | abcd_tbss01.txt | Cognition |
| 15 | Reading | nihtbx_reading_uncorrected | abcd_tbss01.txt | Cognition |
| 16 | Fluid Cognition | nihtbx_fluidcomp_uncorrected | abcd_tbss01.txt | Cognition |
| 17 | Crystallized Cognition | nihtbx_cryst_uncorrected | abcd_tbss01.txt | Cognition |
| 18 | Overall Cognition | nihtbx_totalcomp_uncorrected | abcd_tbss01.txt | Cognition |
| 19 | Negative Urgency | upps_y_ss_negative_urgency | abcd_mhy02.txt | Personality |
| 20 | Lack of Planning | upps_y_ss_lack_of_planning | abcd_mhy02.txt | Personality |
| 21 | Sensation Seeking | upps_y_ss_sensation_seeking | abcd_mhy02.txt | Personality |
| 22 | Positive Urgency | upps_y_ss_positive_urgency | abcd_mhy02.txt | Personality |
| 23 | Lack Perseverance | upps_y_lack_of_perseverance | abcd_mhy02.txt | Personality |
| 24 | Behavioral Inhibition | bis_y_ss_bis_sum | abcd_mhy02.txt | Personality |
| 25 | Reward Responsiveness | bis_y_ss_bas_rr | abcd_mhy02.txt | Personality |
| 26 | Drive | bis_y_ss_bas_drive | abcd_mhy02.txt | Personality |
| 27 | Fun Seeking | bis_y_ss_bas_fs | abcd_mhy02.txt | Personality |
| 28 | Total Psychosis Symptoms | pps_y_ss_number | abcd_mhy02.txt | Mental Health |
| 29 | Psychosis Severity | pps_y_ss_severity_score | abcd_mhy02.txt | Mental Health |
| 30 | Mania | pgbi_p_ss_score | abcd_mhp02.txt | Mental Health |
| 31 | Short Delay Recall | pea_ravlt_sd_trial_vi_tc | abcd_ps01.txt | Cognition |
| 32 | Long Delay Recall | pea_ravlt_ld_trial_vii_tc | abcd_ps01.txt | Cognition |
| 33 | Fluid Intelligence | pea_wiscv_trs | abcd_ps01.txt | Cognition |
| 34 | Visuospatial Accuracy | lmt_scr_perc_correct | lmtp201.txt | Cognition |
| 35 | Visuospatial Reaction Time | lmt_scr_rt_correct | lmtp201.txt | Cognition |
| 36 | Visuospatial Efficiency | lmt_scr_efficiency | lmtp201.txt | Cognition |
| 37 | Cognitive factor score | Obtained from PCA of previous 36 measures | N.A. | Cognition |

**Supplementary Table 3.2** Phenotypic measures in the HCP dataset.

|  | **Description** | **HCP field** | **Category** |
| --- | --- | --- | --- |
| 1 | Visual Episodic Memory | PicSeq_Unadj | Cognition |
| 2 | Cognitive Flexibility (DCCS) | CardSort_Unadj | Cognition |
| 3 | Inhibition (Flanker Task) | Flanker_Unadj | Cognition |
| 4 | Fluid Intelligence (PMAT) | PMAT24_A_CR | Cognition |
| 5 | Reading (Pronunciation) | ReadEng_Unadj | Cognition |
| 6 | Vocabulary (Picture Matching) | PicVocab_Unadj | Cognition |
| 7 | Processing Speed | ProcSpeed_Unadj | Cognition |
| 8 | Delay Discounting | DDic_AUC_40K | Personality |
| 9 | Spatial Orientation | VSPLOT_TC | Cognition |
| 10 | Sustained Attention – Sens. | SCPT_SEN | Cognition |
| 11 | Sustained Attention – Spec. | SCPT_SPEC | Cognition |
| 12 | Verbal Episodic Memory | IWRD_TOT | Cognition |
| 13 | Working Memory (List Sorting) | ListSort_Unadj | Cognition |
| 14 | Cognitive Status (MMSE) | MMSE_Score | Cognition |
| 15 | Sleep Quality (PSQI) | PSQI_Score | Physical |
| 16 | Walking Endurance | Endurance_Unadj | Physical |
| 17 | Walking Speed | GaitSpeed_Unadj | Physical |
| 18 | Manual Dexterity | Dexterity_Unadj | Physical |
| 19 | Grip Strength | Strength_Unadj | Physical |
| 20 | Odor Identification | Odor_Unadj | Physical |
| 21 | Pain Interference Survey | PainInterf_Tscore | Physical |
| 22 | Taste Intensity | Taste_Unadj | Physical |
| 23 | Contrast Sensitivity | Mars_Final | Physical |
| 24 | Emotional Face Matching | Emotion_Task_Face_Acc | Emotion |
| 25 | Arithmetic | Language_Task_Math_Avg_Difficulty_Level | Cognition |
| 26 | Story Comprehension | Language_Task_Story_Avg_Difficulty_Level | Cognition |
| 27 | Relational Processing | Relational_Task_Acc | Cognition |
| 28 | Social Cognition – Random | Social_Task_Perc_Random | Cognition |
| 29 | Social Cognition – Interaction | Social_Task_Perc_TOM | Cognition |
| 30 | Working Memory (N-back) | WM_Task_Acc | Cognition |
| 31 | Agreeableness (NEO) | NEOFAC_A | Personality |
| 32 | Openness (NEO) | NEOFAC_O | Personality |
| 33 | Conscientiousness (NEO) | NEOFAC_C | Personality |
| 34 | Neuroticism (NEO) | NEOFAC_N | Personality |
| 35 | Extraversion (NEO) | NEOFAC_E | Personality |
| 36 | Emot. Recog. – Total | ER40_CR | Emotion |
| 37 | Emot. Recog. – Angry | ER40ANG | Emotion |
| 38 | Emot. Recog. – Fear | ER40FEAR | Emotion |
| 39 | Emot. Recog. – Happy | ER40HAP | Emotion |
| 40 | Emot. Recog. - Neutral | ER40NOE | Emotion |
| 41 | Emot. Recog. – Sad | ER40SAD | Emotion |
| 42 | Anger – Affect | AngAffect_Unadj | Emotion |
| 43 | Anger – Hostility | AngHostil_Unadj | Emotion |
| 44 | Anger – Aggression | AngAggr_Unadj | Emotion |
| 45 | Fear – Affect | FearAffect_Unadj | Emotion |
| 46 | Fear – Somatic Arousal | FearSomat_Unadj | Emotion |
| 47 | Sadness | Sadness_Unadj | Emotion |
| 48 | Life Satisfaction | LifeSatisf_Unadj | Well-being |
| 49 | Meaning & Purpose | MeanPurp_Unadj | Well-being |
| 50 | Positive Affect | PosAffect_Unadj | Well-being |
| 51 | Friendship | Friendship_Unadj | Well-being |
| 52 | Loneliness | Loneliness_Unadj | Well-being |
| 53 | Perceived Hostility | PercHostil_Unadj | Well-being |
| 54 | Perceived Rejection | PercReject_Unadj | Well-being |
| 55 | Emotional Support | EmotSupp_Unadj | Well-being |
| 56 | Instrument Support | InstruSupp_Unadj | Well-being |
| 57 | Perceived Stress | PercStress_Unadj | Well-being |
| 58 | Self-Efficacy | SelfEff_Unadj | Well-being |
| 59 | Cognitive factor score | Obtained from PCA of previous 58 measures | Cognition |

**Supplementary Table 3.3** Phenotypic measures in the SINGER dataset.

|  | **Description** | **SINGER field** | **Category** |
| --- | --- | --- | --- |
| 1 | Age | SC_age | Physical |
| 2 | Montreal Cognitive Assessment | SC_moca_TOTAL | Cognition |
| 3 | Years of Education | BL_Demo6b_eduyr | Cognition |
| 4 | Body Mass Index | BL_bmi | Physical |
| 5 | Grip Strength (Left) | BL_leftgrip_avg | Physical |
| 6 | Grip Strength (Right) | BL_rightgrip_avg | Physical |
| 7 | Mini-Mental State Examination | BL_MMSETotalScore | Cognition |
| 8 | Visual Paired Associates | BL_NTBVPAImmediateTotalScore | Cognition |
| 9 | Logical Memory | BL_NTBLogicalMemoryTotalScore | Cognition |
| 10 | Rey Auditory Visual Learning Test | BL_NTBRAVLTTotalScore | Cognition |
| 11 | Digit Span | BL_NTBDSTotalScore | Cognition |
| 12 | Category Fluency Test | BL_NTBCFTTotalScore | Cognition |
| 13 | Delayed visual paired associates | BL_NTBVPADelayTotalScore | Cognition |
| 14 | Delayed Logical Memory | BL_NTBLMDelayTotalScore | Cognition |
| 15 | Delayed Rey Auditory Visual Learning Test | BL_RAVLTdelayed_TOTAL | Cognition |
| 16 | Rey Auditory Visual Learning Test (Delay Recognition) | BL_NTBRAVLTDelayRecogTotalScore | Cognition |
| 17 | Trail Making Test Part A | BL_TMTPartASec | Cognition |
| 18 | Trail Making Test Part B | BL_TMTPartBSec | Cognition |
| 19 | Letter Digit Substitution | BL_LDSTTotalScore | Cognition |

**Supplementary Table 3.4** Phenotypic measures in the TCP dataset.

|  | **Description** | **TCP field** | **Category** |
| --- | --- | --- | --- |
| 1 | Age | age | Physical |
| 2 | Columbia-Suicide Severity Rating Scale Ideation | cssrs_isi | Mental Health |
| 3 | Montgomery-Ãsberg Depression Rating Scale | madrstot | Mental Health |
| 4 | Positive and Negative Syndrome Scale (General Psychopathology) | panss_gen_clean | Mental Health |
| 5 | Positive and Negative Syndrome Scale (Negative Symptoms) | panss_neg_clean | Mental Health |
| 6 | Positive and Negative Syndrome Scale (Positive Symptoms) | panss_pos_clean | Mental Health |
| 7 | Young Mania Rating Scale | ymrs_tot | Mental Health |
| 8 | Depression Anxiety Stress Scale (Anxiety) | dass_anx_sc | Mental Health |
| 9 | Depression Anxiety Stress Scale (Depression) | dass_depr_sc | Mental Health |
| 10 | Depression Anxiety Stress Scale (Stress) | dass_stress_sc | Mental Health |
| 11 | Depression Anxiety Stress Scale (Total) | dass_total | Mental Health |
| 12 | Marder Factor Score (Anxiety / Depression) | panss_marder_AnxDep | Mental Health |
| 13 | Marder Factor Score (Cognitive / Disorganisation) | panss_marder_CogDis | Mental Health |
| 14 | Marder Factor Score (Negative Symptoms) | panss_marder_Neg | Mental Health |
| 15 | Marder Factor Score (Positive Symptoms) | panss_marder_Pos | Mental Health |
| 16 | Marder Factor Score (Uncontrolled Hostility / Excitement) | panss_marder_UHE | Mental Health |
| 17 | Positive and Negative Syndrome Scale (Total) | panss_tot | Mental Health |
| 18 | Perceived Stress | pss_totalscore | Mental Health |
| 19 | Body Mass Index | bmi | Physical |

**Supplementary Table 3.5** Phenotypic measures in the MDD dataset.

|  | **Description** | **MDD field** | **Category** |
| --- | --- | --- | --- |
| 1 | Age | Age | Physical |
| 2 | Hamilton Depression Rating Scale (Total) | HAMD_baseline_total | Mental Health |
| 3 | Hamilton Anxiety Rating Scale (Total) | HAMA_baseline_total | Mental Health |
| 4 | Depressed Mood | HAMD01_baseline | Mental Health |
| 5 | Guilt | HAMD02_baseline | Mental Health |
| 6 | Suicide | HAMD03_baseline | Mental Health |
| 7 | Early Insomnia | HAMD04_baseline | Mental Health |
| 8 | Middle Insomnia | HAMD05_baseline | Mental Health |
| 9 | Late Insomnia | HAMD06_baseline | Mental Health |
| 10 | Work and Interests | HAMD07_baseline | Mental Health |
| 11 | Retardation | HAMD08_baseline | Mental Health |
| 12 | Agitation | HAMD09_baseline | Mental Health |
| 13 | Anxiety (Psychic) | HAMD10_baseline | Mental Health |
| 14 | Anxiety (Somatic) | HAMD11_baseline | Mental Health |
| 15 | Somatic Symptoms (Gastrointestinal) | HAMD12_baseline | Mental Health |
| 16 | Somatic Symptoms (General) | HAMD13_baseline | Mental Health |
| 17 | Genital Symptoms | HAMD14_baseline | Mental Health |
| 18 | Hypochondriasis | HAMD15_baseline | Mental Health |
| 19 | Weight Loss | HAMD16_baseline | Physical |
| 20 | Insight | HAMD17_baseline | Mental Health |

**Supplementary Table 3.6** Phenotypic measures in the ADNI dataset.

|  | **Description** | **ADNI field** | **Category** |
| --- | --- | --- | --- |
| 1 | Age | age | Physical |
| 2 | Body Mass Index | body_mass_index | Physical |
| 3 | Delayed Logical Memory | Logical_memory_delayed | Cognition |
| 4 | Mini-Mental State Examination | MMSE | Cognition |
| 5 | Beta Amyloid level (Default A network) | amyloid_defaultA | PET |
| 6 | Beta Amyloid level (Default B network) | amyloid_defaultB | PET |

**Supplementary Table 4.1** Demographics of each dataset.

|  | **N** | **Age (Years)** | **Sex** | **Racial Groups** |
| --- | --- | --- | --- | --- |
| HCP | 792 | 22 - 36 (mean = 28.6) | 371M / 421F | White (N = 612),  Black / African Am. (N = 94),  Am. Indian / Alaskan Nat. (N = 1),  Asian / Nat. Hawaiian / Other Pacific Is. (N = 53),  More than one (N = 21),  Unknown or Not Reported (N = 11) |
| ABCD-rest | 2565 | 9.00 - 10.9 (mean = 10.0) | 1251M / 1314F | White (N = 1443), Black (N = 277), Hispanic (N = 501),  Asian (N = 66),  Other (N = 273),  Not declared (N = 5) |
| ABCD-task | 2262 | 9.00 - 10.91 (mean = 10.01) | 1030M / 1232F | White (N = 1335), Black (N = 201), Hispanic (N = 425),  Asian (N = 60),  Other (N = 236),  Not declared (N = 5) |
| SINGER | 642 | 60 - 80 (mean = 68.8) | 309M / 333F | Chinese (N = 625), Malay (N = 2), Indian (N = 12), Other (N = 3) |
| TCP | 194 | 18.08 – 65.0 (mean = 33.4) | 81M / 110F / 3 SD | White (N = 116), Black or African American (N = 30), Asian (N = 29), More than one (N = 13), Other (N = 6) |
| MDD | 287 | 16 – 64 (mean = 32.3) | 101M / 186F | Chinese (N = 287) |
| ADNI | 586 | 50.8 – 97.5 (mean = 74.4) | 278M / 308F | White (N = 510), Black (N = 43), Asian (N = 15), American Indian (N = 1), More than one (N = 12), Unknown (N = 5) |

**Supplementary Table 4.2** Diagnostic distributions for each dataset

|  | Diagnostic distributions |
| --- | --- |
| HCP | Healthy Controls |
| ABCD | Healthy Controls |
| SINGER | Elderly at risk for vascular cognitive impairment |
| TCP | Control (N = 76), MDD (N = 31), PTSD (N = 14), GAD (N = 12), Dysthymia (N = 9), Social Anxiety Disorder (N = 8), SUD (N = 8), BPD I (N = 6), BPD II (N = 6), Other Anxiety Disorder (N = 5), Other Mood Disorder (N = 4), Schizophrenia (N = 4), Schizoaffective Disorder (N = 4), ADHD (N = 3), Eating Disorder (N = 2), OCD (N = 2) |
| MDD | Major Depressive Disorder (N = 287) |
| ADNI | Control (N = 334), Mild Cognitive Impairment (N = 184), AD dementia (N = 68) |

**Supplementary Table 4.3** Acquisition information for each dataset

|  | Acquisition | Voxel size/mm^3^ | TR/s | Atlas Space |
| --- | --- | --- | --- | --- |
| HCP | Single-echo Multi-band Custom Skyra | 2.0 | 0.72 | fsLR |
| ABCD | Single-echo Multi-band on GE & Siemens scanners (more information in Table S6.1) | 2.4 | 0.8 | fsaverage |
| SINGER | Multi-echo multi-band Prisma Fit | 3.0 | 1.0 | fsaverage |
| TCP | Single-echo multi-band on two Prisma scanners | 2.0 | 0.8 | MNI152 |
| MDD | Single-echo single-band on five Prisma scanners | 3.0 | 3.0 | fsaverage |
| ADNI | Single-echo multi-band (N = 78) & single-echo single-band (N = 508) on GE, Philips & Siemens scanners (more information in Table S6.2) | 2.5 & ~3.4 | 0.6 & 3 | fsaverage |

**Supplementary Table 5.1** Site clusters for ABCD resting-state fMRI

| **ABCD Site** | **Make** | **Model** | **N** | **Site-cluster** |
| --- | --- | --- | --- | --- |
| 2 | Siemens | Prisma fit | 97 | A |
| 6 | Siemens | Prisma fit | 67 | A |
| 12 | Siemens | Prisma fit | 79 | A |
| 4 | GE | Discovery MR750 | 315 | B |
| 3 | Siemens | Prisma | 183 | C |
| 9 | Siemens | Prisma fit | 51 | C |
| 5 | Siemens | Prisma fit | 97 | D |
| 20 | Siemens | Prisma/Prisma fit | 139 | D |
| 10 | GE | Discovery MR750 | 218 | E |
| 22 | GE | Discovery MR750 | 14 | E |
| 7 | Siemens | Prisma fit | 90 | F |
| 14 | Siemens | Prisma/Prisma fit | 150 | F |
| 13 | GE | Discovery MR750 | 245 | G |
| 11 | Siemens | Prisma | 100 | H |
| 15 | Siemens | Prisma fit | 69 | H |
| 21 | Siemens | Prisma fit/Prisma | 97 | H |
| 16 | Siemens | Prisma | 327 | I |
| 8 | GE | Discovery MR750 | 106 | J |
| 18 | GE | Discovery MR750 | 121 | J |

**Supplementary Table 5.2** Site clusters for ADNI

| **ADNI site** | **Make** | **Model** | **N** | **Site-cluster** |
| --- | --- | --- | --- | --- |
| 58 | Siemens | Prisma fit | 33 | A |
| 2 | Siemens | Prisma | 27 | B |
| 23 | Siemens | Prisma fit | 4 | B |
| 28 | Siemens | Prisma fit | 25 | C |
| 3 | GE | Discovery MR750 | 4 | C |
| 33 | Siemens | Prisma fit | 15 | D |
| 55 | Siemens | Verio | 14 | D |
| 59 | Siemens | Prisma fit | 18 | E |
| 39 | GE | Signa premier | 11 | E |
| 18 | Siemens | Prisma fit | 10 | F |
| 16 | GE | Discovery MR750 | 19 | F |
| 20 | GE | Discovery MR750 | 22 | G |
| 52 | GE | Signa premier | 7 | G |
| 52 | GE | Discovery MR750w | 15 | H |
| 8 | Siemens | Prisma fit | 12 | H |
| 7 | Philips | Achieva | 1 | H |
| 9 | Siemens | Prisma | 1 | H |
| 50 | Philips | Achieva | 20 | I |
| 46 | GE | Discovery MR750 | 9 | I |
| 47 | GE | Discovery MR750 | 23 | J |
| 1 | Philips | Intera | 5 | J |
| 10 | Siemens | Biograph mMR | 1 | J |
| 25 | Siemens | Prisma fit | 18 | K |
| 40 | GE | Discovery MR750 | 11 | K |
| 4 | Philips | Ingenia | 23 | L |
| 13 | Philips | Achieva | 3 | L |
| 16 | GE | Signa UHP | 3 | L |
| 27 | Siemens | Prisma | 22 | M |
| 9 | Philips | Achieva | 7 | M |
| 49 | GE | Discovery MR750 | 17 | N |
| 1 | Siemens | Prisma fit | 11 | N |
| 21 | Philips | Achieva | 1 | N |
| 11 | Siemens | Verio | 9 | O |
| 15 | Siemens | Prisma fit | 9 | O |
| 21 | GE | Discovery MR750w | 9 | O |
| 22 | Philips | Achieva | 2 | O |
| 38 | Siemens | Prisma | 11 | P |
| 62 | Philips | Achieva | 8 | P |
| 14 | Philips | Achieva | 8 | P |
| 17 | Siemens | Prisma fit | 2 | P |
| 40 | Philips | Achieva | 5 | Q |
| 41 | Philips | Achieva | 5 | Q |
| 43 | Siemens | Skyra fit | 8 | Q |
| 26 | Siemens | Skyra | 9 | Q |
| 29 | Siemens | Prisma fit | 2 | Q |
| 43 | Siemens | Verio | 8 | R |
| 44 | Siemens | TrioTim | 5 | R |
| 45 | Philips | Ingenia | 4 | R |
| 5 | Siemens | Prisma | 5 | R |
| 60 | Philips | Ingenia | 2 | R |
| 61 | Philips | Ingenia | 5 | R |
| 13 | Philips | Achieva | 5 | S |
| 19 | Siemens | Prisma fit | 5 | S |
| 30 | Philips | Intera | 5 | S |
| 5 | Philips | Singa HDxt | 5 | S |
| 50 | Philips | Achieva | 6 | S |
| 63 | Siemens | Prisma | 3 | S |
| 17 | Siemens | TrioTim | 2 | T |
| 30 | Siemens | Prisma | 3 | T |
| 31 | Philips | Ingenia Elition X | 1 | T |
| 31 | Philips | Intera | 1 | T |
| 32 | GE | Discovery MR750 | 2 | T |
| 34 | Siemens | TrioTim | 1 | T |
| 37 | Philips | Singa HDxt | 3 | T |
| 37 | Siemens | Verio | 3 | T |
| 39 | GE | Discovery MR750 | 1 | T |
| 41 | Philips | Ingenia Elition X | 1 | T |
| 51 | Philips | Ingenia | 2 | T |
| 59 | Philips | Achieva | 3 | T |
| 6 | GE | Discovery MR750w | 4 | T |
| 7 | GE | Signa premier | 2 | T |

**Supplementary Table 5.3** Site clusters for ABCD task-fMRI

| **ABCD Site** | **Make** | **Model** | **N** | **Site-cluster** |
| --- | --- | --- | --- | --- |
| 2 | Siemens | Prisma fit | 129 | A |
| 6 | Siemens | Prisma fit | 150 | A |
| 12 | Siemens | Prisma fit | 116 | A |
| 4 | GE | Discovery MR750 | 170 | B |
| 3 | GE | Discovery MR750 | 160 | C |
| 9 | Siemens | Prisma | 66 | C |
| 5 | Siemens | Prisma fit | 72 | D |
| 20 | Siemens | Prisma fit | 103 | D |
| 10 | Siemens | Prisma/Prisma fit | 161 | E |
| 22 | GE | Discovery MR750 | 13 | E |
| 7 | Siemens | Prisma fit | 71 | F |
| 14 | Siemens | Prisma/Prisma fit | 165 | F |
| 13 | GE | Discovery MR750 | 192 | G |
| 11 | Siemens | Prisma | 71 | H |
| 15 | Siemens | Prisma fit | 37 | H |
| 12 | Siemens | Prisma fit/Prisma | 93 | H |
| 16 | Siemens | Prisma | 337 | I |
| 8 | GE | Discovery MR750 | 74 | J |
| 18 | GE | Discovery MR750 | 82 | J |

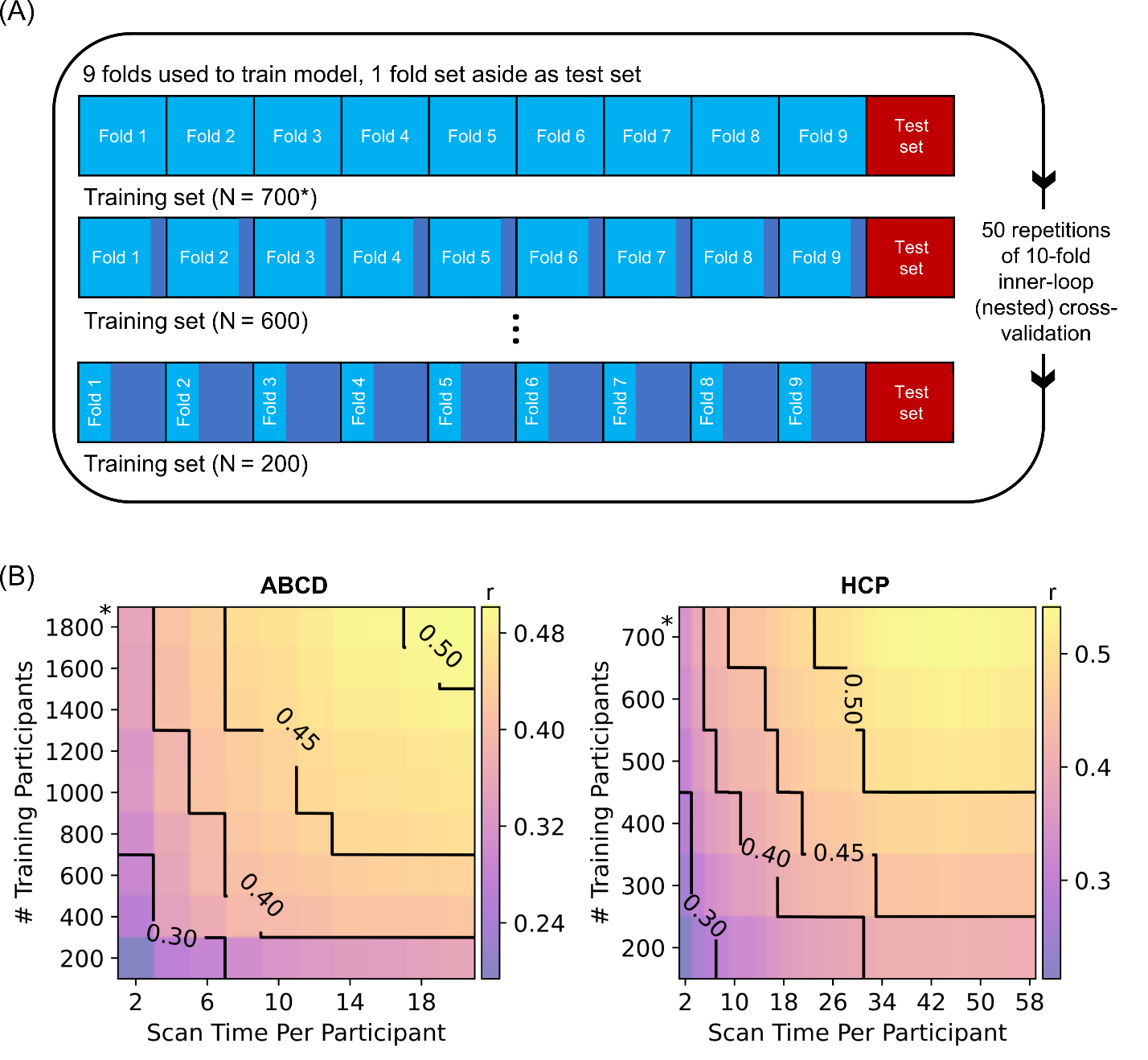

#### **Supplementary Figures**

##### **Supplementary Fig. 1 |** Contour plot of prediction accuracy (Pearson’s correlation) of the cognitive factor score as a function of the scan time T used to generate the functional connectivity matrix, and the number of training participants N used to train the predictive model in the Human Connectome Project (HCP) dataset. Increasing training participants and scan time both improved prediction performance. The * indicates that all available participants were used, therefore the sample size will be close to, but not exactly the number shown.

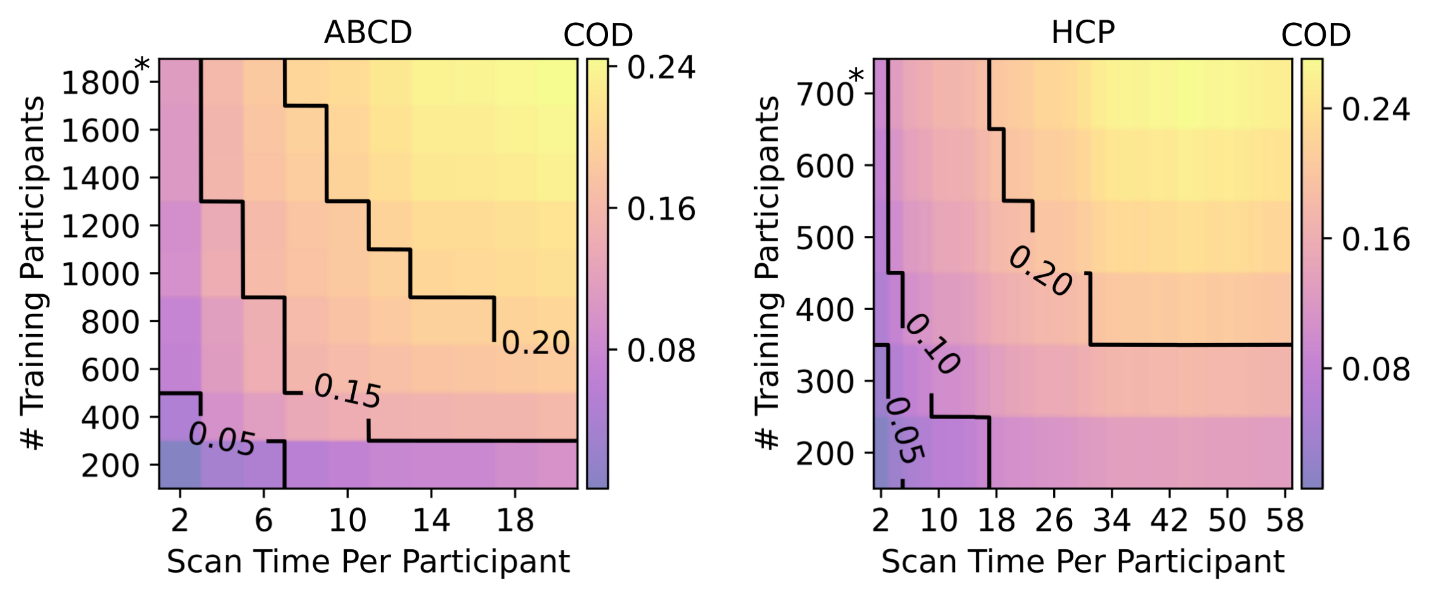

**Supplementary Fig. 2 |** Same as Fig. 1a except prediction accuracy was calculated with Coefficient of Determination (COD or R^2^) instead of Pearson’s correlation. Contour plot of prediction accuracy (COD) of the cognitive component score as a function of the scan time used to generate the functional connectivity matrix (x-axis), and the number of training participants used to train the predictive model (y-axis) in the ABCD and HCP datasets. Increasing training participants and scan time both led to increases in prediction performance. The * in both figures indicates that all available participants were used, therefore the sample size will be close to, but not exactly the number shown.

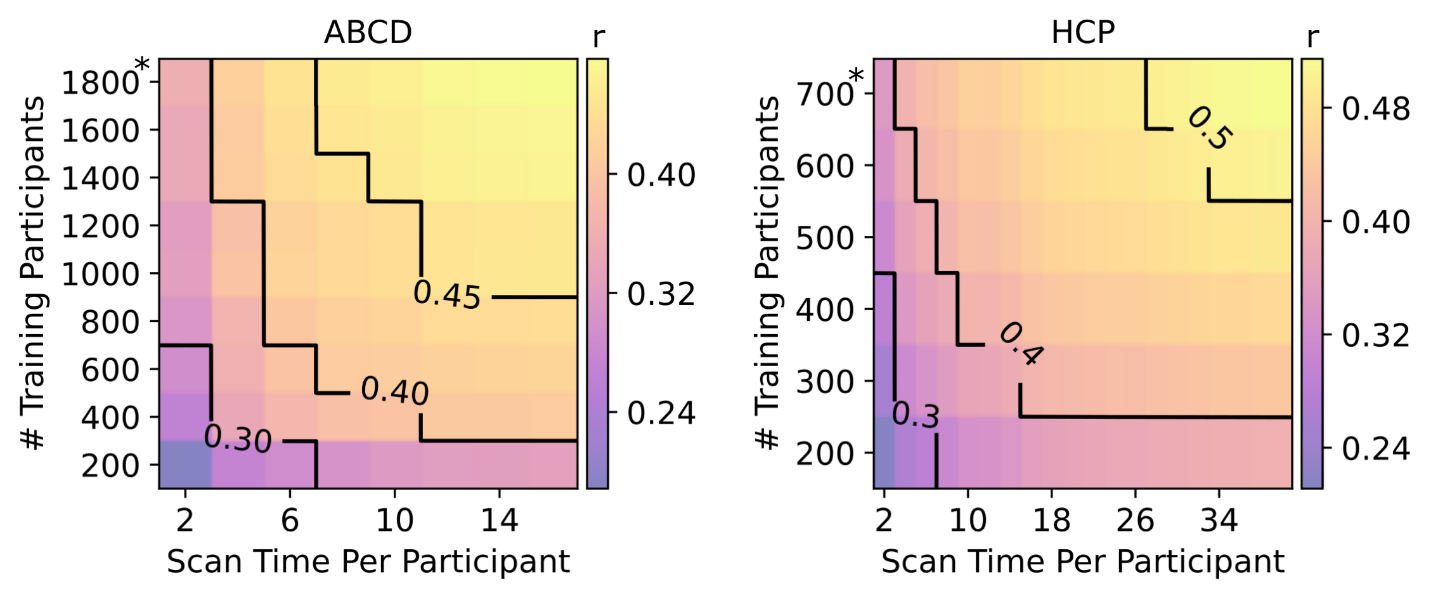

##### **Supplementary Fig. 3 |** Same as Fig. 1a except functional connectivity matrices with constructed with first *T* minutes of uncensored data. Contour plot of prediction accuracy (Pearson’s correlation) of the cognitive component score as a function of the scan time used to generate the functional connectivity matrix (x-axis), and the number of training participants used to train the predictive model (y-axis) in the ABCD and HCP datasets. Increasing training participants and scan time both led to increases in prediction performance. The * in both figures indicates that all available participants were used, therefore the sample size will be close to, but not exactly the number shown.

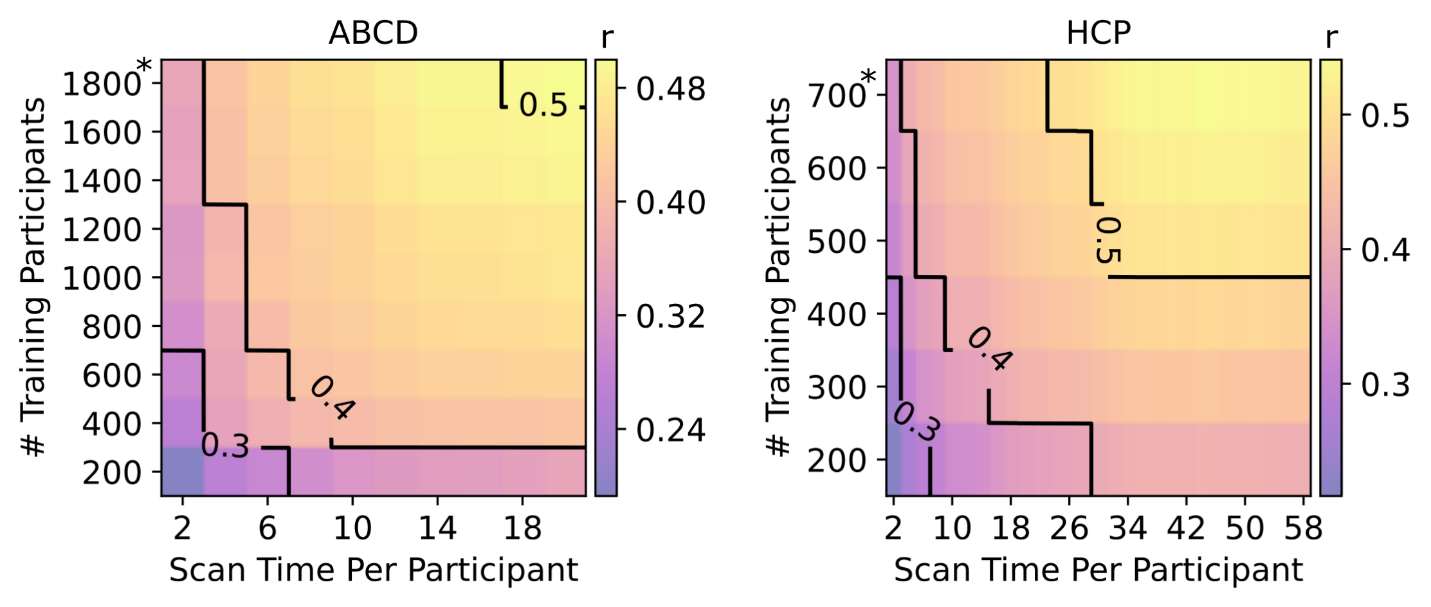

##### **Supplementary Fig. 4 |** Same as Fig. 1a except censored frames were not excluded when computing the functional connectivity matrices. Contour plot of prediction accuracy (Pearson’s correlation) of the cognitive component score as a function of the scan time used to generate the functional connectivity matrix (x-axis), and the number of training participants used to train the predictive model (y-axis) in the ABCD and HCP datasets. Increasing training participants and scan time both led to increases in prediction performance. The * in both figures indicates that all available participants were used, therefore the sample size will be close to, but not exactly the number shown.

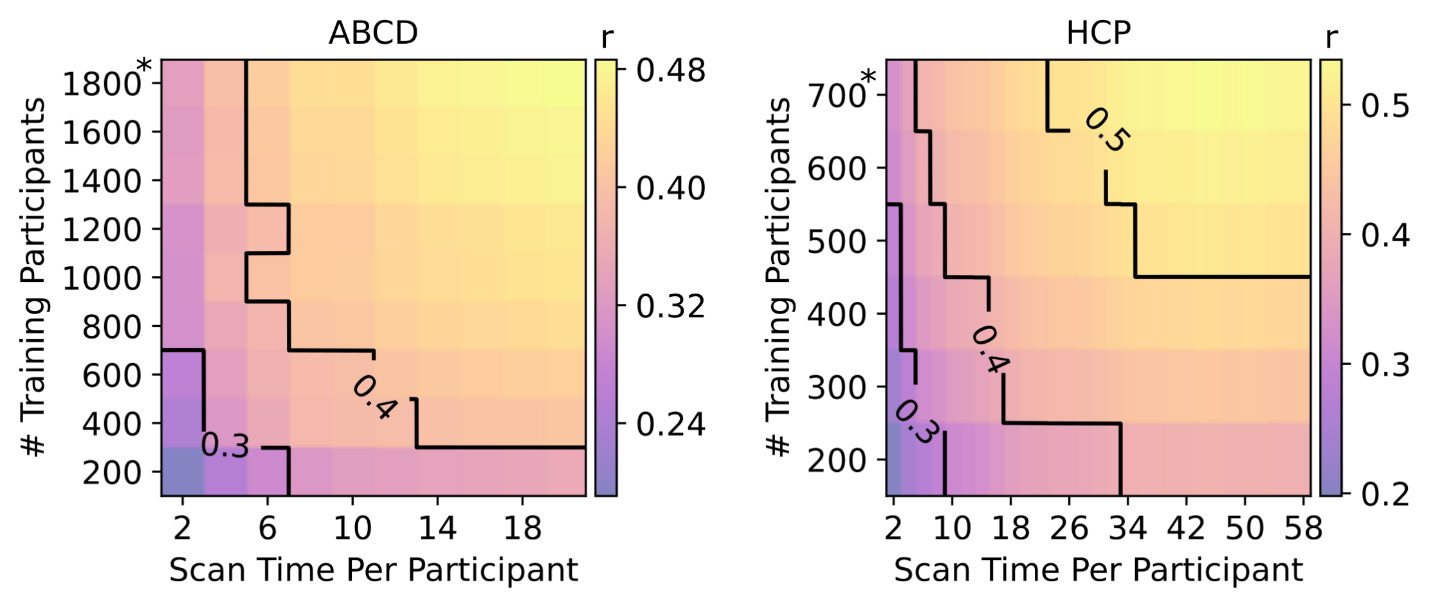

##### **Supplementary Fig. 5 |** Same as Fig. 1a except linear ridge regression was used as the prediction algorithm instead of kernel ridge regression. Contour plot of prediction accuracy (Pearson’s correlation) of the cognitive component score as a function of the scan time used to generate the functional connectivity matrix (x-axis), and the number of training participants used to train the predictive model (y-axis) in the ABCD and HCP datasets. Increasing training participants and scan time both led to increases in prediction performance. The * in both figures indicates that all available participants were used, therefore the sample size will be close to, but not exactly the number shown.

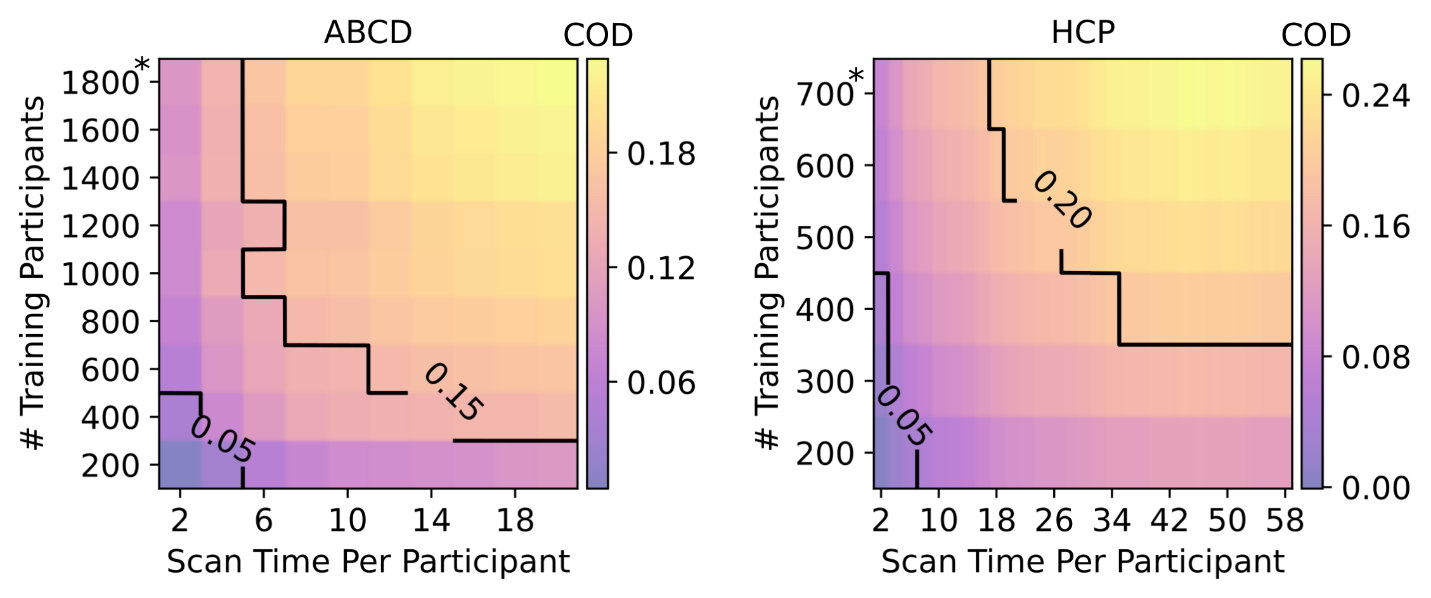

##### **Supplementary Fig. 6 |** Same as Fig. 1a except linear ridge regression was used as the prediction algorithm instead of kernel ridge regression and prediction accuracy was calculated with Coefficient of Determination (COD or R^2^) instead of Pearson’s correlation. Contour plot of prediction accuracy (COD) of the cognitive component score as a function of the scan time used to generate the functional connectivity matrix (x-axis), and the number of training participants used to train the predictive model (y-axis) in the ABCD and HCP datasets. Increasing training participants and scan time both led to increases in prediction performance. The * in both figures indicates that all available participants were used, therefore the sample size will be close to, but not exactly the number shown.

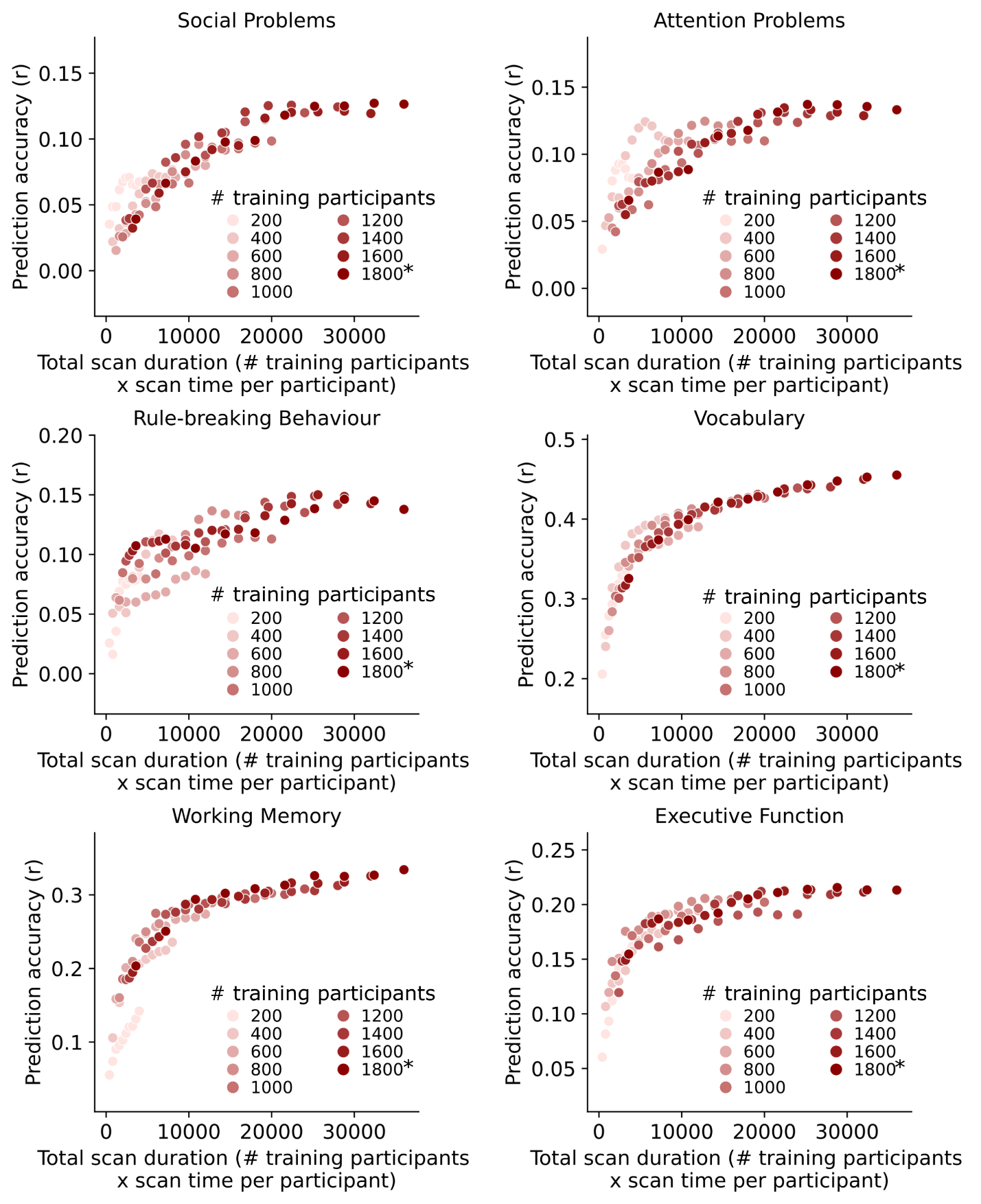

**Supplementary Fig. 7.1 |** Same as Fig. 2a except showing the scatter plots for 6 of the 17 phenotypic measures in the ABCD dataset that visually follow a logarithmic pattern. Scatter plots showing prediction accuracy (Pearson’s correlation) as a function of total scan duration (defined as # training participants x scan time per participant). The * in the figures indicates that all available participants were used, therefore the sample size will be close to, but not exactly the number shown.

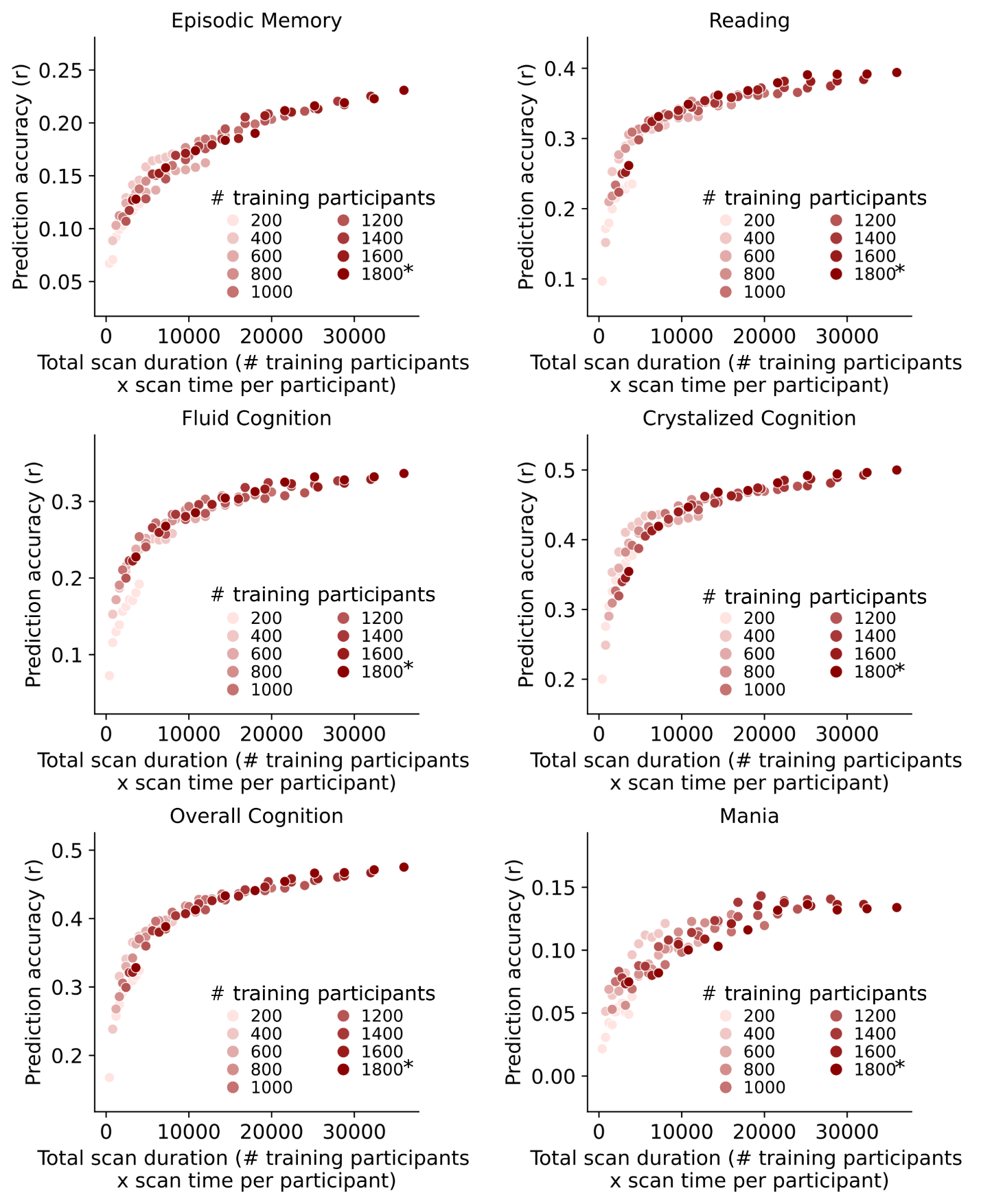

**Supplementary Fig. 7.2 |** Same as Fig. 2a except showing the scatter plots for 6 of the 17 phenotypic measures in the ABCD dataset that visually follow a logarithmic pattern. Scatter plots showing prediction accuracy (Pearson’s correlation) as a function of total scan duration (defined as # training participants x scan time per participant). The * in the figures indicates that all available participants were used, therefore the sample size will be close to, but not exactly the number shown.

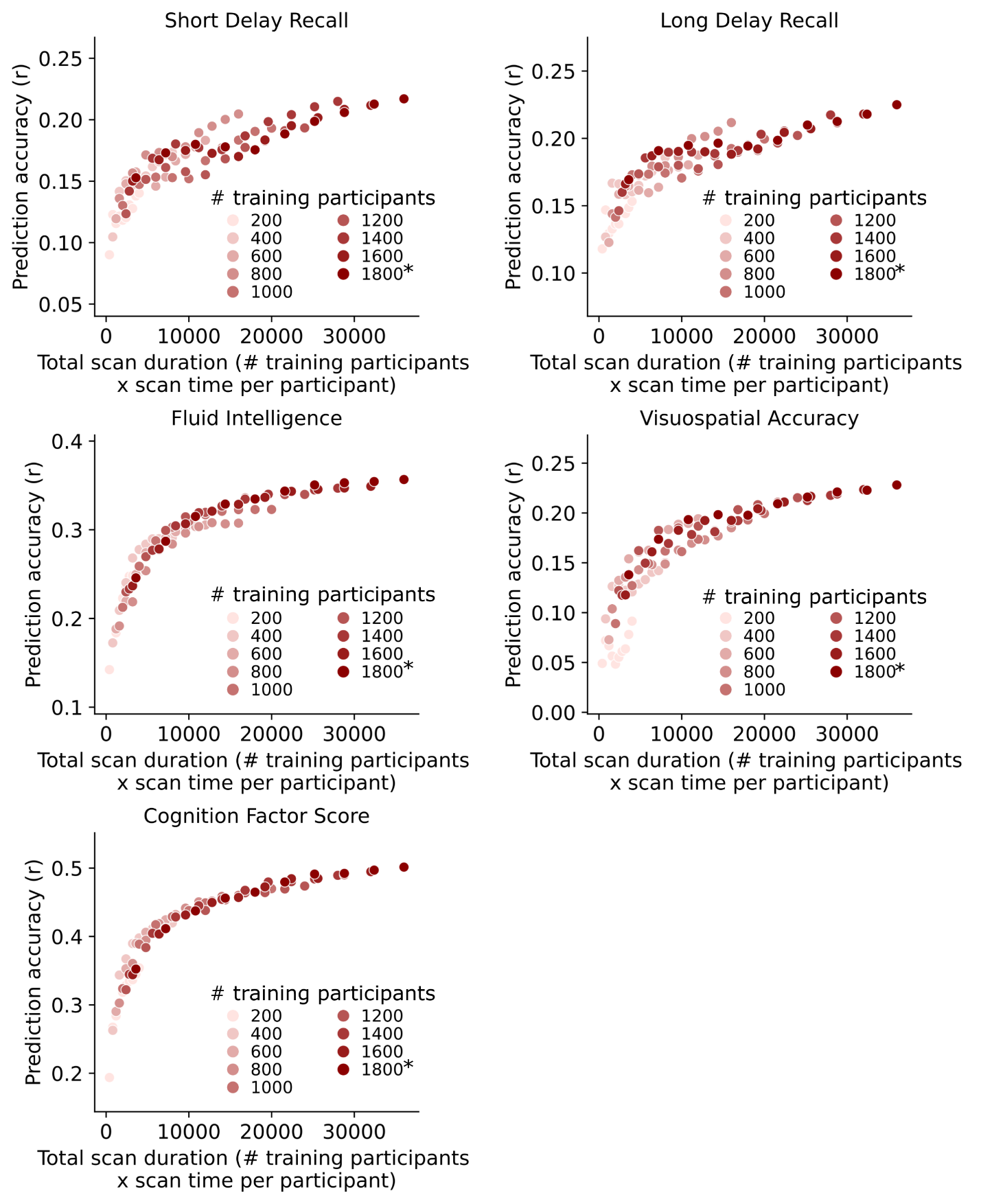

**Supplementary Fig. 7.3 |** Same as Fig. 2a except showing the scatter plots for 5 of the 17 phenotypic measures in the ABCD dataset that visually follow a logarithmic pattern. Scatter plots showing prediction accuracy (Pearson’s correlation) as a function of total scan duration (defined as # training participants x scan time per participant). The * in the figures indicates that all available participants were used, therefore the sample size will be close to, but not exactly the number shown.

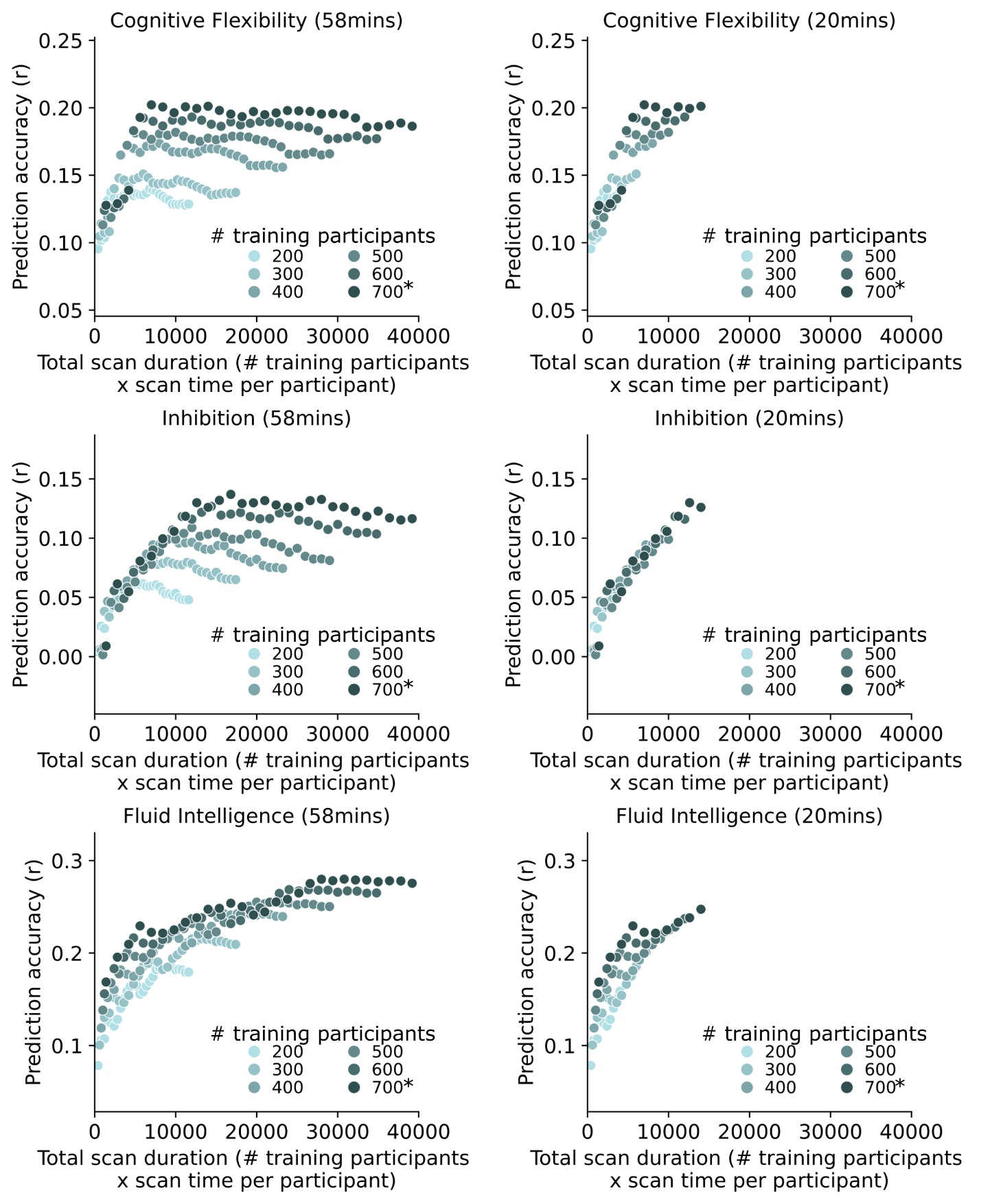

**Supplementary Fig. 8.1 |** Same as Fig. 2a except showing the scatter plots for 3 of the 19 phenotypic measures in the HCP dataset that visually follow a logarithmic pattern. Scatter plots showing prediction accuracy (Pearson’s correlation) as a function of total scan duration (defined as # training participants x scan time per participant). Scatter plots are shown for the full scan time (left panels) and up to 20 mins of scan time per participant (right panels). The * in the figures indicates that all available participants were used, therefore the sample size will be close to, but not exactly the number shown.

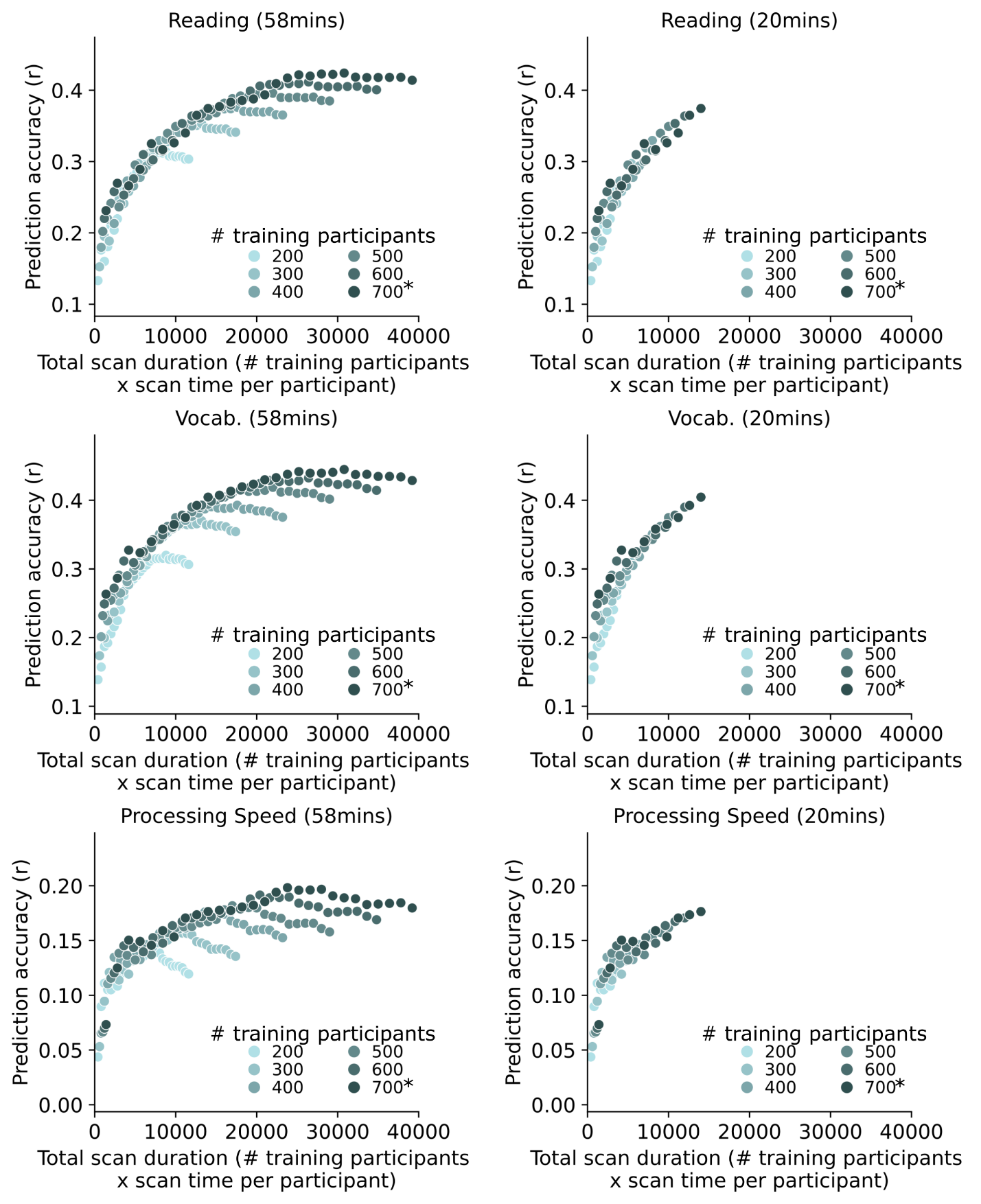

**Supplementary Fig. 8.2 |** Same as Fig. 2a except showing the scatter plots for 3 of the 19 phenotypic measures in the HCP dataset that visually follow a logarithmic pattern. Scatter plots showing prediction accuracy (Pearson’s correlation) as a function of total scan duration (defined as # training participants x scan time per participant). Scatter plots are shown for the full scan time (left panels) and up to 20 mins of scan time per participant (right panels). The * in the figures indicates that all available participants were used, therefore the sample size will be close to, but not exactly the number shown.

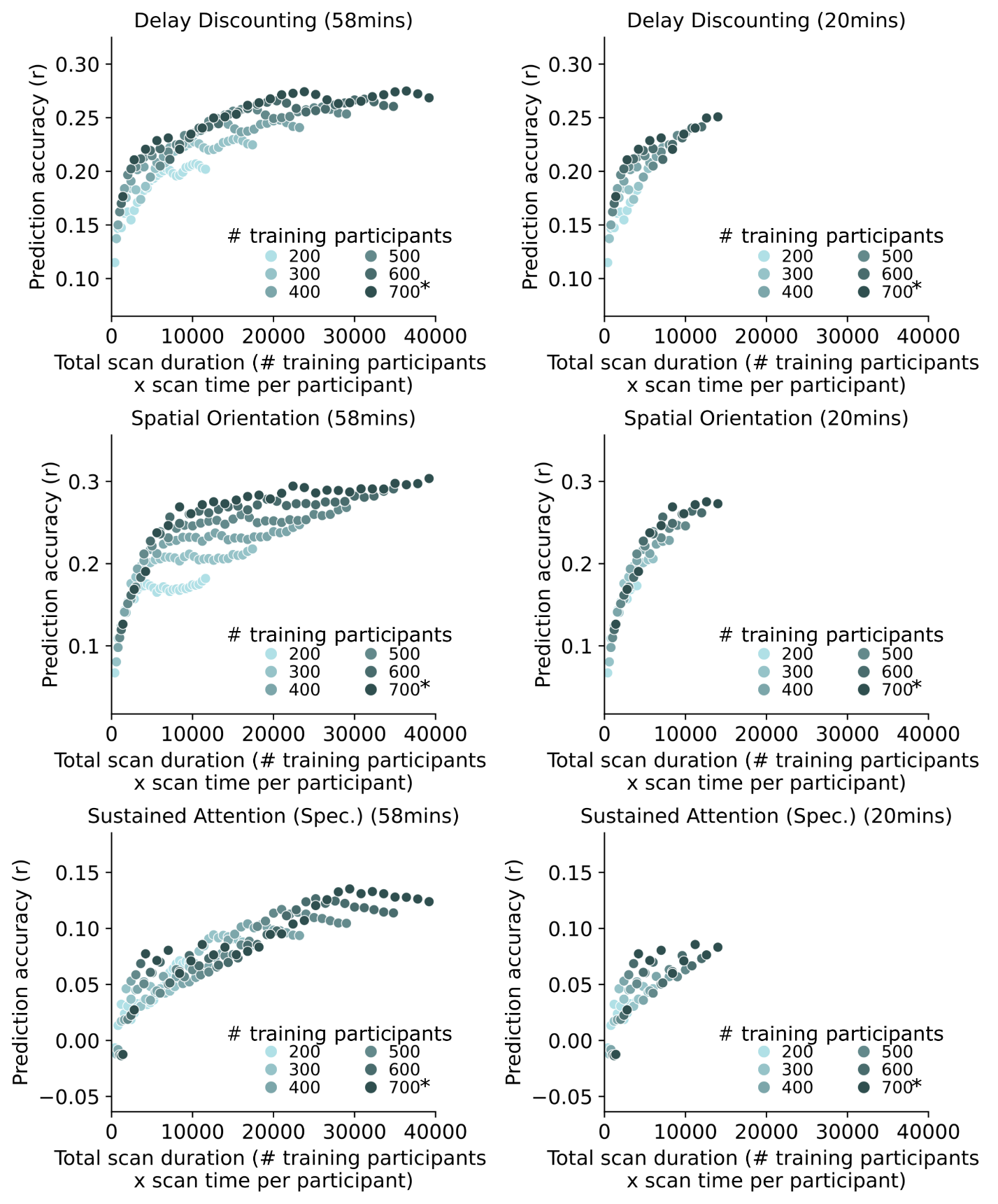

**Supplementary Fig. 8.3 |** Same as Fig. 2a except showing the scatter plots for 3 of the 19 phenotypic measures in the HCP dataset that visually follow a logarithmic pattern. Scatter plots showing prediction accuracy (Pearson’s correlation) as a function of total scan duration (defined as # training participants x scan time per participant). Scatter plots are shown for the full scan time (left panels) and up to 20 mins of scan time per participant (right panels). The * in the figures indicates that all available participants were used, therefore the sample size will be close to, but not exactly the number shown.

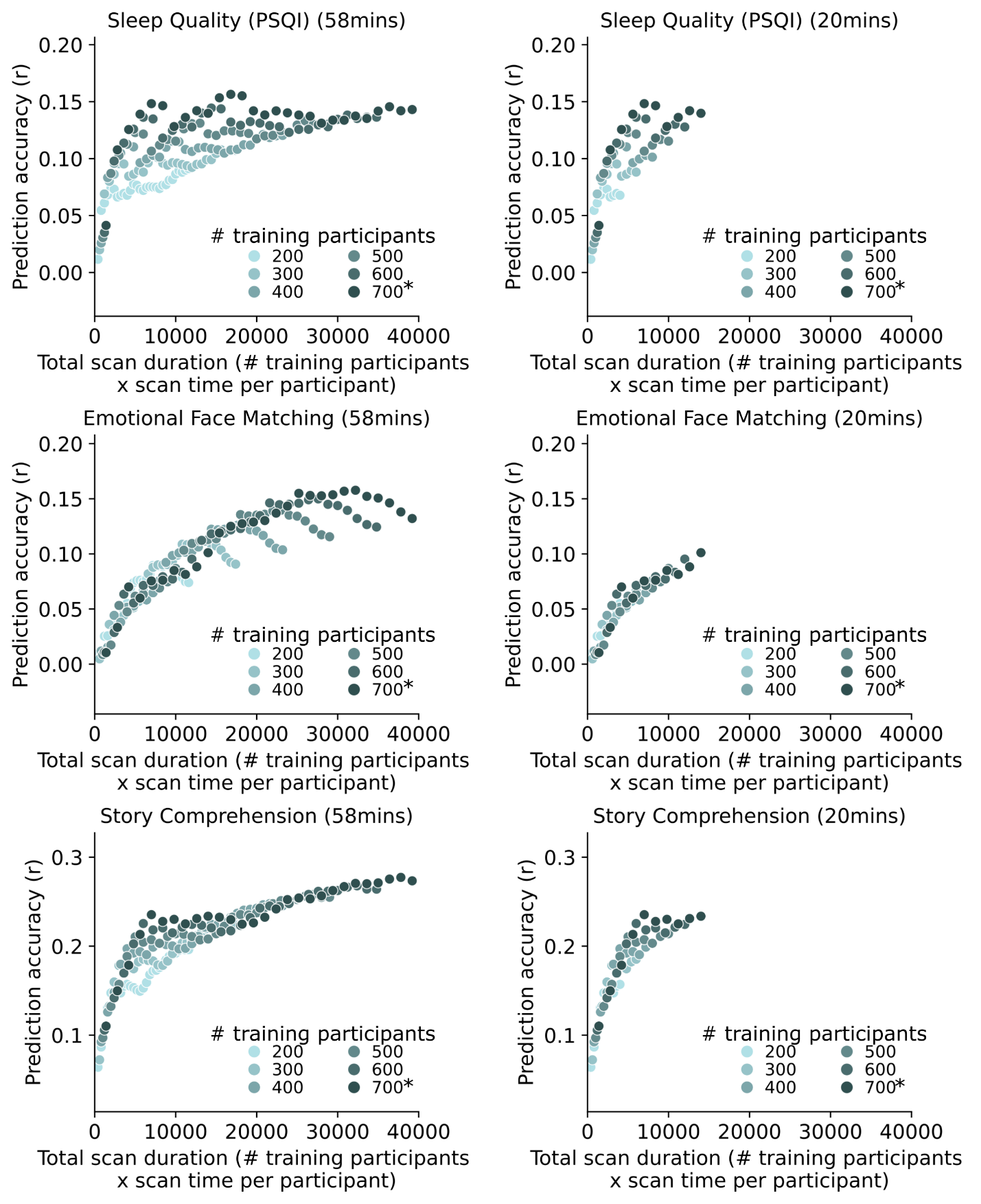

**Supplementary Fig. 8.4 |** Same as Fig. 2a except showing the scatter plots for 3 of the 19 phenotypic measures in the HCP dataset that visually follow a logarithmic pattern. Scatter plots showing prediction accuracy (Pearson’s correlation) as a function of total scan duration (defined as # training participants x scan time per participant). Scatter plots are shown for the full scan time (left panels) and up to 20 mins of scan time per participant (right panels). The * in the figures indicates that all available participants were used, therefore the sample size will be close to, but not exactly the number shown.

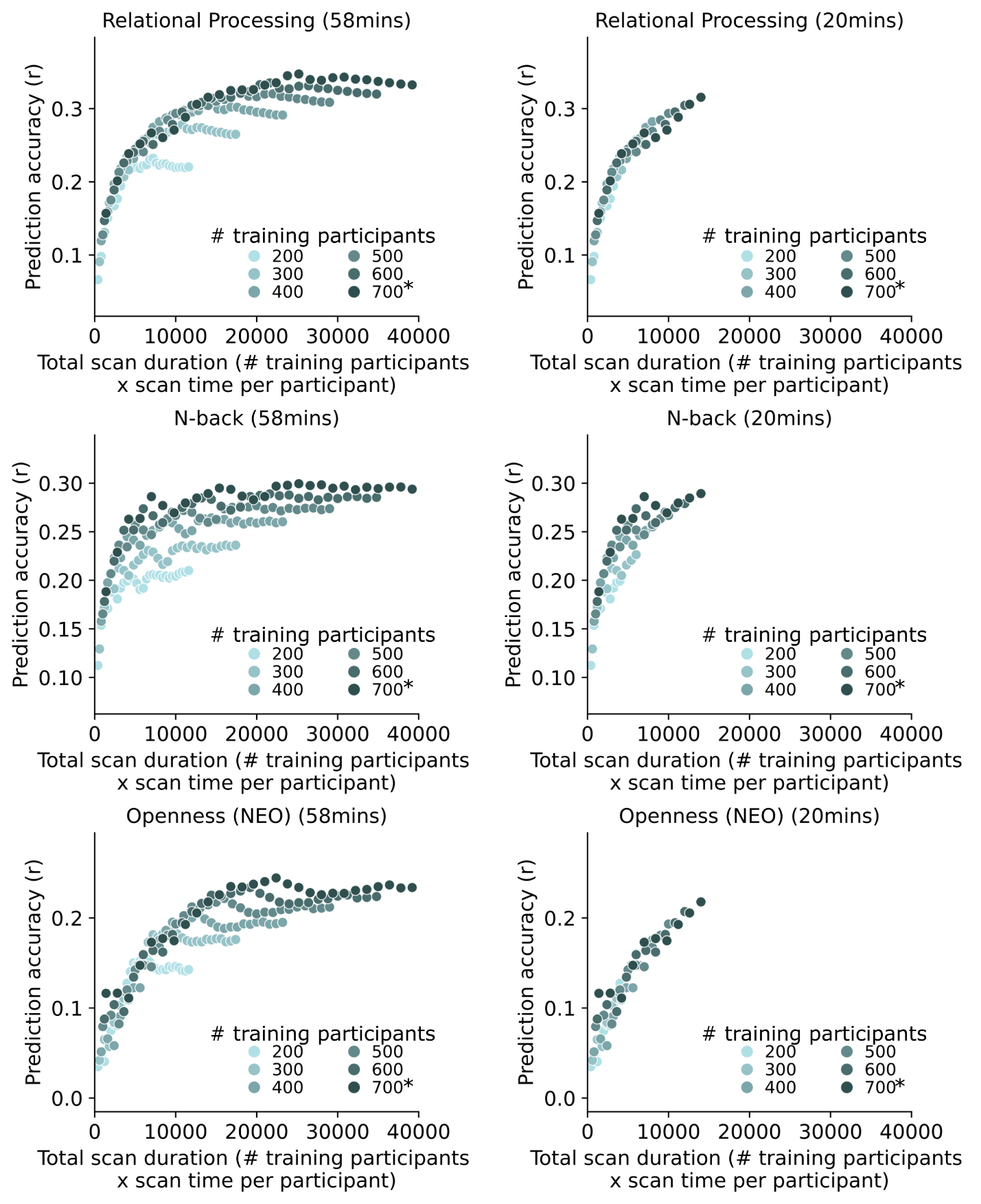

**Supplementary Fig. 8.5 |** Same as Fig. 2a except showing the scatter plots for 3 of the 19 phenotypic measures in the HCP dataset that visually follow a logarithmic pattern. Scatter plots showing prediction accuracy (Pearson’s correlation) as a function of total scan duration (defined as # training participants x scan time per participant). Scatter plots are shown for the full scan time (left panels) and up to 20 mins of scan time per participant (right panels). The * in the figures indicates that all available participants were used, therefore the sample size will be close to, but not exactly the number shown.

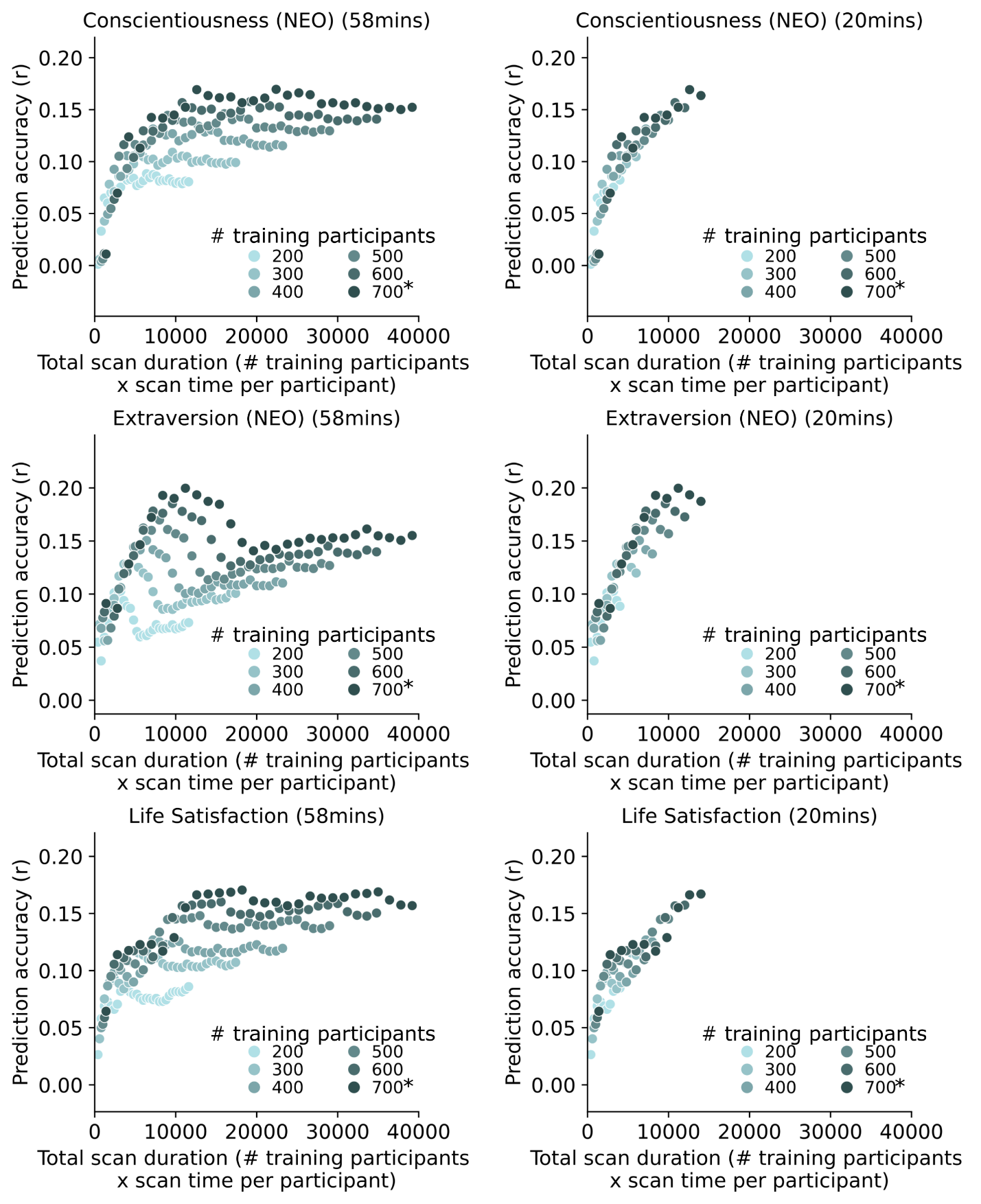

**Supplementary Fig. 8.6 |** Same as Fig. 2a except showing the scatter plots for 3 of the 19 phenotypic measures in the HCP dataset that visually follow a logarithmic pattern. Scatter plots showing prediction accuracy (Pearson’s correlation) as a function of total scan duration (defined as # training participants x scan time per participant). Scatter plots are shown for the full scan time (left panels) and up to 20 mins of scan time per participant (right panels). The * in the figures indicates that all available participants were used, therefore the sample size will be close to, but not exactly the number shown.

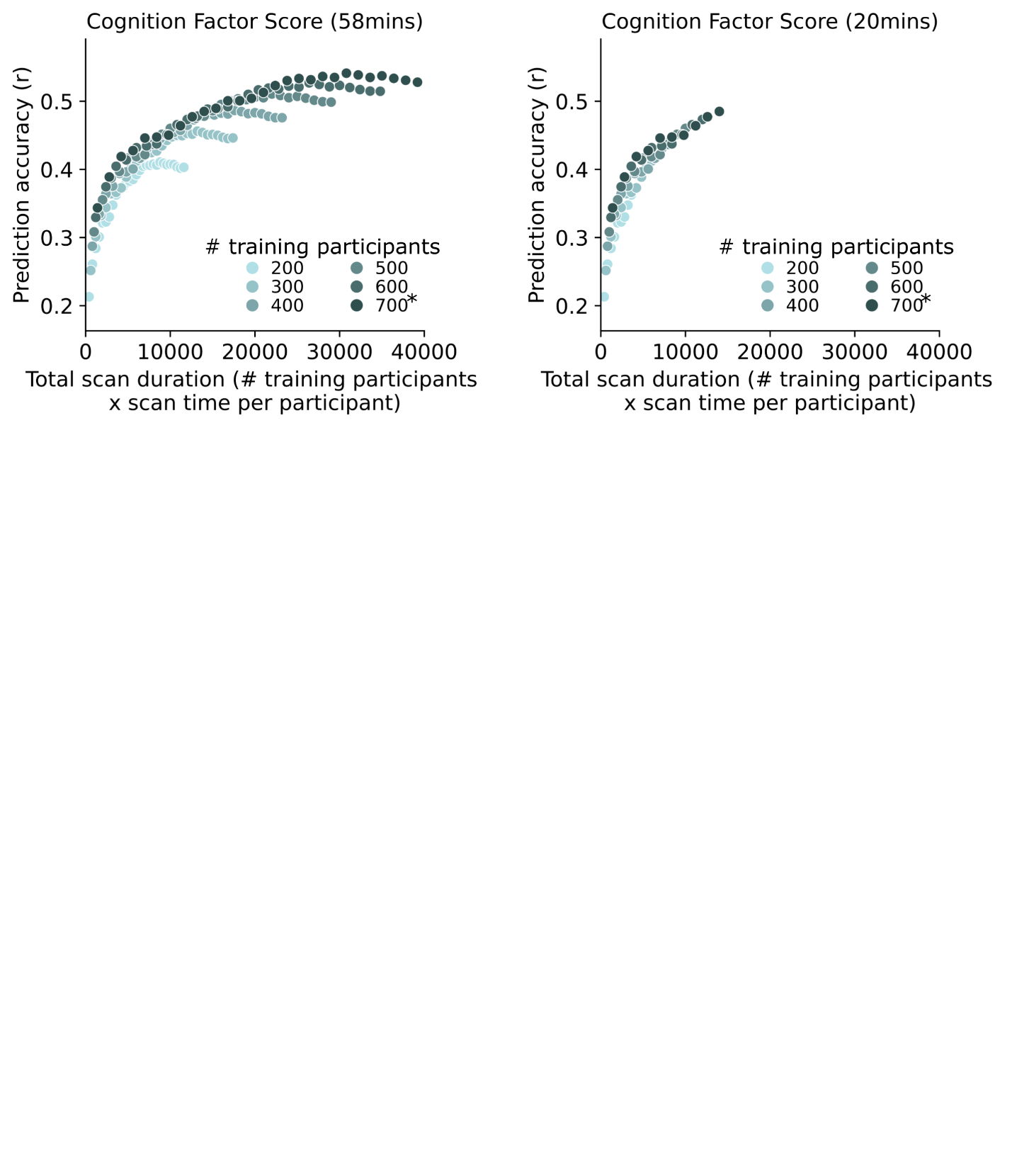

**Supplementary Fig. 8.7 |** Same as Fig. 2a except showing the scatter plots for 1 of the 19 phenotypic measures in the HCP dataset that visually follow a logarithmic pattern. Scatter plots showing prediction accuracy (Pearson’s correlation) as a function of total scan duration (defined as # training participants x scan time per participant). Scatter plots are shown for the full scan time (left panels) and up to 20 mins of scan time per participant (right panels). The * in the figures indicates that all available participants were used, therefore the sample size will be close to, but not exactly the number shown.

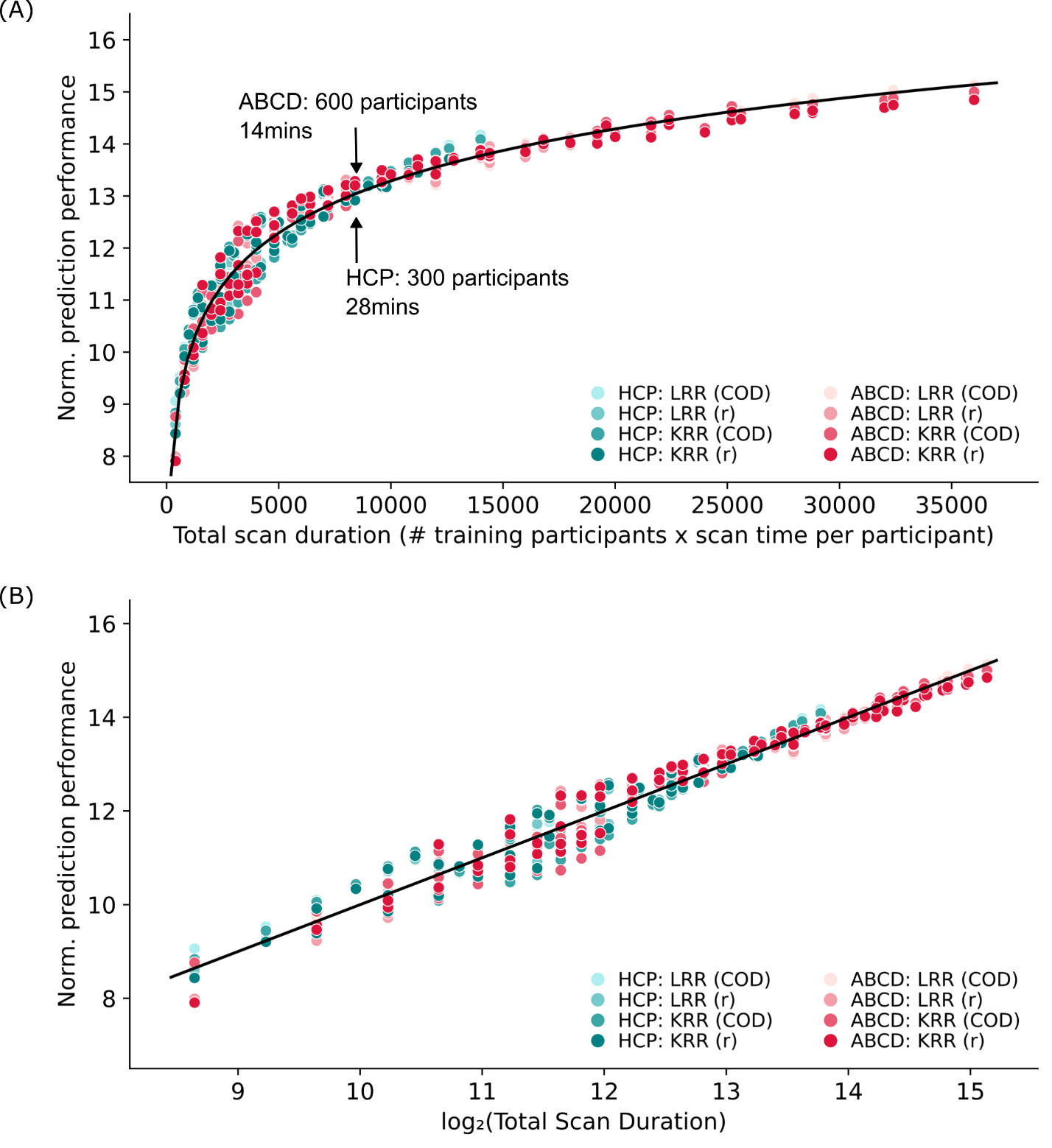

##### **Supplementary Fig. 9 | a.** Same as Fig. 2b1, except showing the relationship between total scan duration and prediction accuracy of for the cognitive factor scores versus total scan duration ignoring data beyond 20 min of scan time. Shown for different regressions (KRR, LRR) and different ways of calculating accuracy (Pearson’s correlation and Coefficient of Determination). Black arrows show that scanning 300 participants for 28 minutes (total scan duration = 300 × 28 = 8400 minutes) in the HCP dataset, or 600 participants for 14 minutes (total scan duration = 600 × 14 = 8400 minutes) in the ABCD study yielded very similar normalized prediction accuracies **b.** Same as Fig. 2b2, except showing the relationship between the logarithm of total scan duration and prediction accuracy of for the cognitive factor scores versus total scan duration ignoring data beyond 30 min of scan time. Shown for different regressions (KRR, LRR) and different ways of calculating accuracy (Pearson’s correlation and Coefficient of Determination).

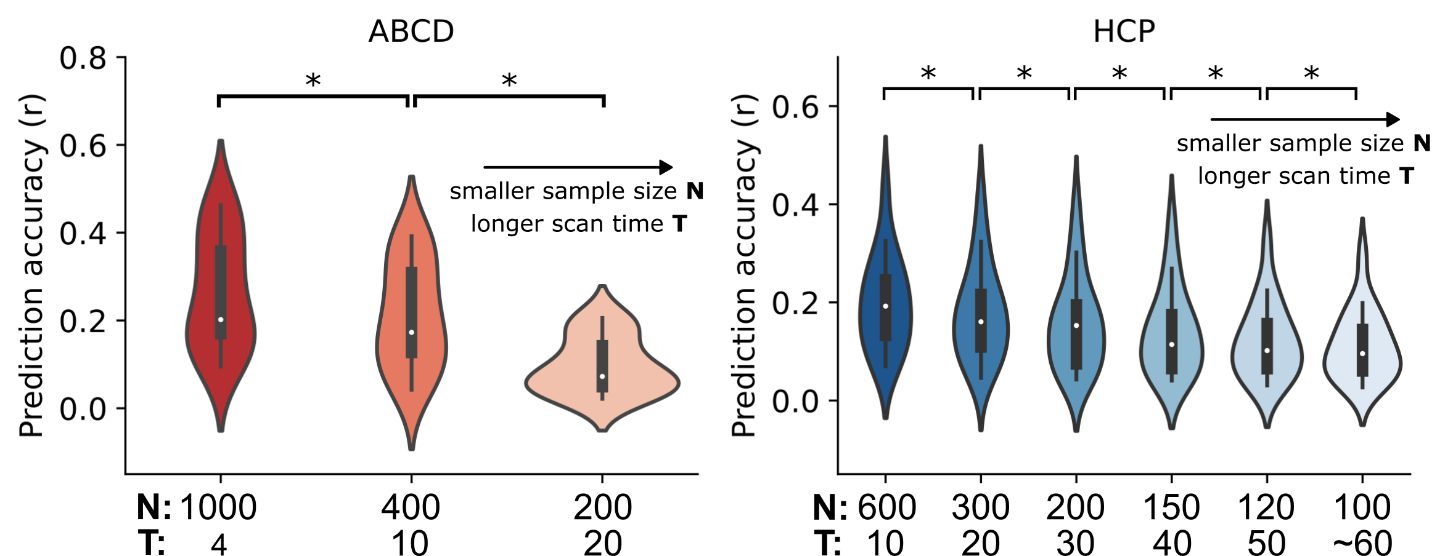

**Supplementary Fig. 10 |** Same as Fig. 3a, except in the left panel, each violin shows the distribution of average prediction accuracies across 17 ABCD study phenotypic measures. Each violin contains 17 data points and has a total scan duration of 4000 mins (left panel). In the right panel, each violin shows the distribution of average prediction accuracies across the 19 HCP phenotypic measures. Each violin contains 19 data points and has a total scan duration of 6000 mins (right panel). * indicates that the distributions of prediction accuracies were significantly different after false discovery rate (FDR) q < 0.05 correction.

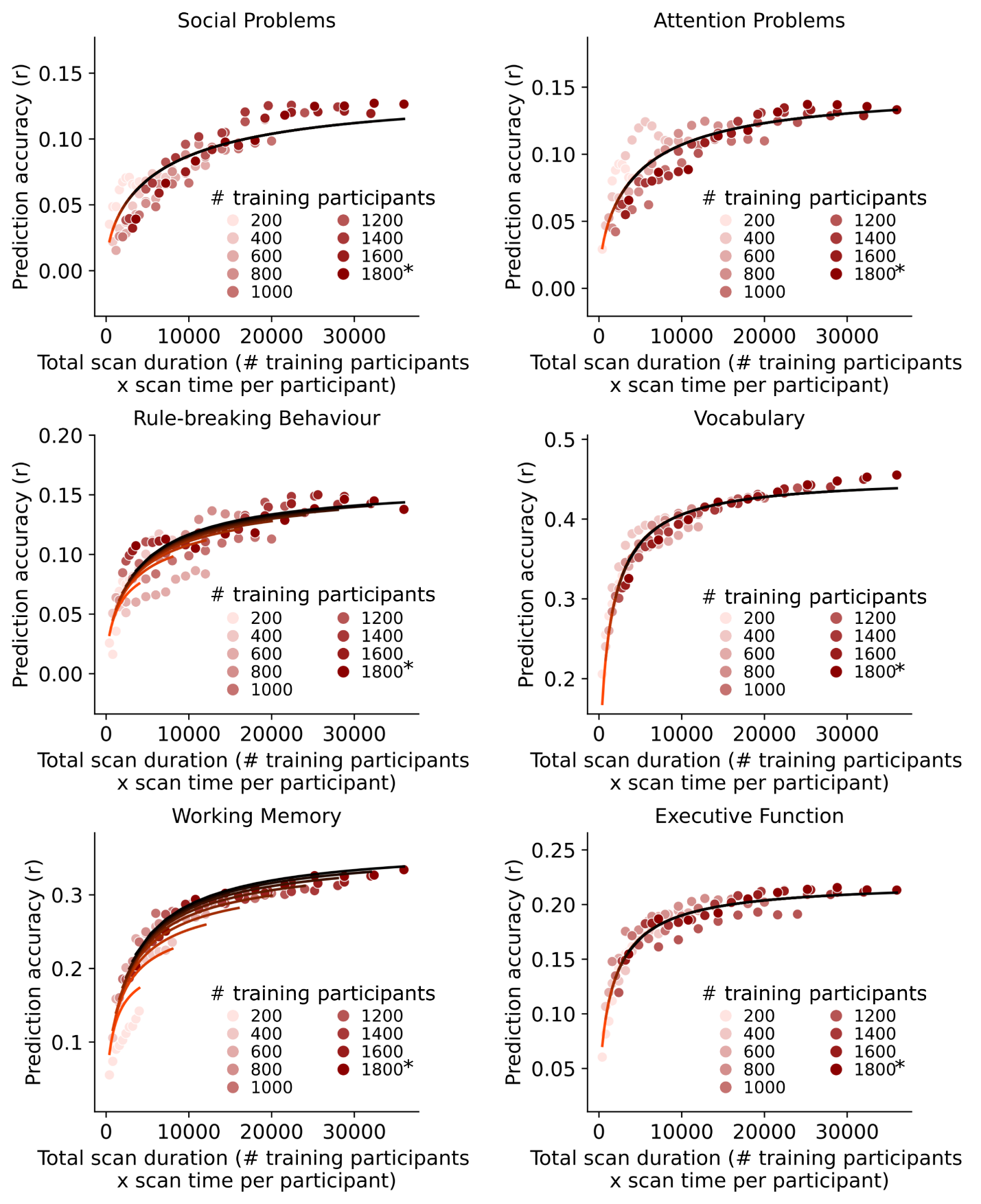

##### **Supplementary Fig. 11.1 |** Same as Fig. 3b except showing the scatter plots and the fit of prediction accuracy theoretical models for 6 of 17 phenotypic measures in the ABCD dataset that visually follow a logarithmic pattern. Scatter plot of prediction accuracy against total scan duration in the ABCD dataset. The curves were obtained by fitting a theoretical model to the prediction accuracies of the phenotype. The * in the figures indicates that all available participants were used, therefore the sample size will be close to, but not exactly the number shown.

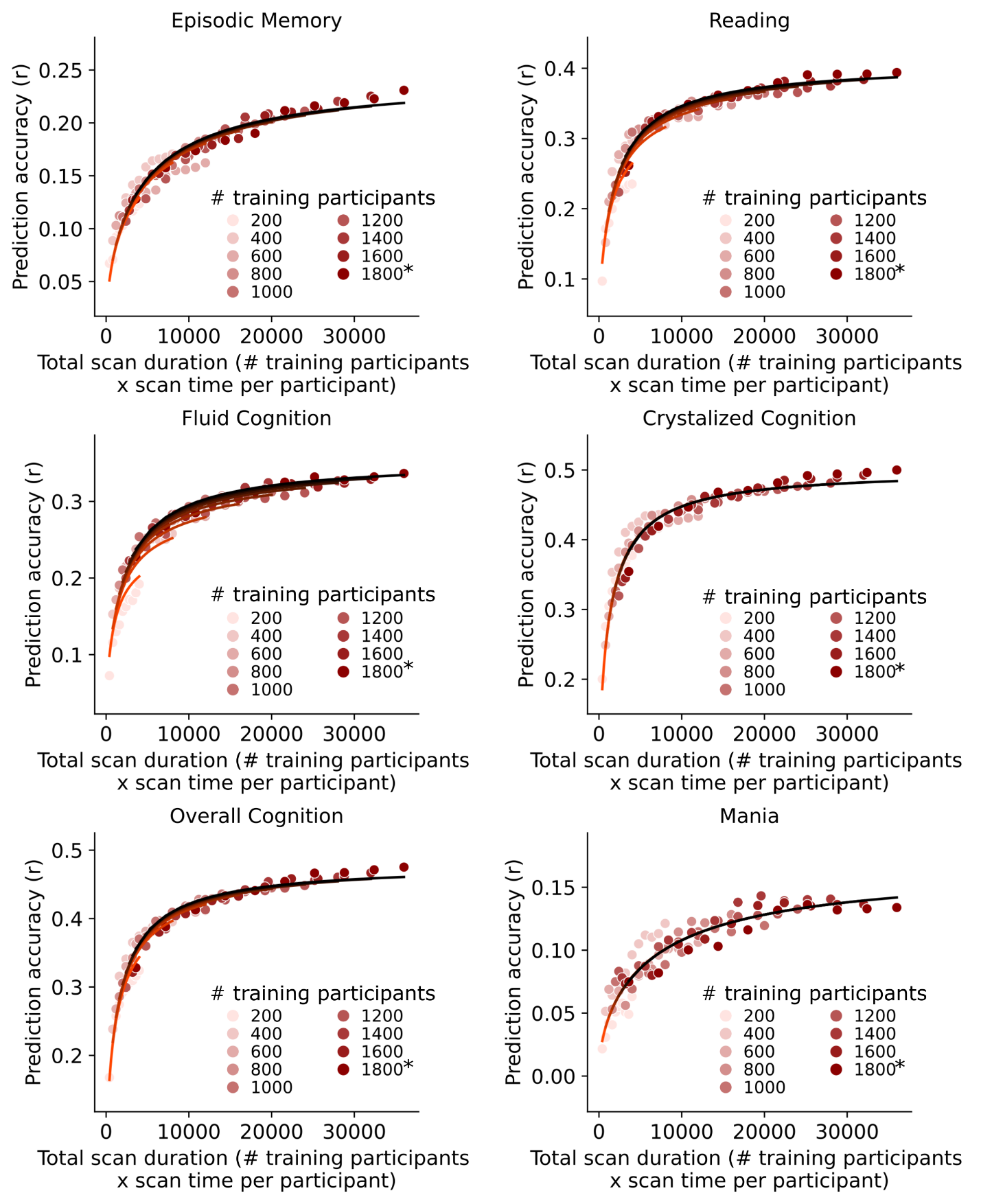

**Supplementary Fig. 11.2 |** Same as Fig. 3b except showing the scatter plots and the fit of prediction accuracy theoretical models for 6 of 17 phenotypic measures in the ABCD that visually follow a logarithmic pattern. Scatter plot of prediction accuracy against total scan duration in the ABCD dataset. The curves were obtained by fitting a theoretical model to the prediction accuracies of the phenotype. The * in the figures indicates that all available participants were used, therefore the sample size will be close to, but not exactly the number shown.

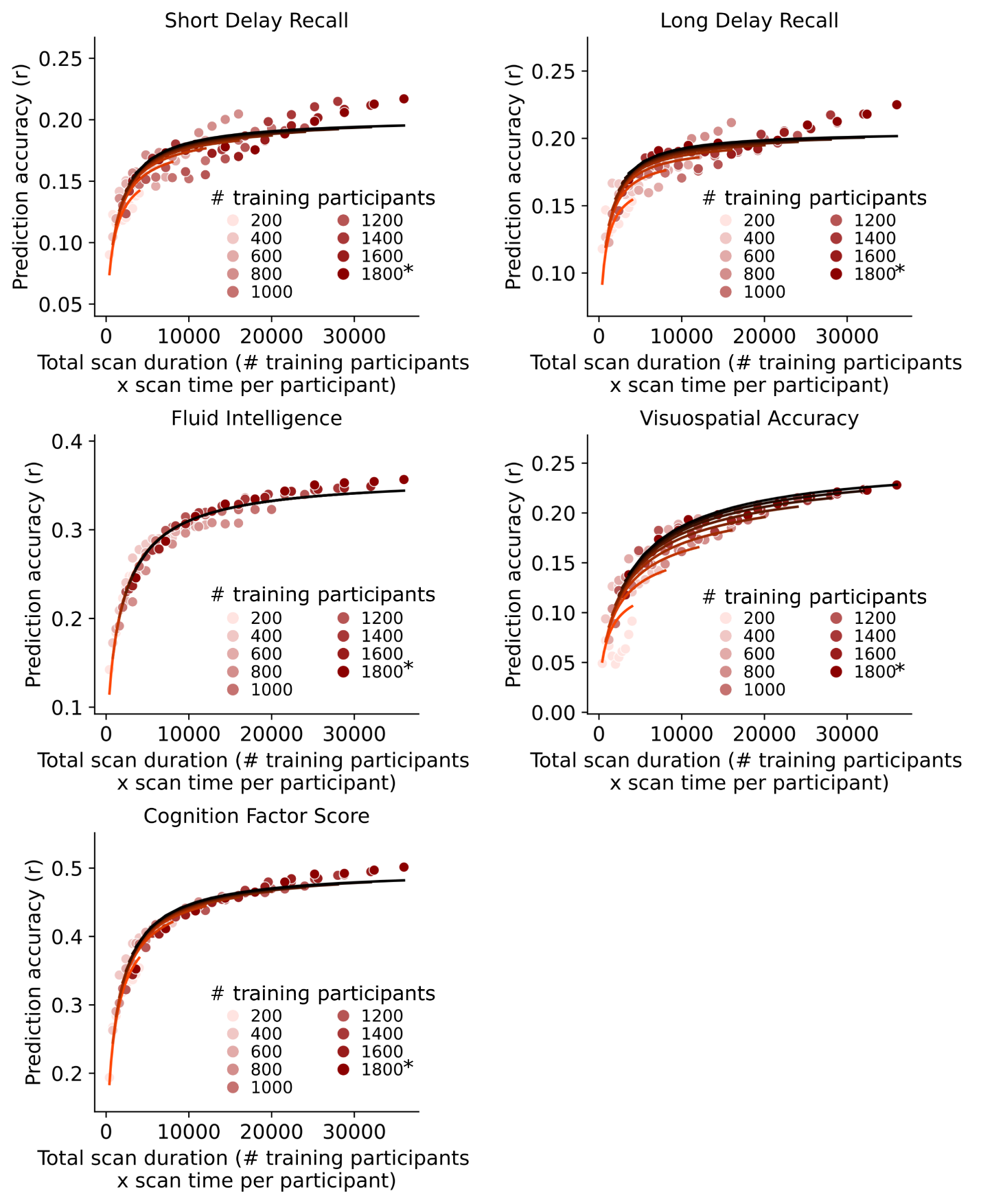

**Supplementary Fig. 11.3 |** Same as Fig. 3b except showing the scatter plots and the fit of prediction accuracy theoretical models for 5 of 17 phenotypic measures in the ABCD dataset that visually follow a logarithmic pattern. Scatter plot of prediction accuracy against total scan duration in the ABCD dataset. The curves were obtained by fitting a theoretical model to the prediction accuracies of the phenotype. The * in the figures indicates that all available participants were used, therefore the sample size will be close to, but not exactly the number shown.

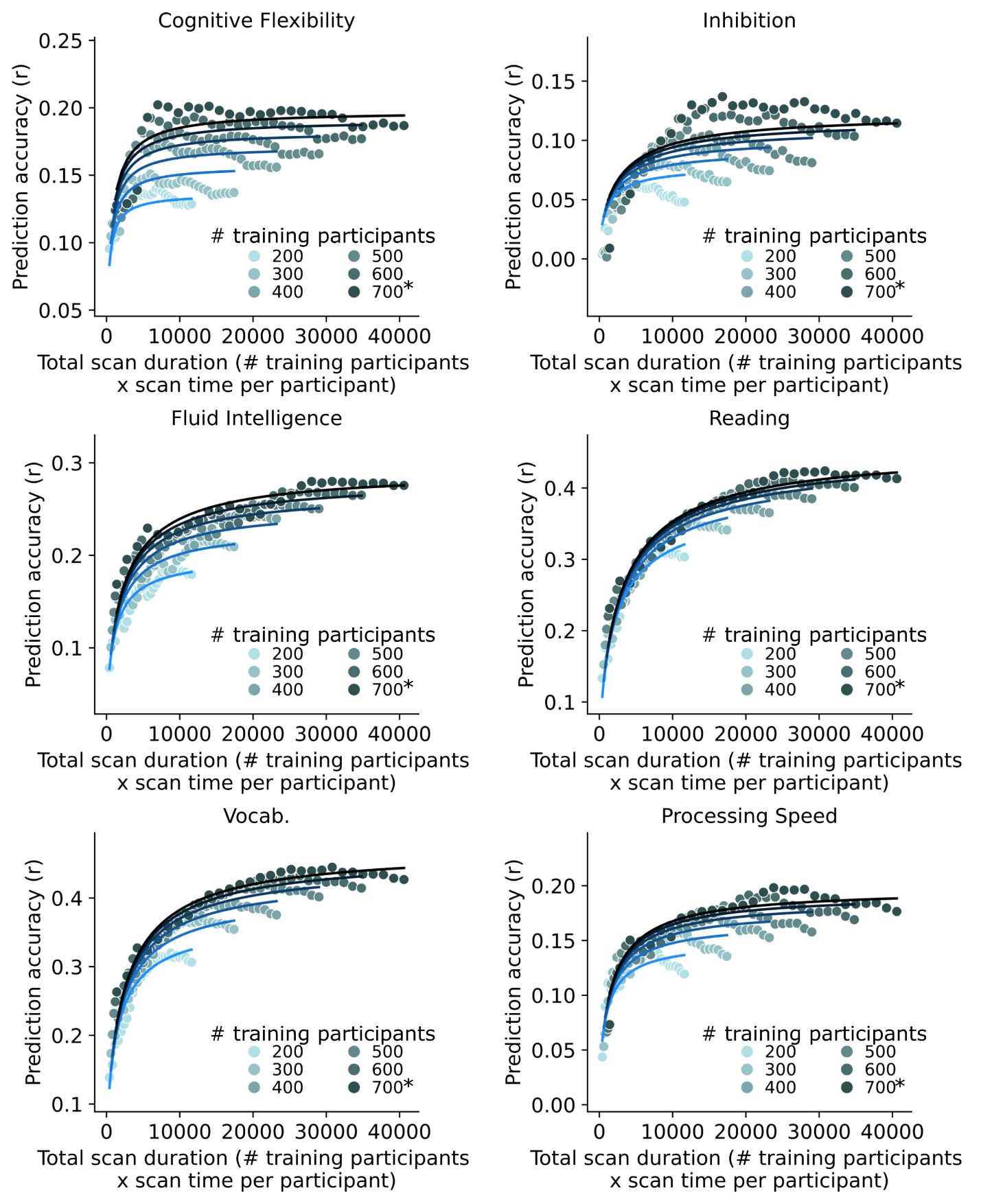

##### **Supplementary Fig. 12.1 |** Same as Fig. 3b except showing the scatter plots and the fit of prediction accuracy theoretical models for 6 of 19 phenotypic measures in the HCP that visually follow a logarithmic pattern. Scatter plot of prediction accuracy against total scan duration in the HCP dataset. The curves were obtained by fitting a theoretical model to the prediction accuracies of the phenotype. The * in the figures indicates that all available participants were used, therefore the sample size will be close to, but not exactly the number shown.

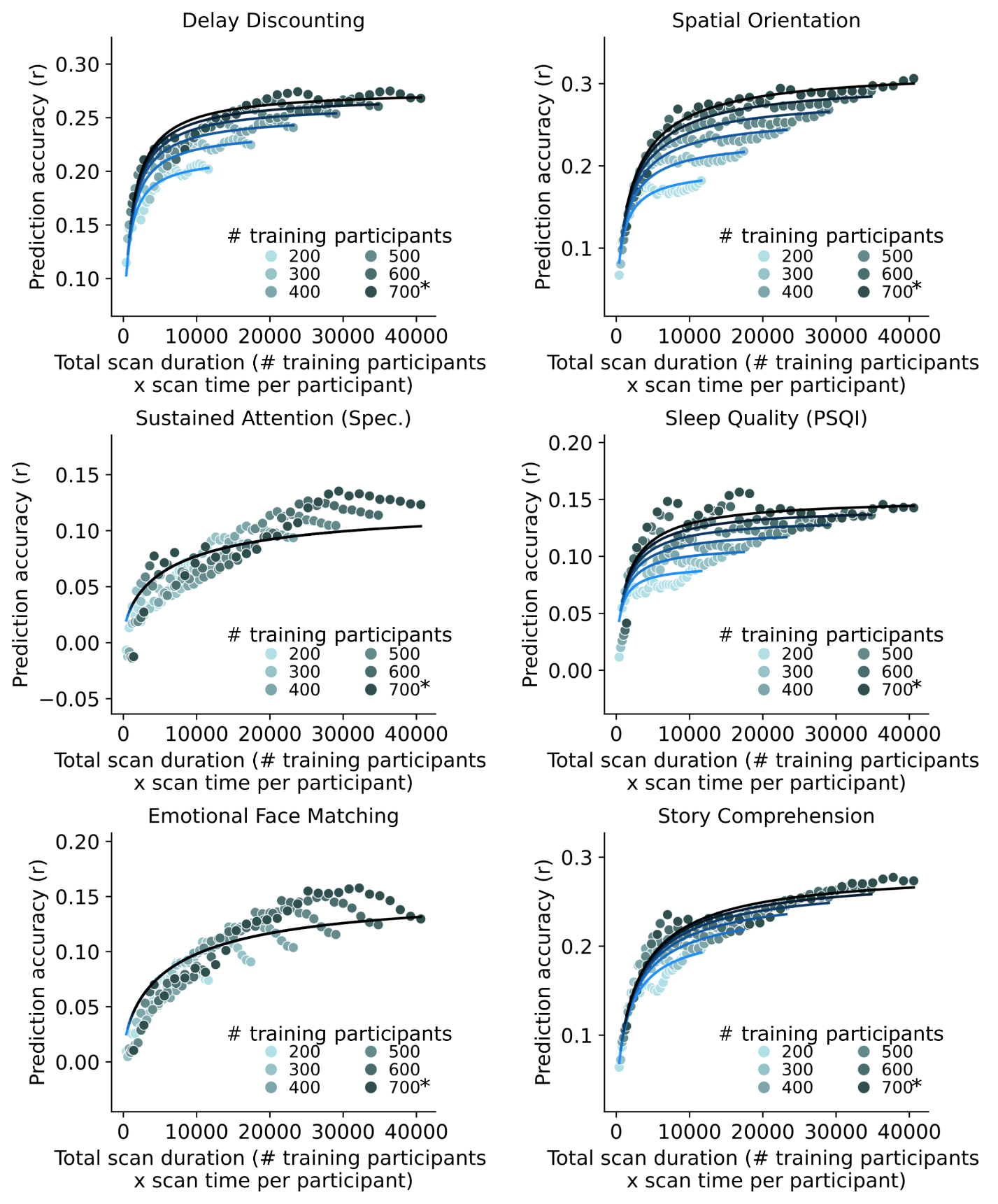

**Supplementary Fig. 12.2 |** Same as Fig. 3b except showing the scatter plots and the fit of prediction accuracy theoretical models for 6 of 19 phenotypic measures in the HCP that visually follow a logarithmic pattern. Scatter plot of prediction accuracy against total scan duration in the HCP dataset. The curves were obtained by fitting a theoretical model to the prediction accuracies of the phenotype. The * in the figures indicates that all available participants were used, therefore the sample size will be close to, but not exactly the number shown.

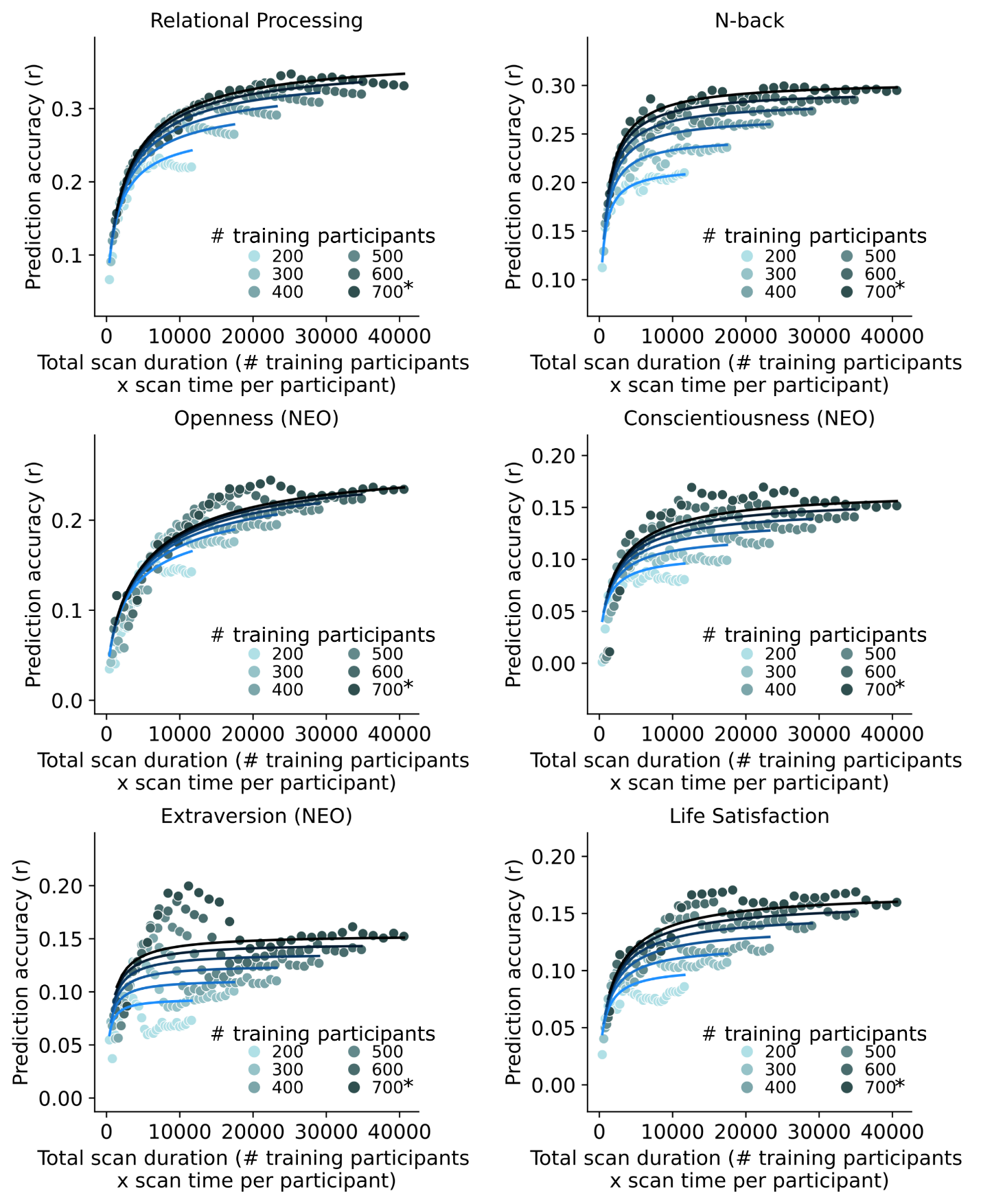

**Supplementary Fig. 12.3 |** Same as Fig. 3b except showing the scatter plots and the fit of prediction accuracy theoretical models for 6 of 19 phenotypic measures in the HCP that visually follow a logarithmic pattern. Scatter plot of prediction accuracy against total scan duration in the HCP dataset. The curves were obtained by fitting a theoretical model to the prediction accuracies of the phenotype. The * in the figures indicates that all available participants were used, therefore the sample size will be close to, but not exactly the number shown.

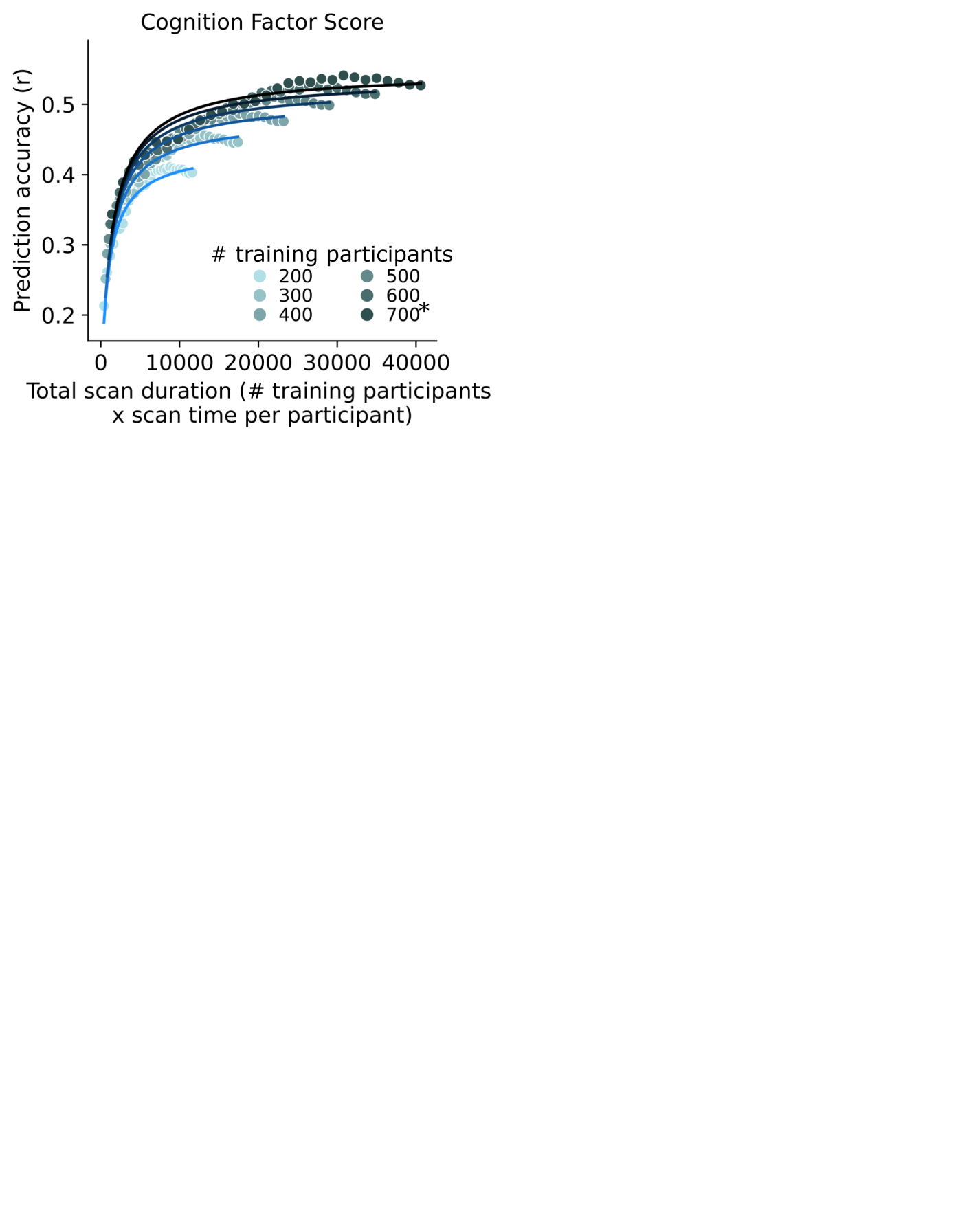

**Supplementary Fig. 12.4 |** Same as Fig. 3b except showing the scatter plots and the fit of prediction accuracy theoretical models for 1 of 19 phenotypic measures in the HCP that visually follow a logarithmic pattern. Scatter plot of prediction accuracy against total scan duration in the HCP dataset. The curves were obtained by fitting a theoretical model to the prediction accuracies of the phenotype. The * in the figures indicates that all available participants were used, therefore the sample size will be close to, but not exactly the number shown.

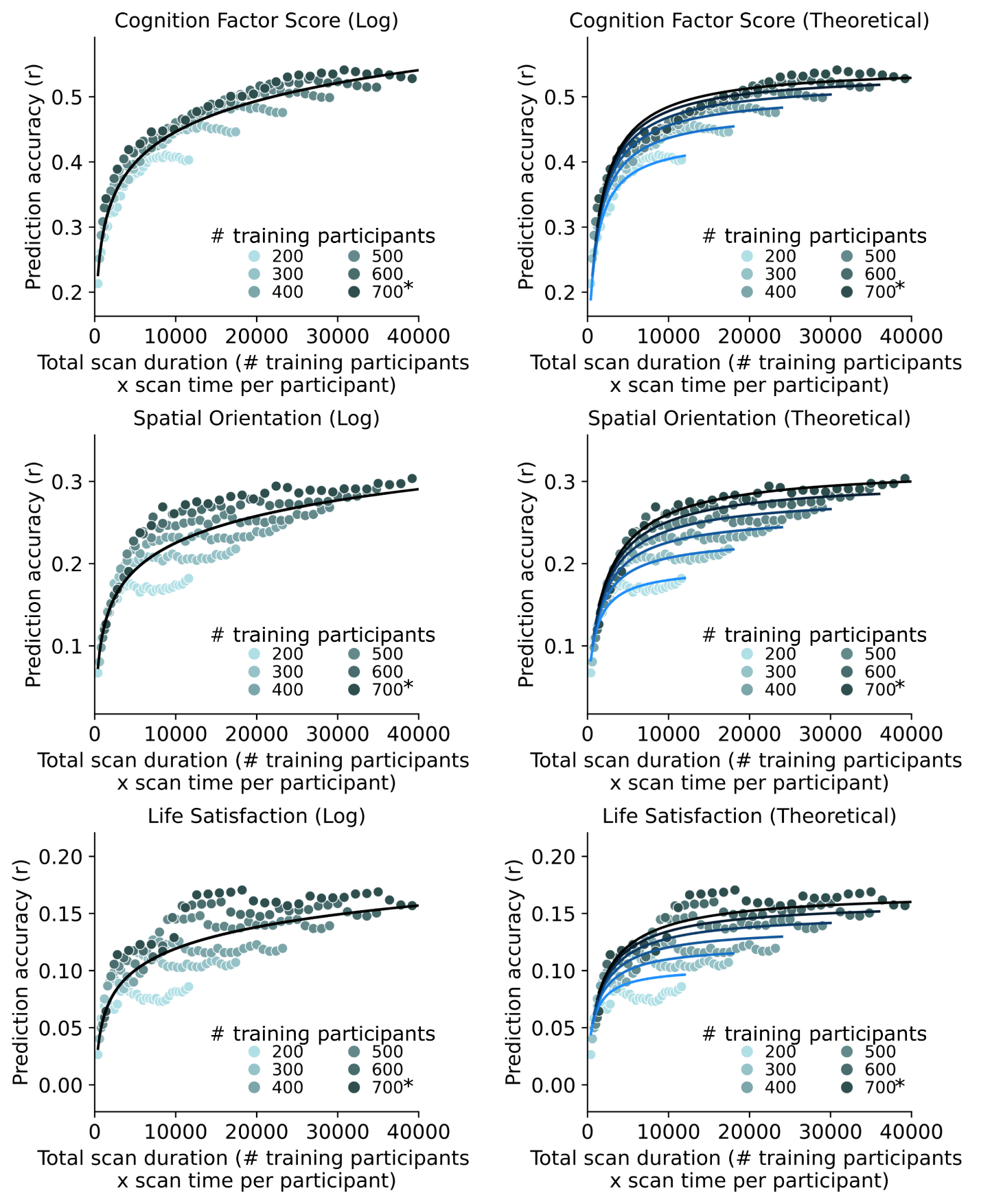

##### **Supplementary Fig. 13 |** Visual comparison of logarithmic and theoretical models fitted to three HCP phenotypes using the full 58 minutes of data. Because the logarithm model treated the sample size N and scan time T as being interchangeable, it was not able to explain the diminishing returns of scan time T relative to sample size N for larger value of T.

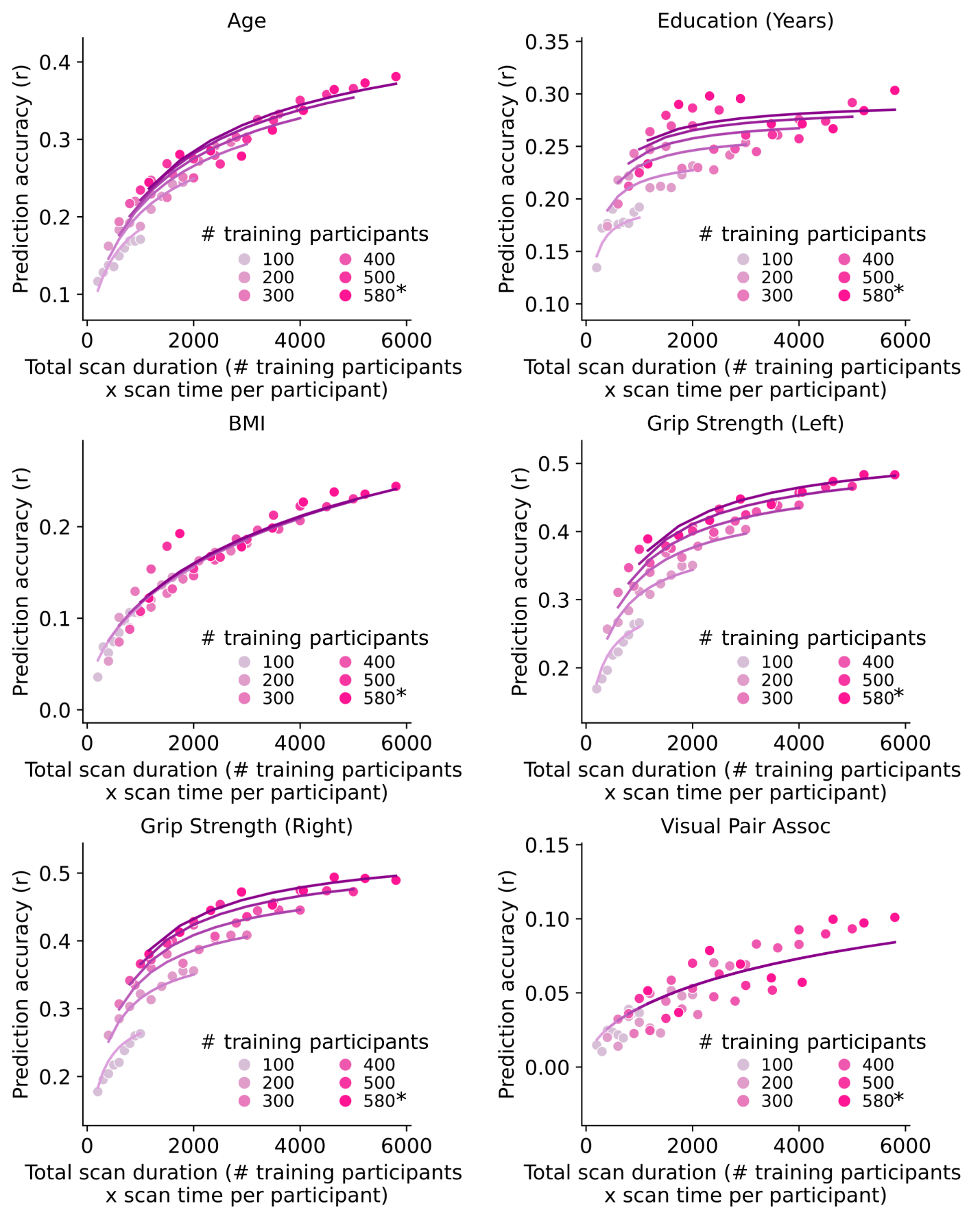

##### **Supplementary Fig. 14.1 |** Same as Fig. 3b except showing the scatter plots and the fit of prediction accuracy theoretical models for 6 of 14 phenotypic measures in the SINGER dataset that exhibit a good fit to the theoretical model. The curves were obtained by fitting a theoretical model to the prediction accuracies of the phenotype. The * in the figures indicates that all available participants were used, therefore the sample size will be close to, but not exactly the number shown.

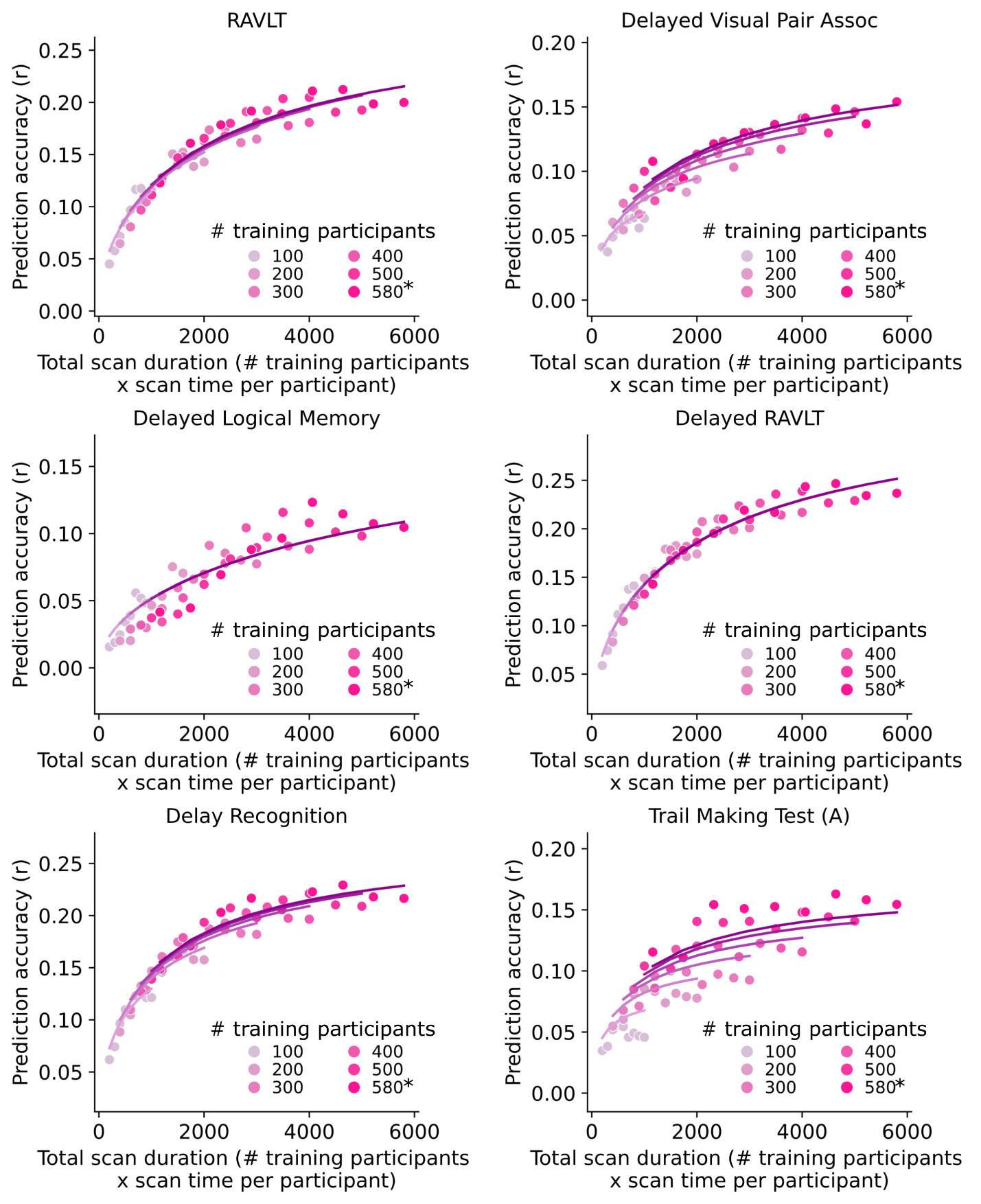

**Supplementary Fig. 14.2 |** Same as Fig. 3b except showing the scatter plots and the fit of prediction accuracy theoretical models for 6 of 14 phenotypic measures in the SINGER dataset that exhibit a good fit to the theoretical model. The curves were obtained by fitting a theoretical model to the prediction accuracies of the phenotype. The * in the figures indicates that all available participants were used, therefore the sample size will be close to, but not exactly the number shown.

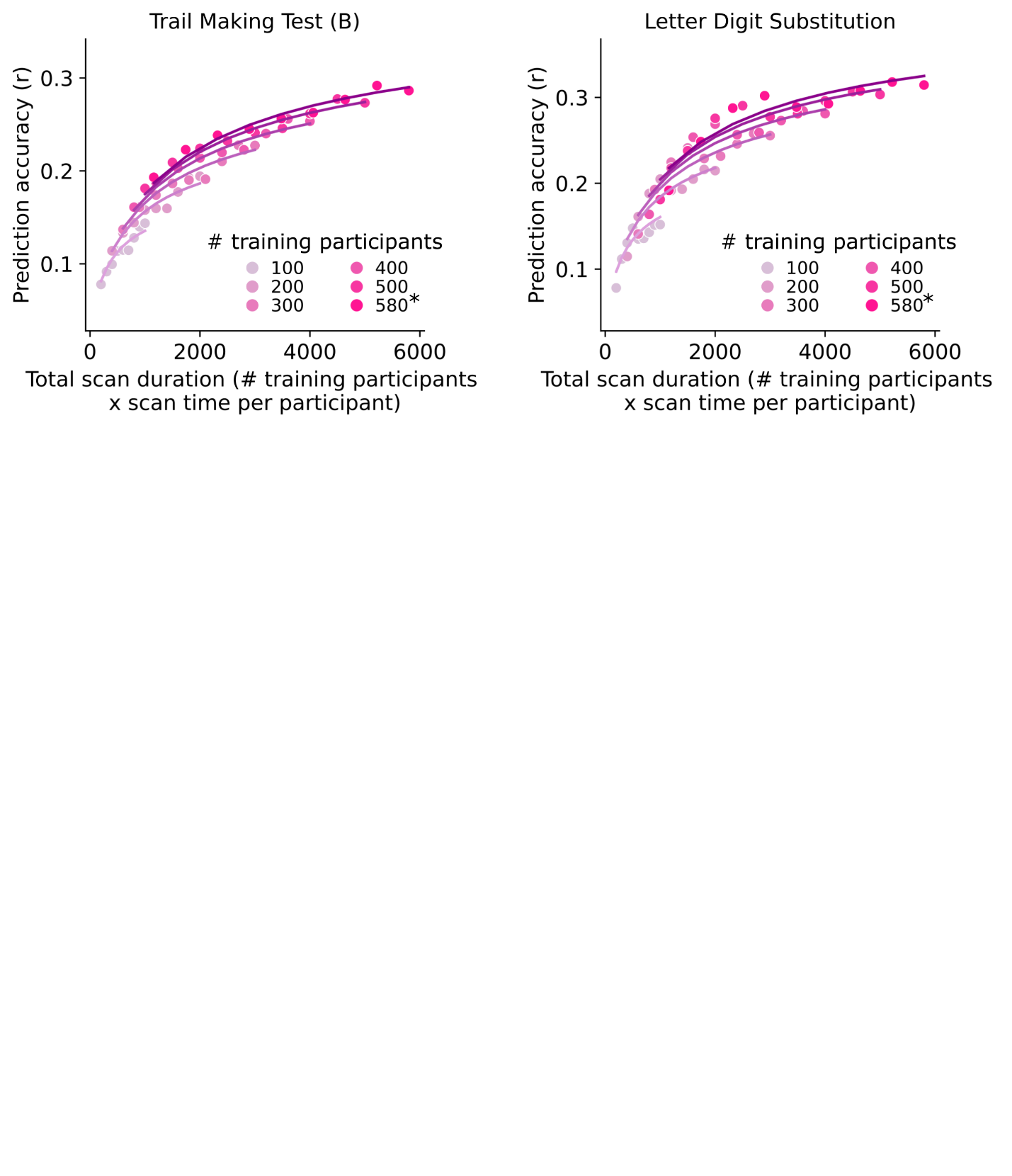

**Supplementary Fig. 14.3 |** Same as Fig. 3b except showing the scatter plots and the fit of prediction accuracy theoretical models for 2 of 14 phenotypic measures in the SINGER dataset that exhibit a good fit to the theoretical model. The curves were obtained by fitting a theoretical model to the prediction accuracies of the phenotype. The * in the figures indicates that all available participants were used, therefore the sample size will be close to, but not exactly the number shown.

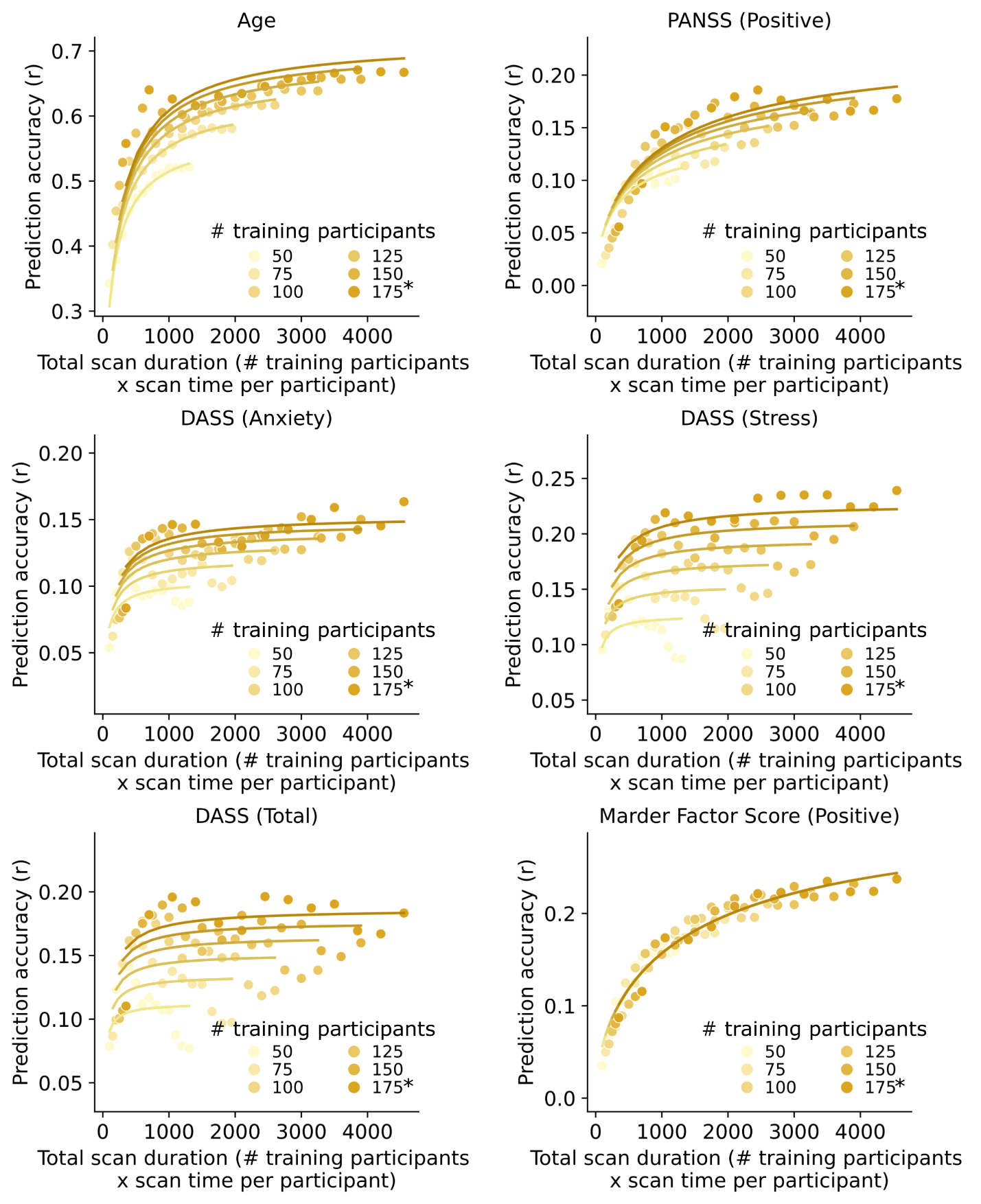

##### **Supplementary Fig. 15.1 |** Same as Fig. 3b except showing the scatter plots and the fit of prediction accuracy theoretical models for 6 of 7 phenotypic measures in the TCP dataset. The curves were obtained by fitting a theoretical model to the prediction accuracies of the phenotype. The * in the figures indicates that all available participants were used, therefore the sample size will be close to, but not exactly the number shown.

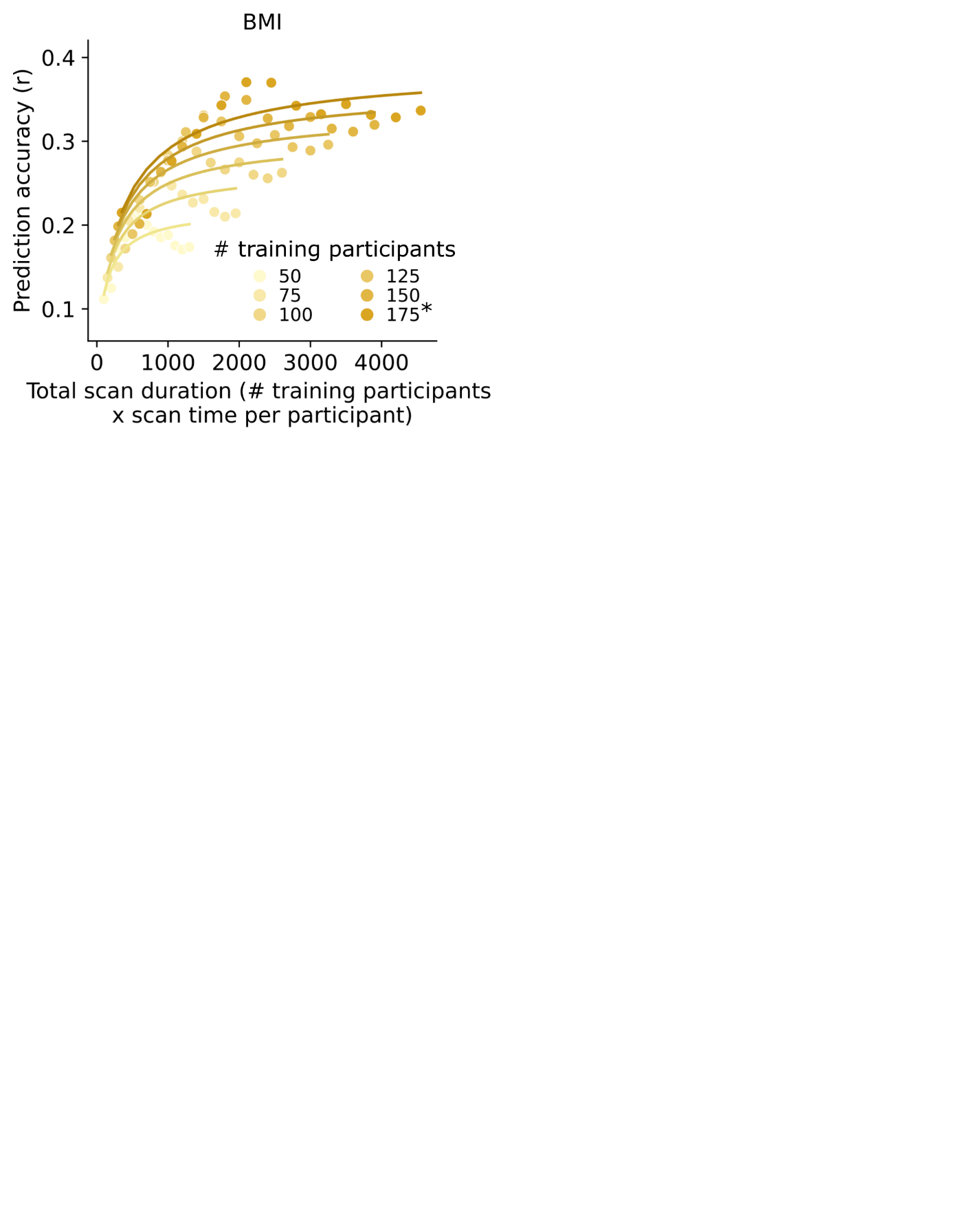
**Supplementary Fig. 15.2 |** Same as Fig. 3b except showing the scatter plots and the fit of prediction accuracy theoretical models for 1 of 7 phenotypic measures in the TCP dataset. The curves were obtained by fitting a theoretical model to the prediction accuracies of the phenotype. The * in the figures indicates that all available participants were used, therefore the sample size will be close to, but not exactly the number shown.

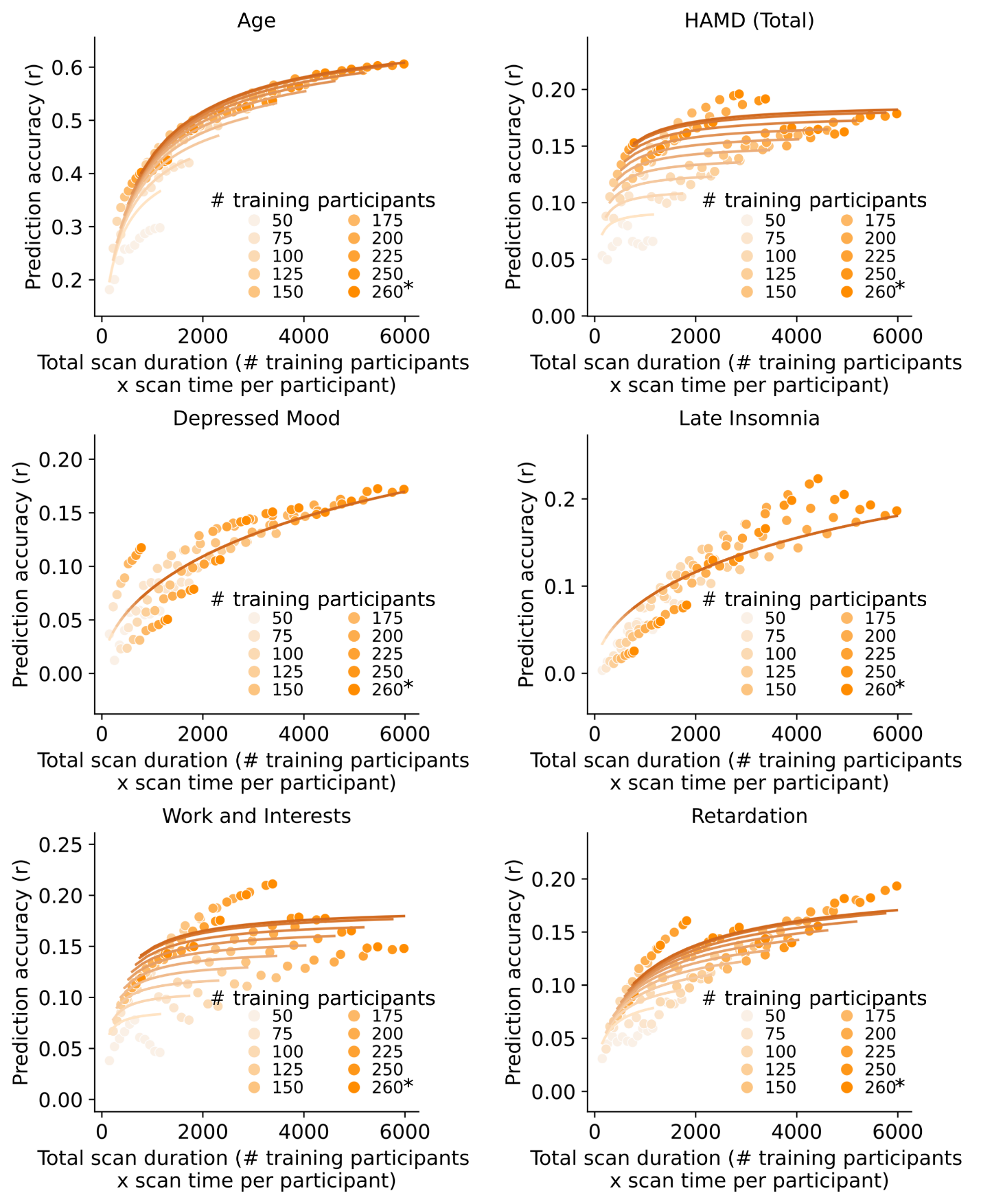

##### **Supplementary Fig. 16.1 |** Same as Fig. 3b except showing the scatter plots and the fit of prediction accuracy theoretical models for 6 of 7 phenotypic measures in the MDD dataset. The curves were obtained by fitting a theoretical model to the prediction accuracies of the phenotype. The * in the figures indicates that all available participants were used, therefore the sample size will be close to, but not exactly the number shown.

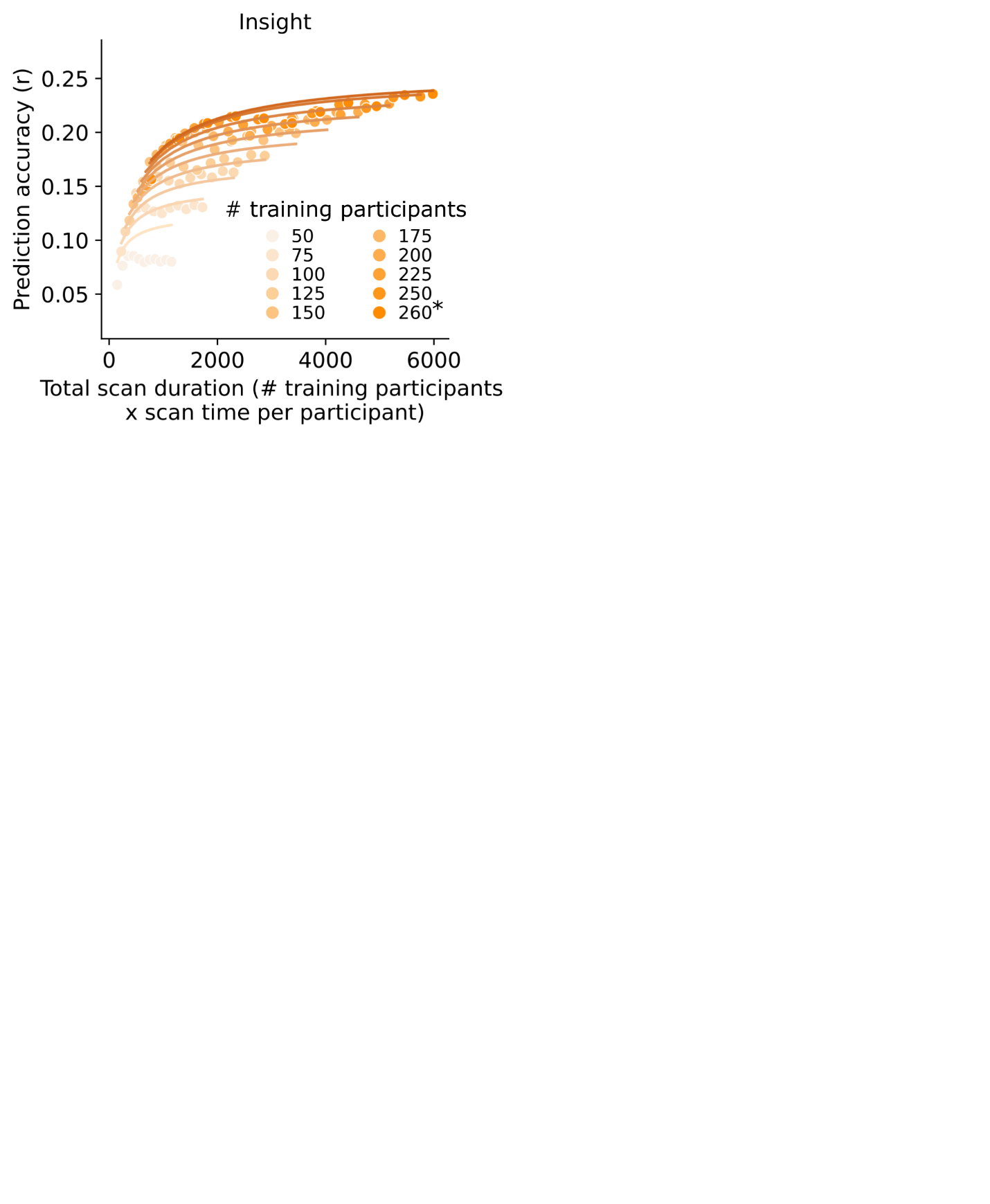
**Fig. 16.2 |** Same as Fig. 3b except showing the scatter plots and the fit of prediction accuracy theoretical models for 1 of 7 phenotypic measures in the MDD dataset. The curves were obtained by fitting a theoretical model to the prediction accuracies of the phenotype. The * in the figures indicates that all available participants were used, therefore the sample size will be close to, but not exactly the number shown.

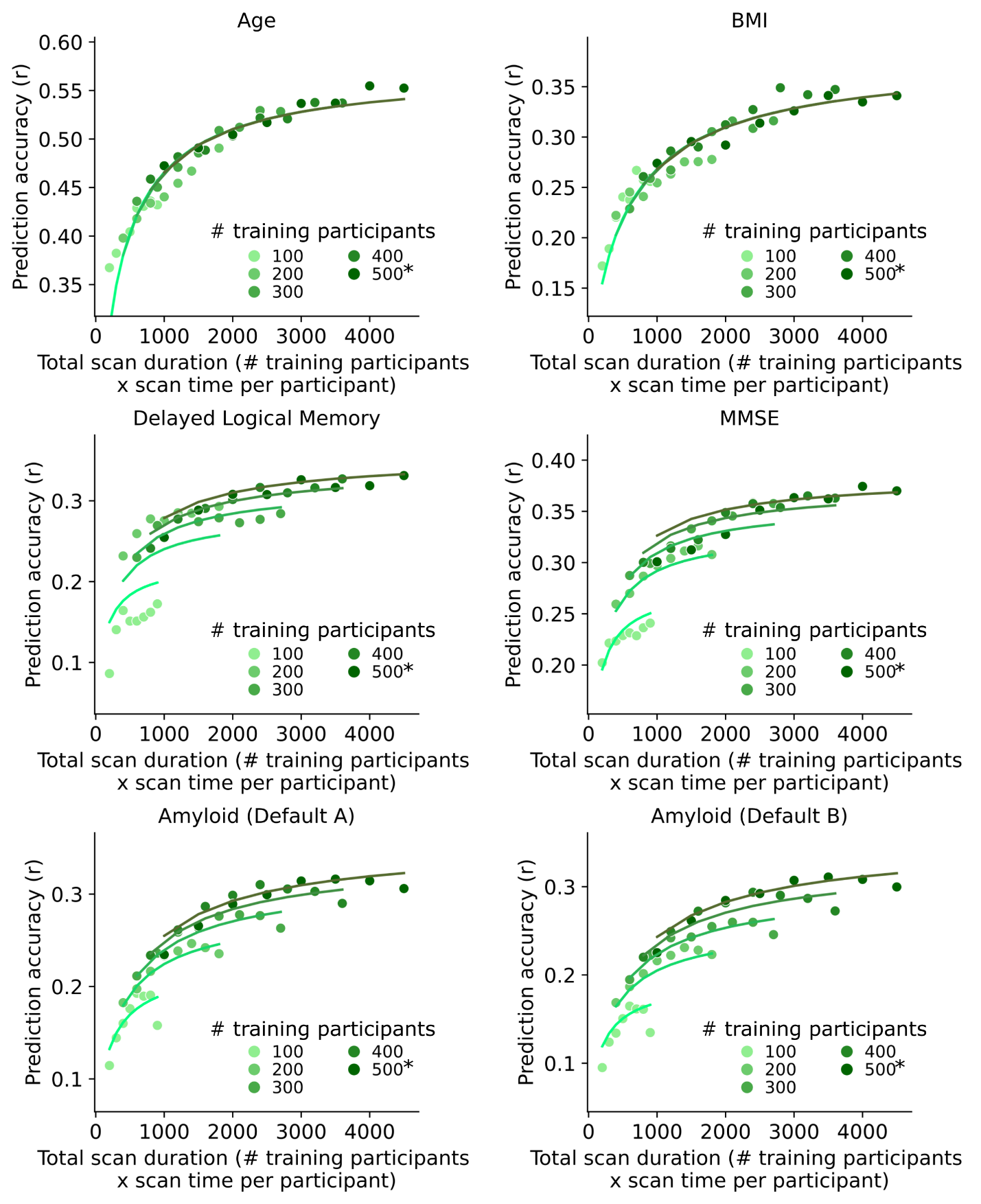

##### **Supplementary Fig. 17 |** Same as Fig. 3b except showing the scatter plots and the fit of prediction accuracy theoretical models for 6 of 6 phenotypic measures in the Alzheimer’s Disease Neuroimaging Initiative (ADNI) dataset. The curves were obtained by fitting a theoretical model to the prediction accuracies of the phenotype. The * in the figures indicates that all available participants were used, therefore the sample size will be close to, but not exactly the number shown.

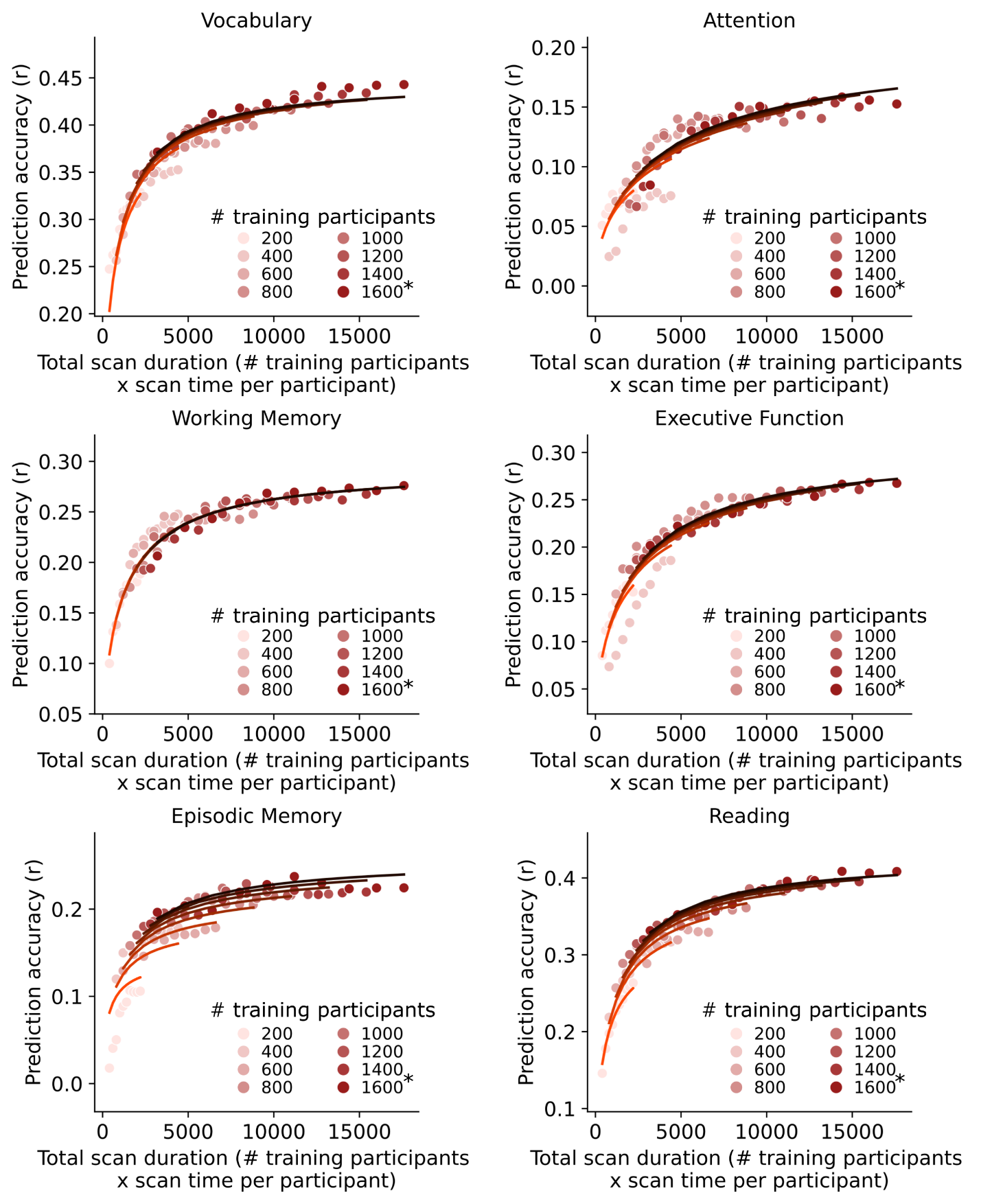

##### **Supplementary Fig. 18.1 |** Same as Fig. 3b except showing the scatter plots and the fit of prediction accuracy theoretical models for 6 of 16 phenotypic measures using Monetary Incentive Delay (MID) task-FC in the ABCD dataset. The curves were obtained by fitting a theoretical model to the prediction accuracies of the phenotype. We only showed phenotypes that exhibited good fit with the theoretical model (Table S2). The * in the figures indicates that all available participants were used, therefore the sample size will be close to, but not exactly the number shown.

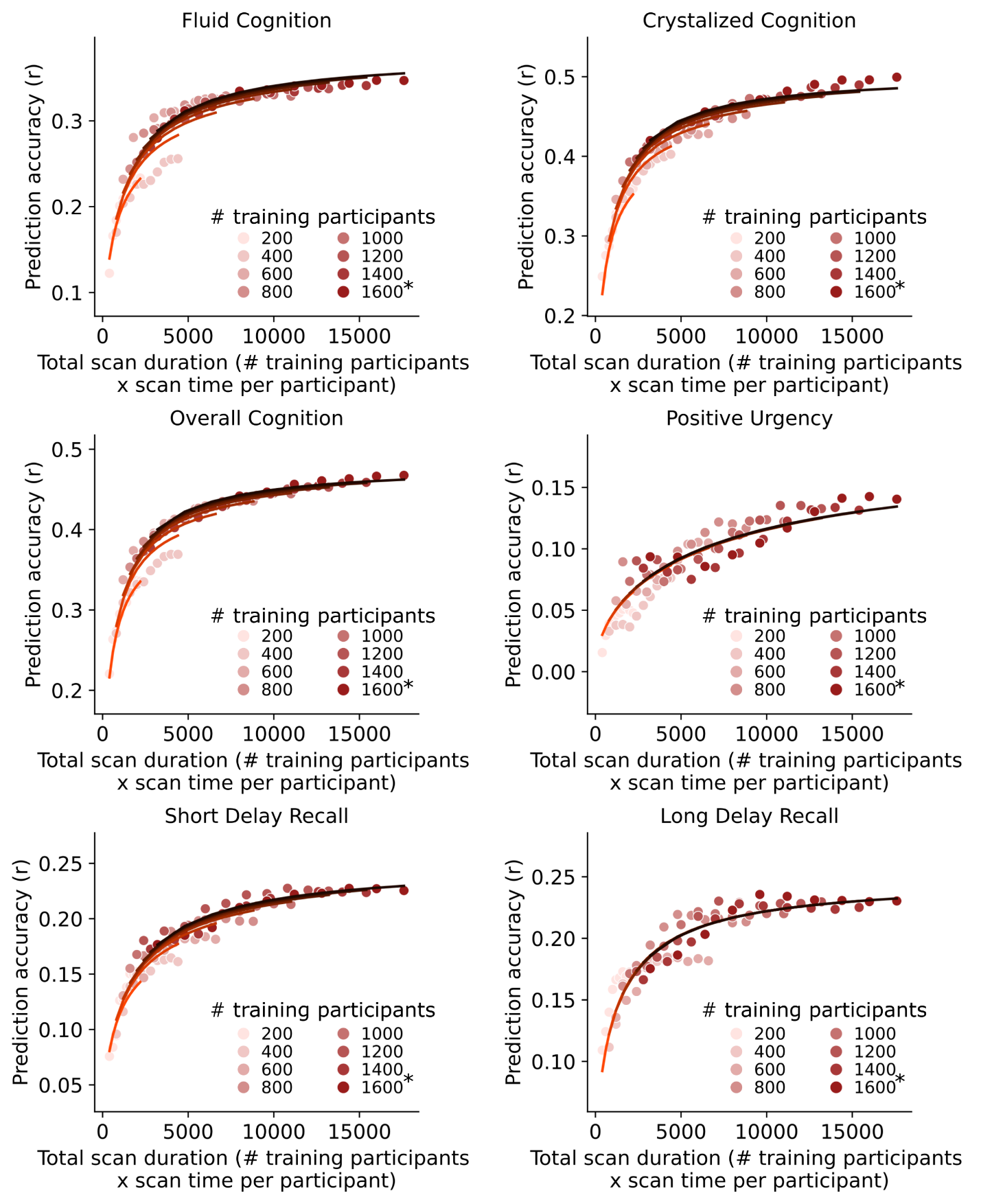
**Supplementary Fig. 18.2 |** Same as Fig. 3b except showing the scatter plots and the fit of prediction accuracy theoretical models for 6 of 16 phenotypic measures using Monetary Incentive Delay (MID) task-FC in the ABCD dataset. The curves were obtained by fitting a theoretical model to the prediction accuracies of the phenotype. We only showed phenotypes that exhibited good fit with the theoretical model (Table S2). The * in the figures indicates that all available participants were used, therefore the sample size will be close to, but not exactly the number shown.

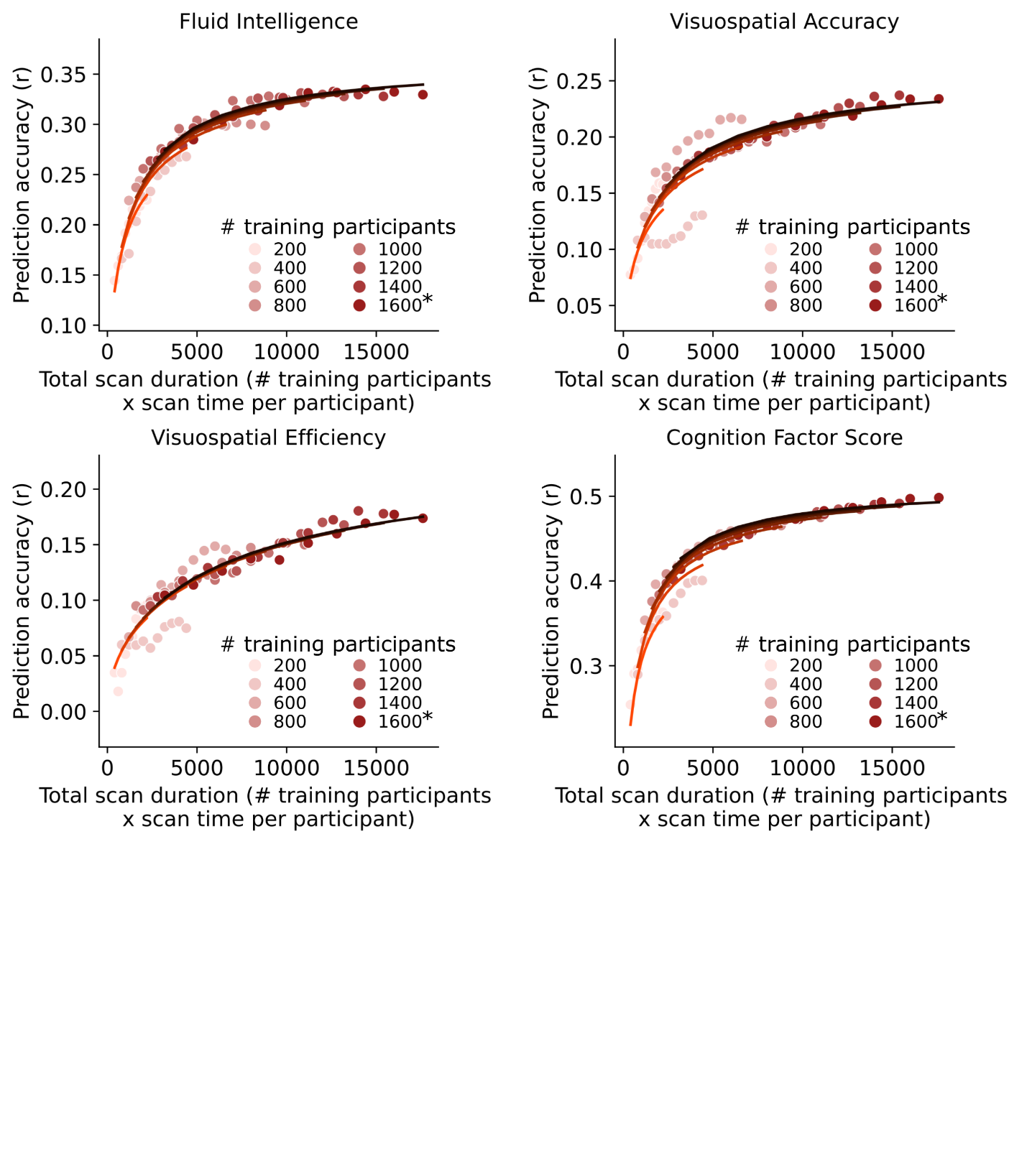

**Supplementary Fig. 18.3 |** Same as Fig. 3b except showing the scatter plots and the fit of prediction accuracy theoretical models for 4 of 16 phenotypic measures using Monetary Incentive Delay (MID) task-FC in the ABCD dataset. The curves were obtained by fitting a theoretical model to the prediction accuracies of the phenotype. We only showed phenotypes that exhibited good fit with the theoretical model (Table S2). The * in the figures indicates that all available participants were used, therefore the sample size will be close to, but not exactly the number shown.

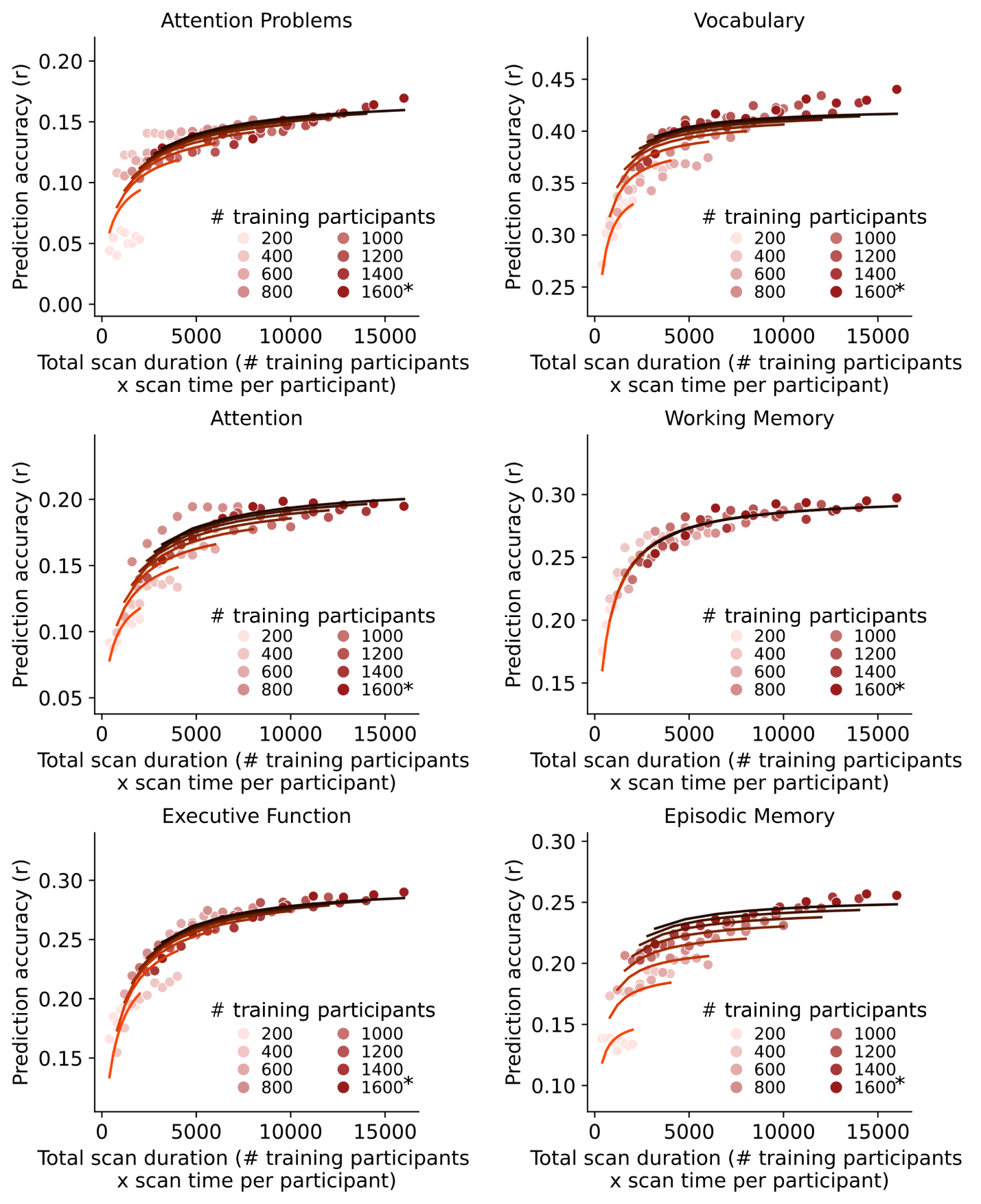
**Supplementary Fig. 19.1 |** Same as Fig. 3b except showing the scatter plots and the fit of prediction accuracy theoretical models for 6 of 19 phenotypic measures for the N-Back Task in the ABCD dataset. The curves were obtained by fitting a theoretical model to the prediction accuracies of the phenotype. We only showed phenotypes that exhibited good fit with the theoretical model (Table S2). The * in the figures indicates that all available participants were used, therefore the sample size will be close to, but not exactly the number shown.

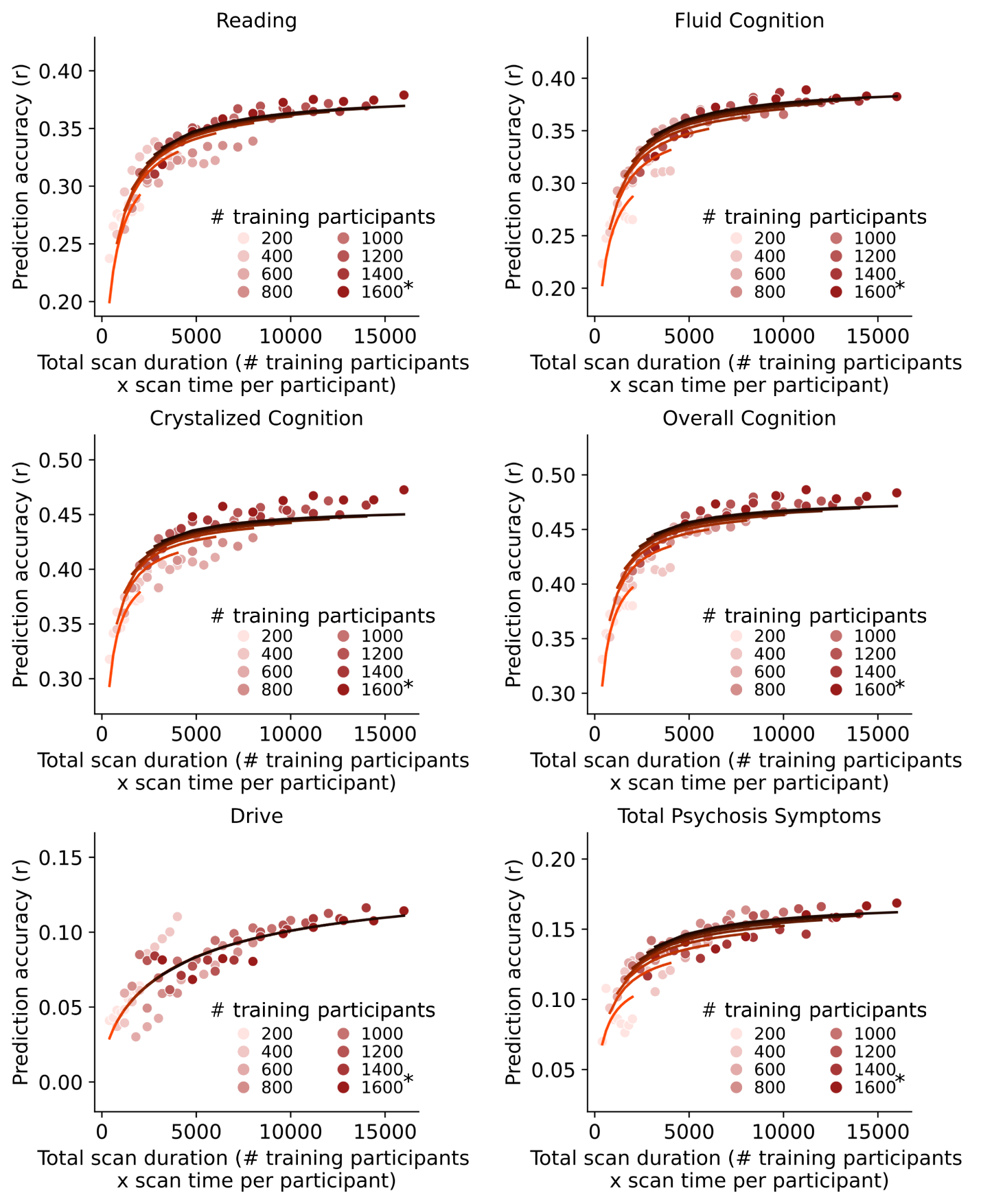

**Supplementary Fig. 19.2 |** Same as Fig. 3b except showing the scatter plots and the fit of prediction accuracy theoretical models for 6 of 19 phenotypic measures for the N-Back Task in the ABCD dataset. The curves were obtained by fitting a theoretical model to the prediction accuracies of the phenotype. We only showed phenotypes that exhibited good fit with the theoretical model (Table S2). The * in the figures indicates that all available participants were used, therefore the sample size will be close to, but not exactly the number shown.

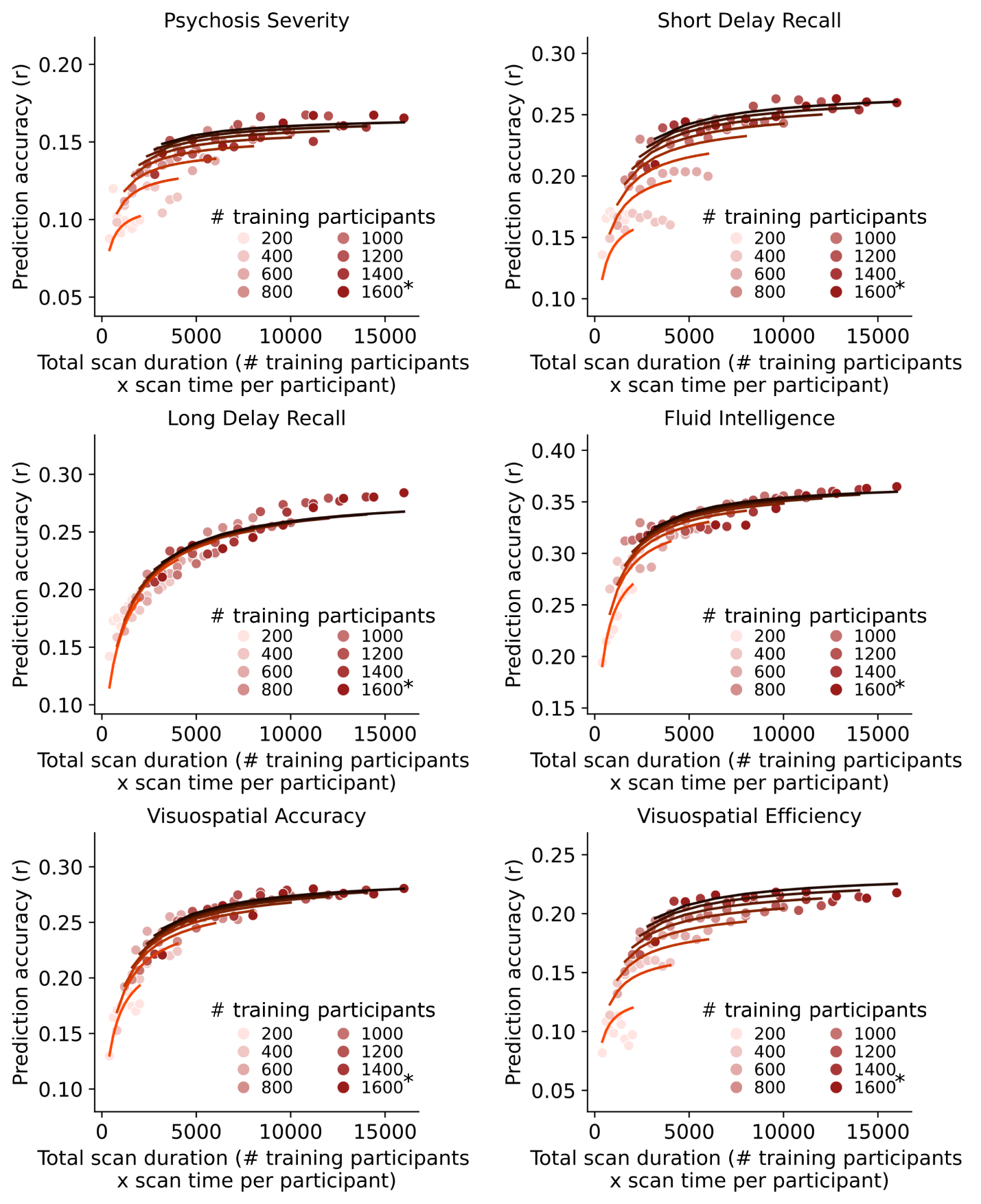

**Supplementary Fig. 19.3 |** Same as Fig. 3b except showing the scatter plots and the fit of prediction accuracy theoretical models for 6 of 19 phenotypic measures for the N-Back Task in the ABCD dataset. The curves were obtained by fitting a theoretical model to the prediction accuracies of the phenotype. We only showed phenotypes that exhibited good fit with the theoretical model (Table S2). The * in the figures indicates that all available participants were used, therefore the sample size will be close to, but not exactly the number shown.

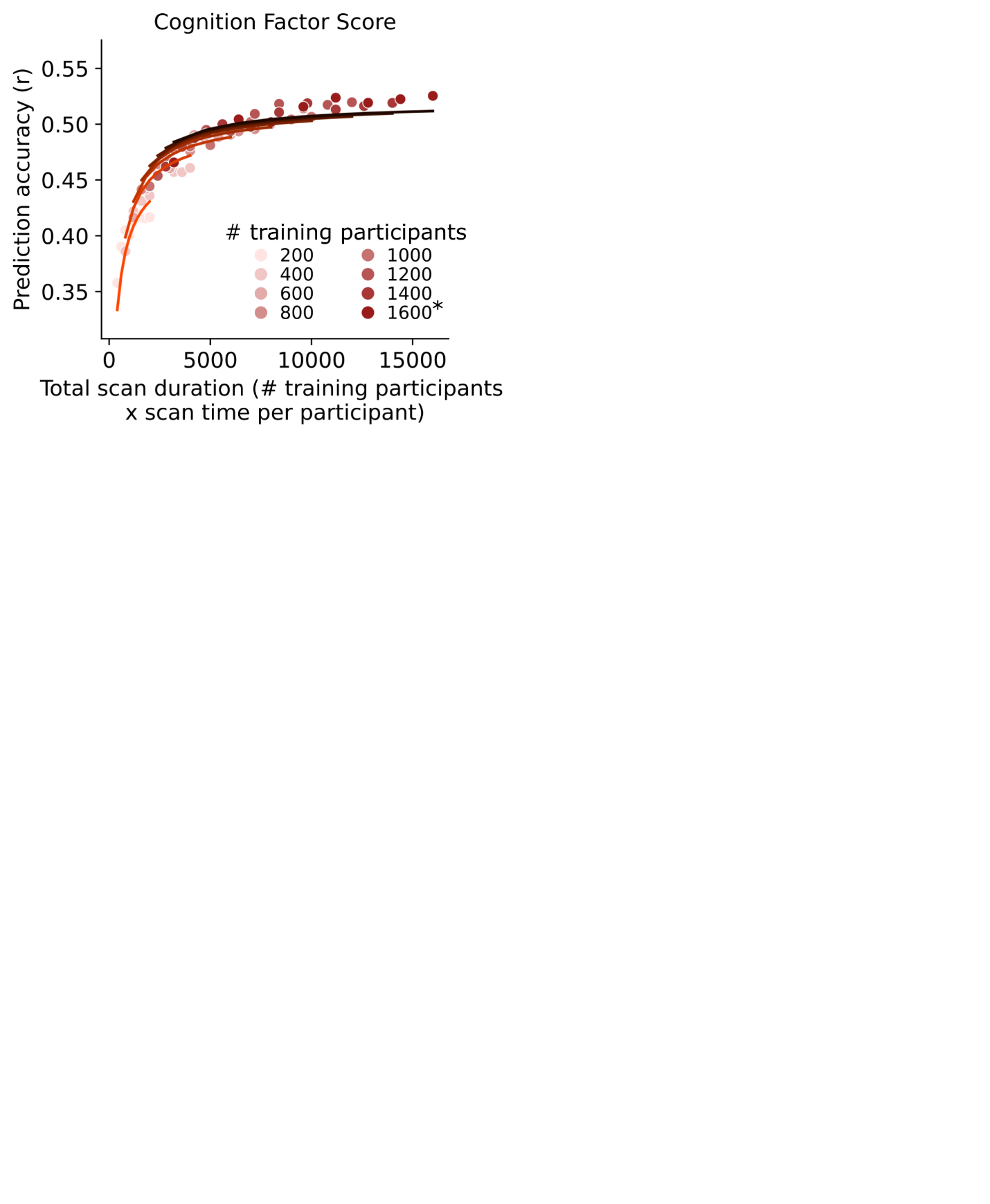

**Supplementary Fig. 19.4 |** Same as Fig. 3b except showing the scatter plots and the fit of prediction accuracy theoretical models for 1 of 19 phenotypic measures for the N-Back Task in the ABCD dataset. The curves were obtained by fitting a theoretical model to the prediction accuracies of the phenotype. We only showed phenotypes that exhibited good fit with the theoretical model (Table S2). The * in the figures indicates that all available participants were used, therefore the sample size will be close to, but not exactly the number shown.

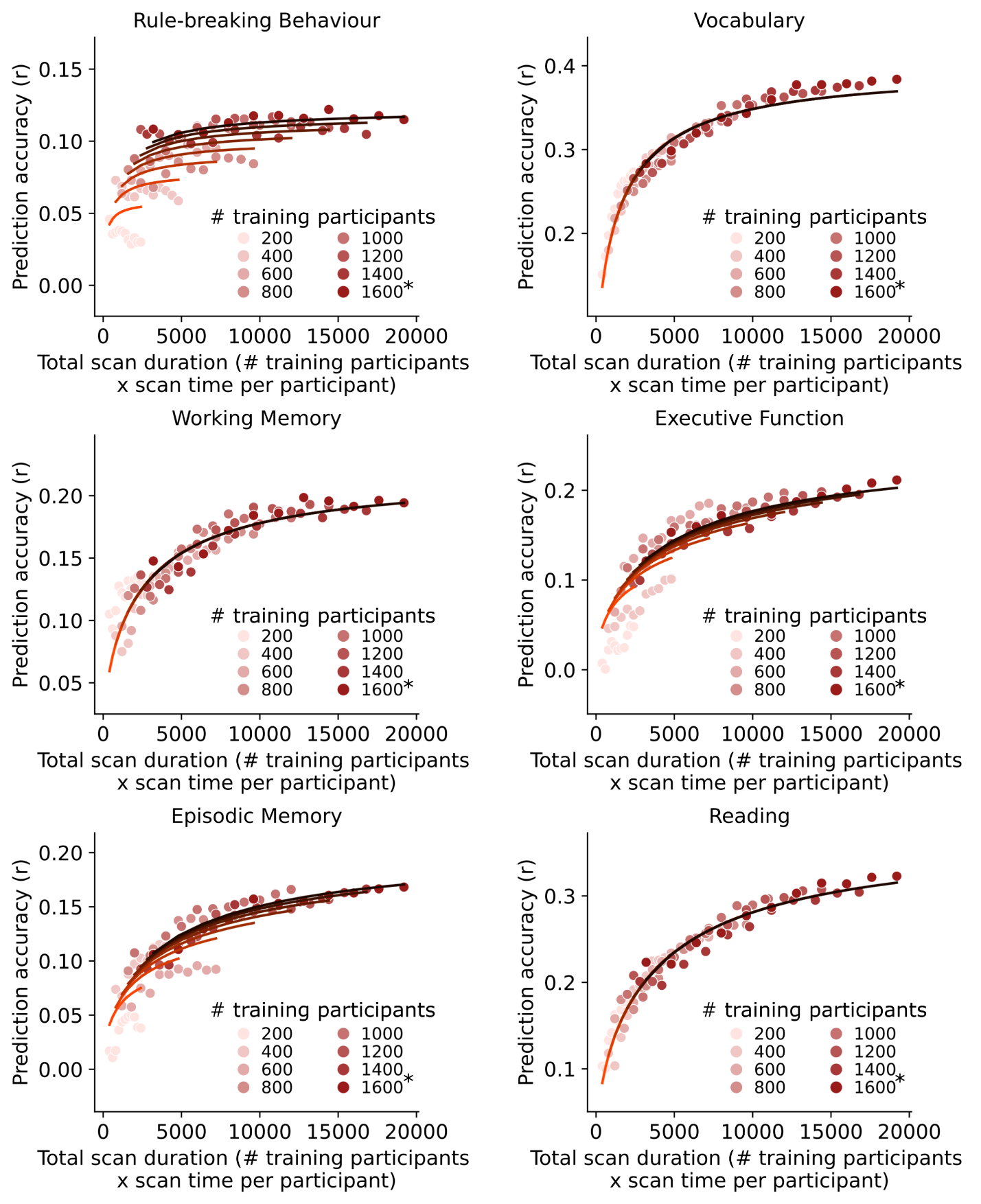

##### **Supplementary Fig. 20.1 |** Same as Fig. 3b except showing the scatter plots and the fit of prediction accuracy theoretical models for 6 of 18 phenotypic measures for the Stop Signal Task (SST) in the ABCD dataset. The curves were obtained by fitting a theoretical model to the prediction accuracies of the phenotype. We only showed phenotypes that exhibited good fit with the theoretical model (Table S2). The * in the figures indicates that all available participants were used, therefore the sample size will be close to, but not exactly the number shown.

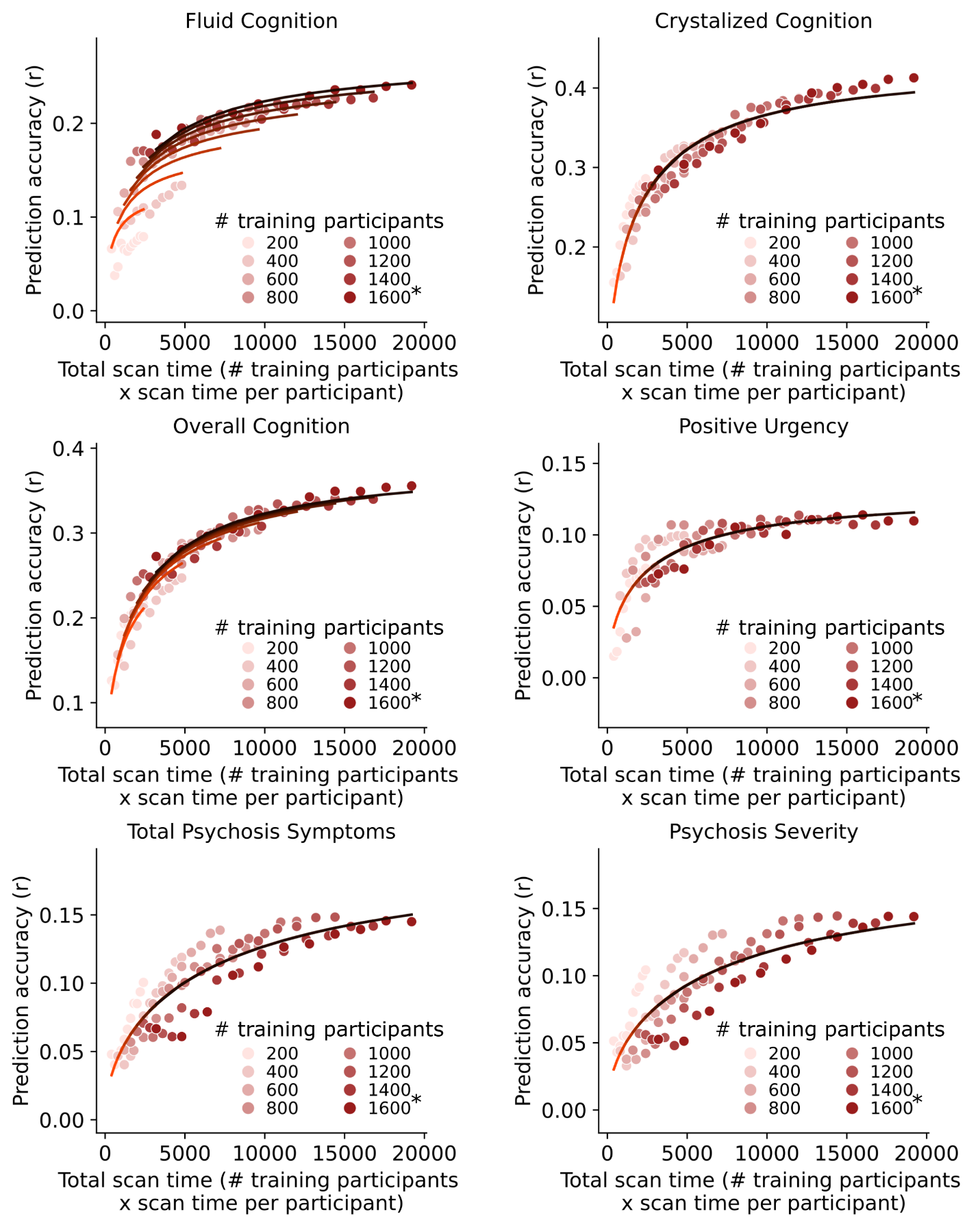
**Supplementary Fig. 20.2 |** Same as Fig. 3b except showing the scatter plots and the fit of prediction accuracy theoretical models for 6 of 18 phenotypic measures for the Stop Signal Task (SST) in the ABCD dataset. The curves were obtained by fitting a theoretical model to the prediction accuracies of the phenotype. We only showed phenotypes that exhibited good fit with the theoretical model (Table S2). The * in the figures indicates that all available participants were used, therefore the sample size will be close to, but not exactly the number shown.

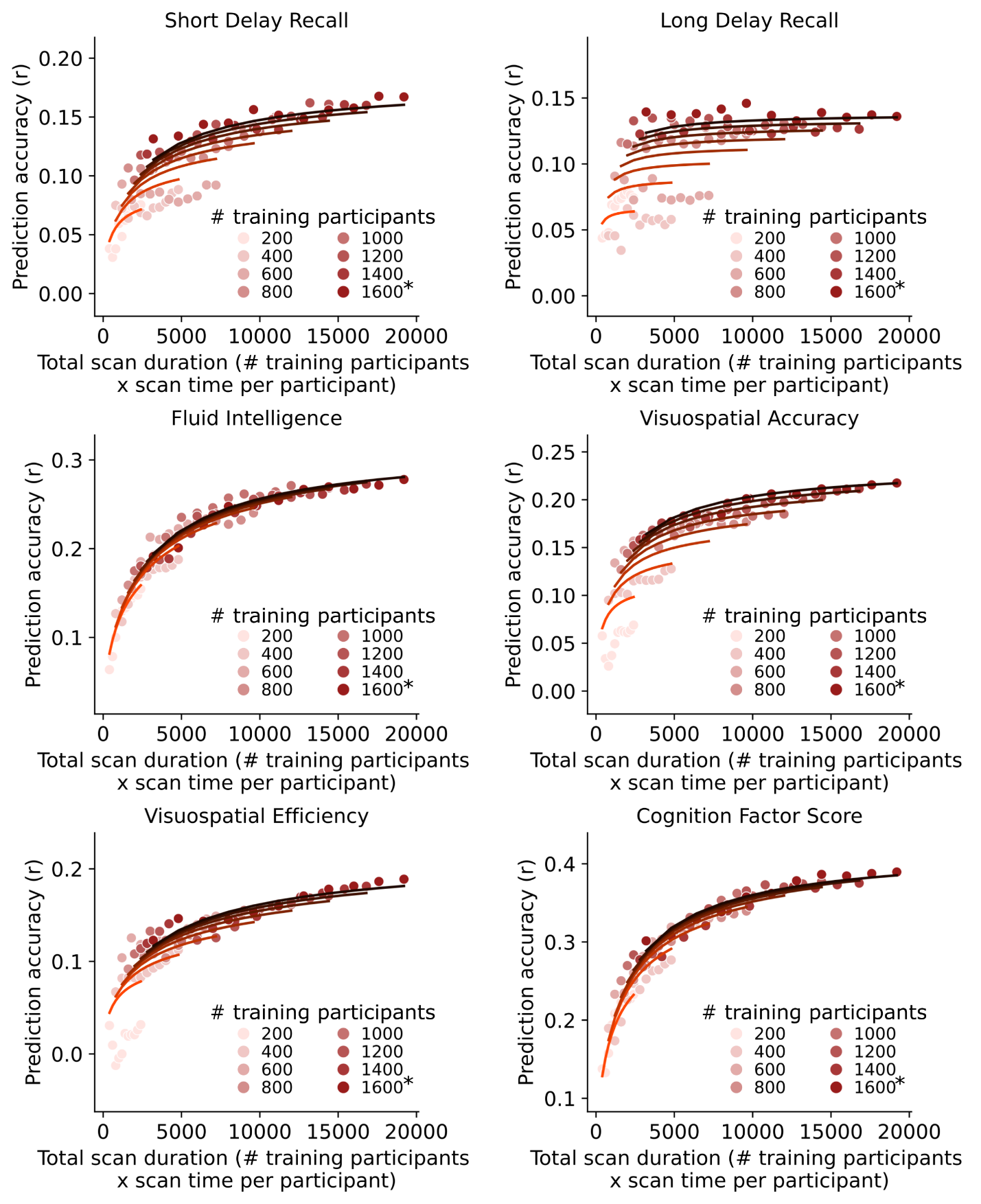
**Supplementary Fig. 20.3 |** Same as Fig. 3b except showing the scatter plots and the fit of prediction accuracy theoretical models for 6 of 18 phenotypic measures for the Stop Signal Task (SST) in the ABCD dataset. The curves were obtained by fitting a theoretical model to the prediction accuracies of the phenotype. We only showed phenotypes that exhibited good fit with the theoretical model (Table S2). The * in the figures indicates that all available participants were used, therefore the sample size will be close to, but not exactly the number shown.

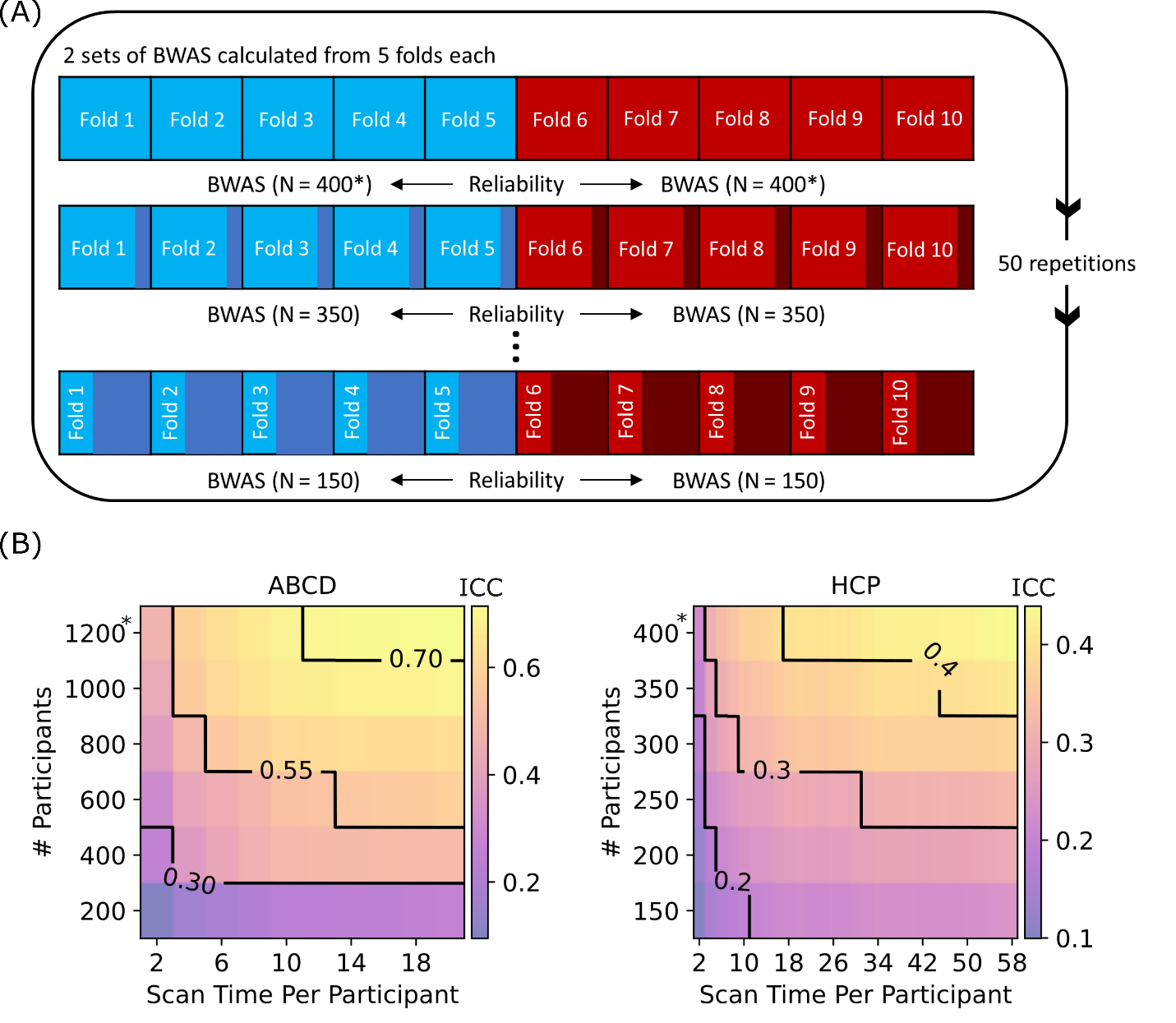

##### **Supplementary Fig. 21 | a.** Reliability analysis workflow for the HCP dataset. The participants were split into 2 sets. Univariate brain-wide association (BWAS) was performed on each set and the agreement (i.e., split-half reliability) between the two sets was computed based on the intra-class correlation metric (see Methods). To vary sample size, each set was subsampled and the whole procedure was repeated. Finally, the procedure was repeated with different amount of fMRI data *T* (not shown in panel) and 50 times for stability. A similar workflow was used in the ABCD dataset. Similar to the prediction analysis (Extended Data Fig. 1), in the case of HCP, care was taken so siblings were not split across sets, while in the case of ABCD, participants from the same site were not split across sets. See Methods for details. **b.** Contour plot of univariate brain-wide association analyses (BWAS) reliability (intra-class correlation) of the cognitive factor score as a function of the scan time used to generate the functional connectivity matrix (x-axis), and the number of training participants used to train the predictive model (y-axis) in the ABCD and HCP datasets. Increasing training participants and scan time both led to increases in split-half reliability. The * in both figures indicates that all available participants were used, therefore the sample size will be close to, but not exactly the number shown.

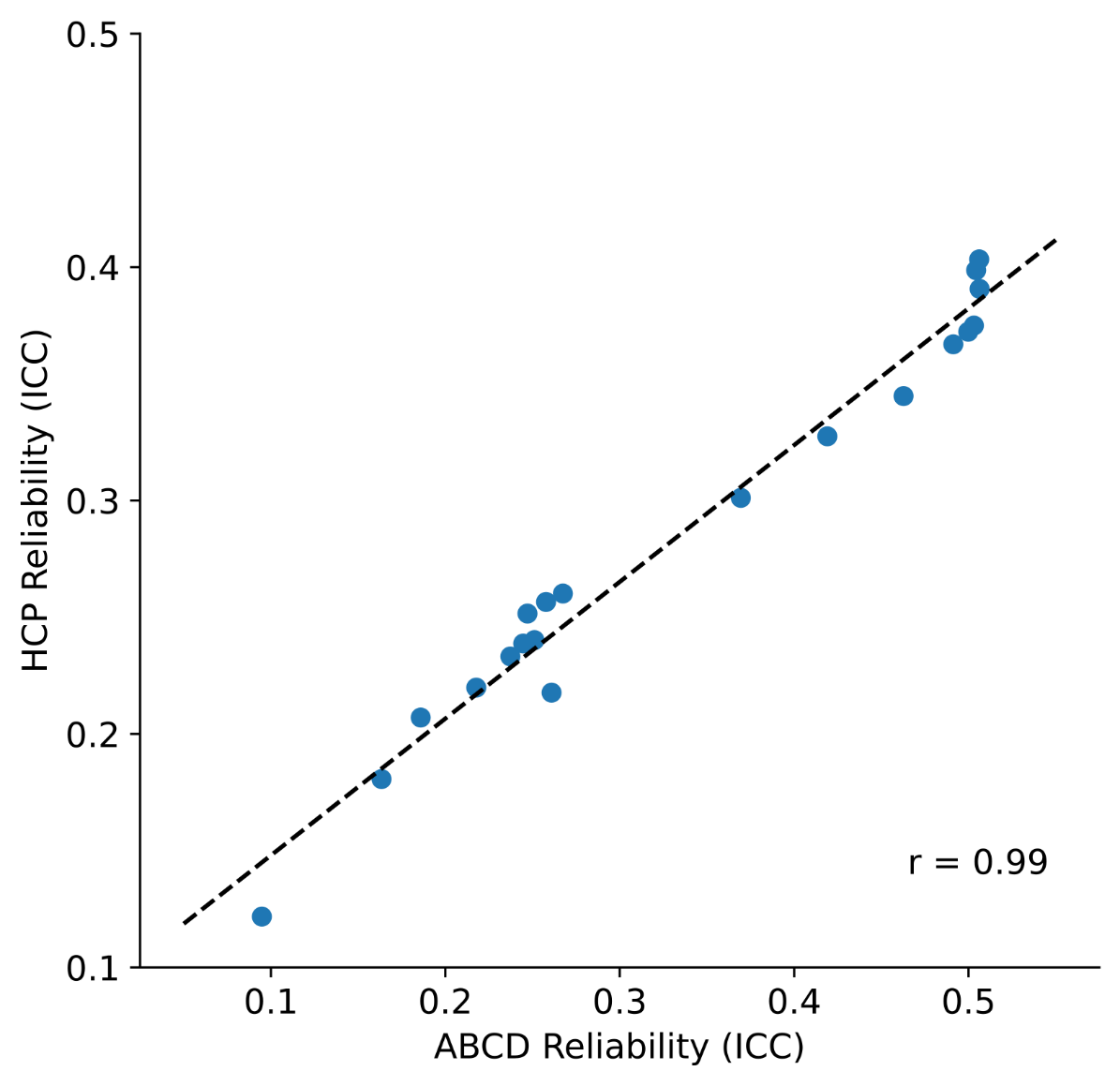

##### **Supplementary Fig. 22** | Scatter plot of the cognition factor univariate reliability (ICC) in the ABCD (x-axis) and HCP (y-axis) datasets. Each dot represents the prediction accuracy for each dataset with the same sample size and scan time per participant (extracted from Supplementary Fig. 21b). Although the cognitive factor score is not comparable across datasets, we observed a strong correlation between the two datasets (r = 0.99).

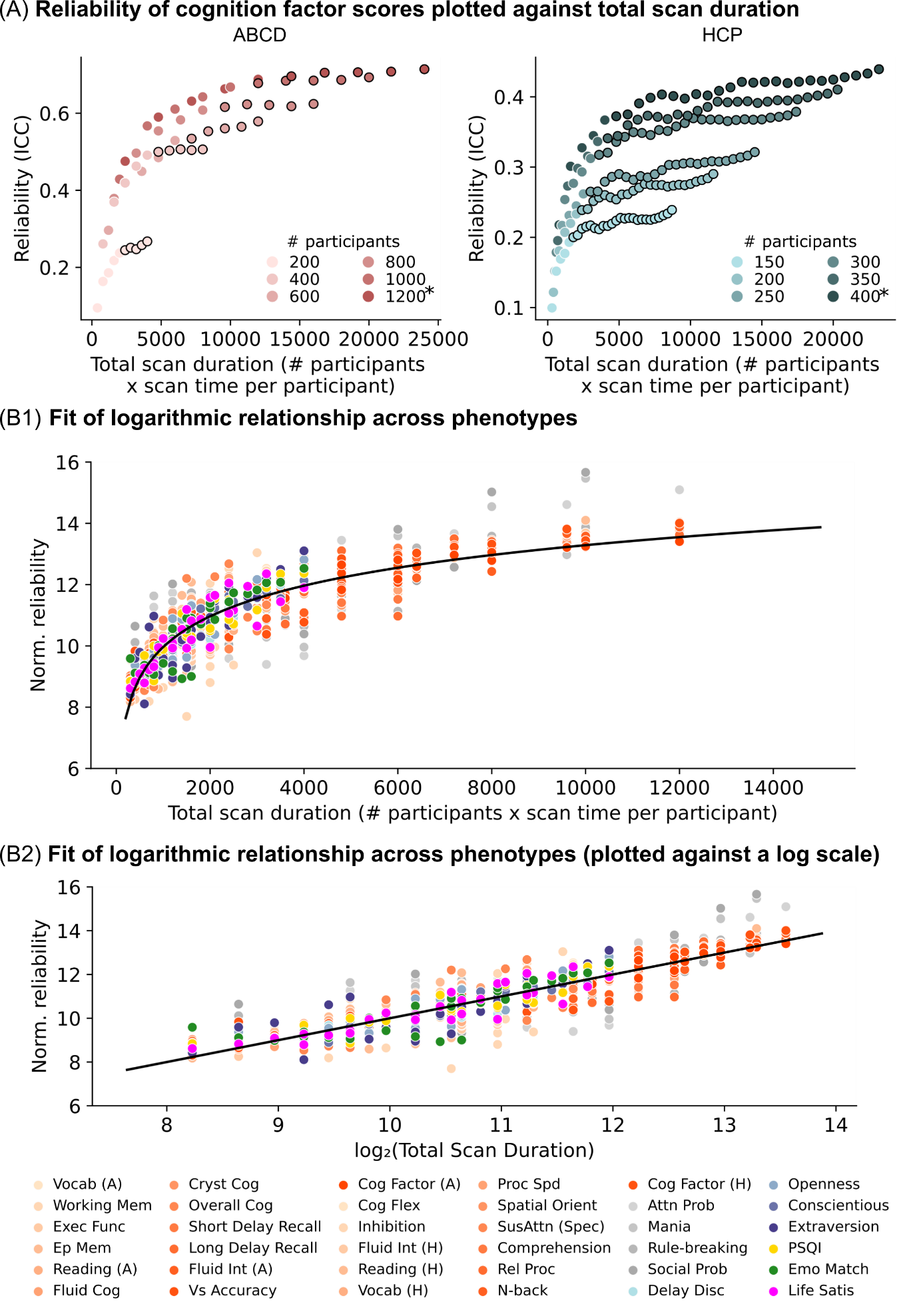

##### **Supplementary Fig. 23 | a.** Scatter plot showing reliability of univariate brain-wide association (intra-class correlation) of the cognitive factor as a function of total scan duration (defined as # training participants x scan time per participant). Each color represents a different number of total participants used to train the prediction algorithm. Plots were repeated for ABCD and HCP datasets. The * indicates that all available participants were used, therefore the sample size will be close to, but not exactly the number shown. Increase in reliability observed diminishing returns (relative to sample size) when scan time per participant reached approximately 10 minutes in the HCP dataset; data points with more than 10 minutes of scan time are shown with black outlines. **b1.** By plotting total scan duration (number of participants × scan time per participant) against univariate BWAS reliability for each phenotype, we observed that for most phenotypes, scanning beyond 10 minutes per participant yielded diminishing marginal returns to reliability. Therefore, we performed the same logarithmic curve fitting procedure as before, but using only up to 10 minutes of scan time per participant. Scatter plot showing normalized reliability of the cognitive factor scores and 34 other phenotypes versus total scan duration ignoring data beyond 10 minutes of scan time. Blue and red dots represent results from the HCP and the ABCD datasets respectively. The logarithmic black curve suggests that total scan duration explained reliability well across phenotypic domains and datasets. **b2.** Same as panel b1, except the horizontal axis (total scan duration) is plotted on a logarithm scale. The linear black line suggests that the logarithm of total scan duration explained prediction performance well across phenotypic domains and datasets.

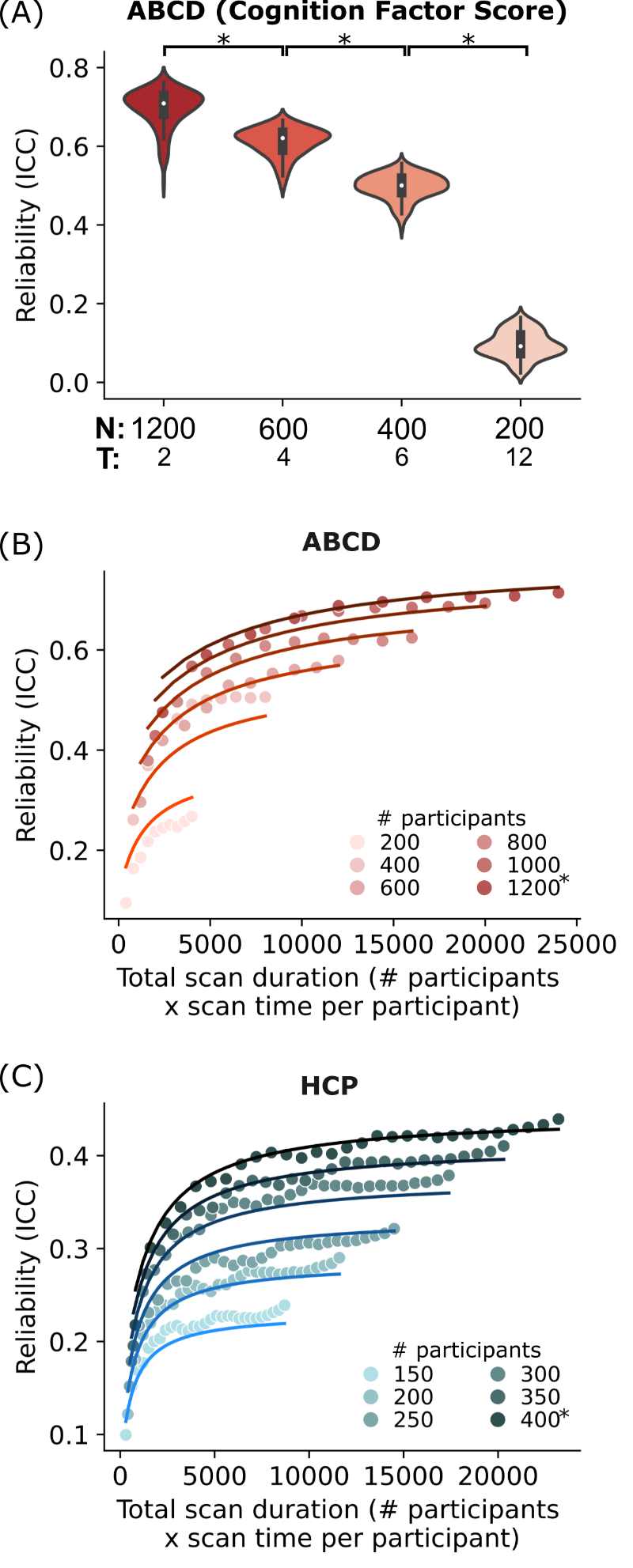

##### **Supplementary Fig. 24 |** Sample size and scan time are not 1-to-1 interchangeable **a.** Each violin shows the distribution of univariate brain-wide association analyses (BWAS) split-half reliability for the Adolescent Brain and Cognitive Development (ABCD) cognition factor score across 126 unique site combinations for a given set of scan parameters. All violins have the same total scan duration of 2400 minutes. * indicate that the distribution of reliabilities were significantly different (after FDR correction). Having a larger sample size is more beneficial for reliability. **b.** Scatter plot of reliability against total scan duration in the ABCD dataset. The curves were obtained by fitting a theoretical model to the reliabilities of the cognitive factor score that explains the contribution of sample size and scan time (see Supplementary methods 1.3). We fitted the function ${Rel}_{p}=\frac{K_{0,p}}{K_{0,p}+\frac{1}{\frac{N}{2}}\left( 1-2K_{1,b}\left( \frac{1}{1+\frac{K_{2,p}}{T}} \right) \right)}$ to the data, where ${Rel}_{p}$ was the univariate split-half reliability (in terms of ICC) for phenotypic measure $p$, $N$ is the sample size and $T$ is the scan time per participant. $K_{0,p}, K_{1,p}$ and $K_{2,p}$ were estimated from data through a gradient descent, minimizing the mean squared error for each phenotypic measure. The theoretical model was able to explain why sample size is more important than scan time. **c.** Same as panel b but for the Human Connectome Project (HCP) dataset.

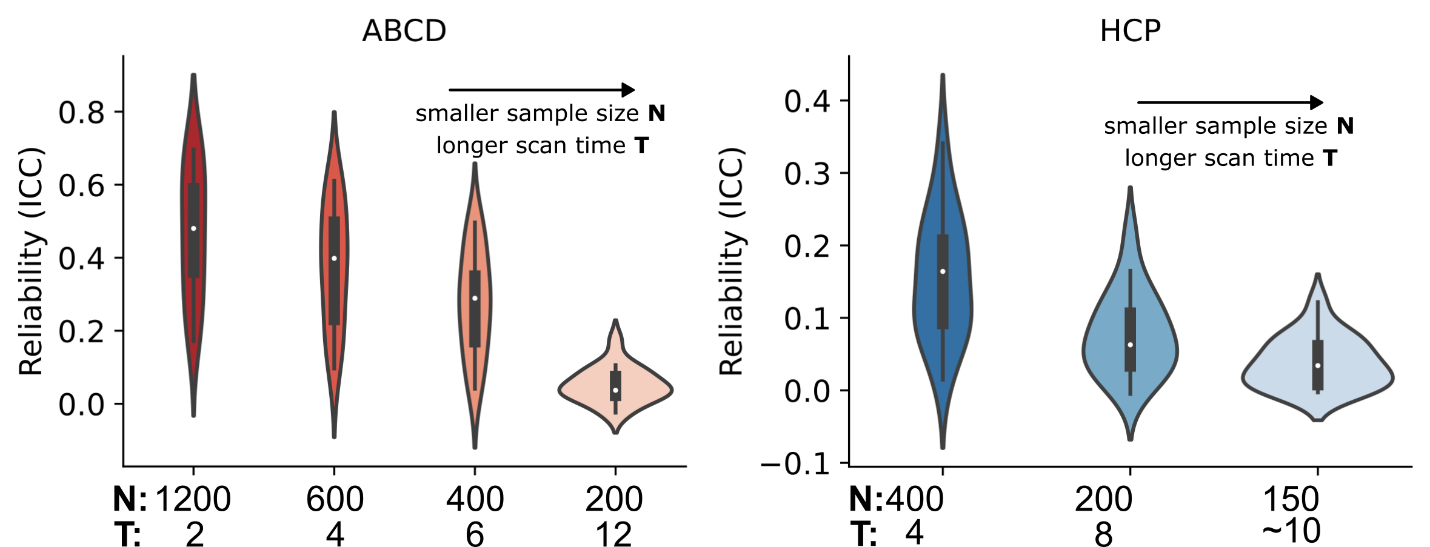

##### **Supplementary Fig. 25 |** Same as Supplementary Fig.24a, except in the left panel, each violin shows the distribution of average univariate brain-wide association analyses (BWAS) split-half reliability across 17 phenotypic measures in the ABCD dataset. Each violin has a total scan duration of 2400 mins (left panel). In the right panel, each violin shows the distribution average BWAS split-half reliability of the 19 HCP phenotypic measures. Each violin Each violin contains 19 data points and has a total scan duration of 1600 mins (right panel).

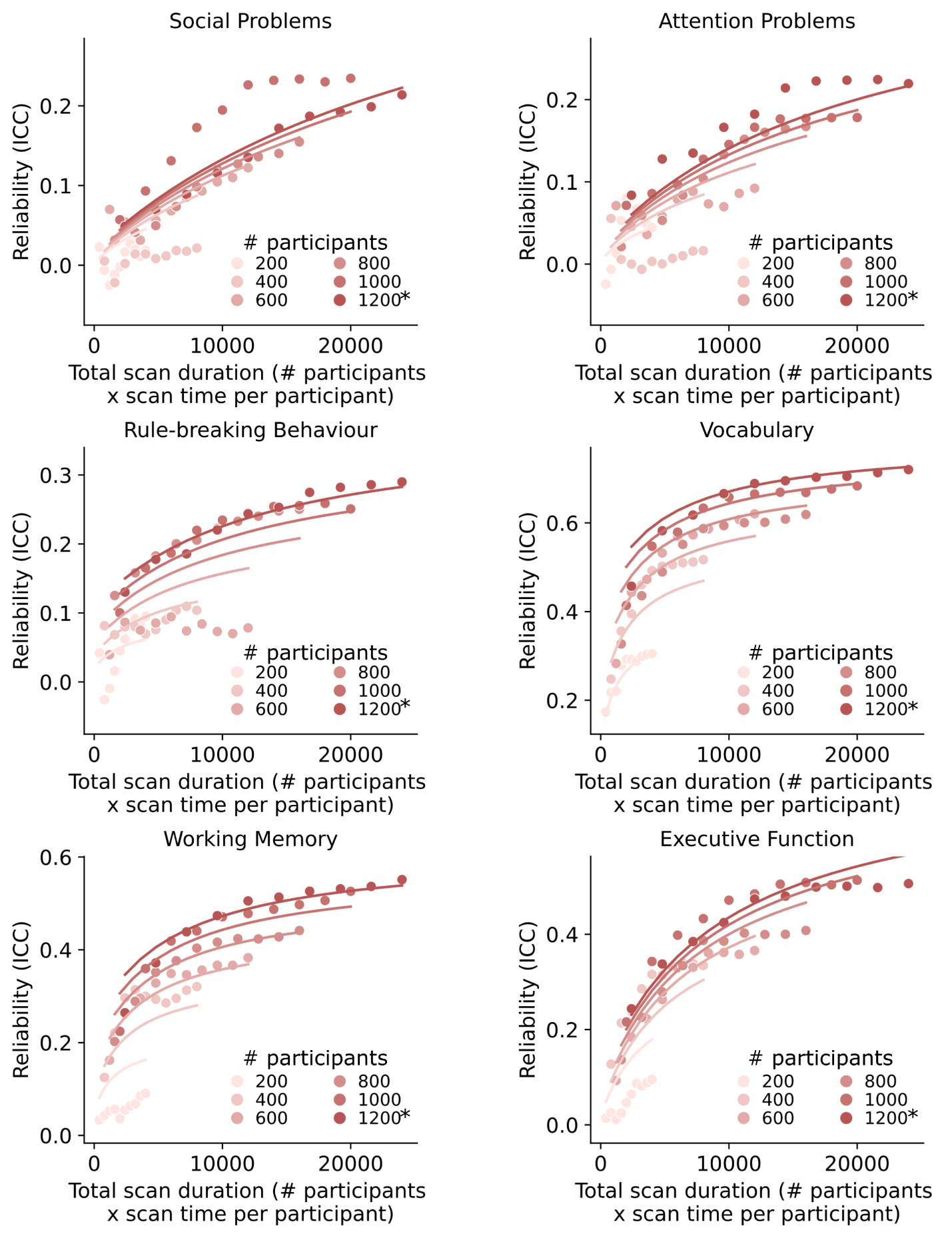

##### **Supplementary Fig. 26.1 |** Same as Supplementary Fig. 24b except showing the scatter plots and the fit of reliability theoretical models for 6 of 17 phenotypic measures in the ABCD dataset that visually follow a logarithmic pattern for prediction accuracy. Scatter plot of split-half univariate brain-wise association analyses reliability (intra-class correlation) against total scan duration in the ABCD dataset. The curves were obtained by fitting a theoretical model to the reliabilities of the phenotype. The * in the figures indicates that all available participants were used, therefore the sample size will be close to, but not exactly the number shown.

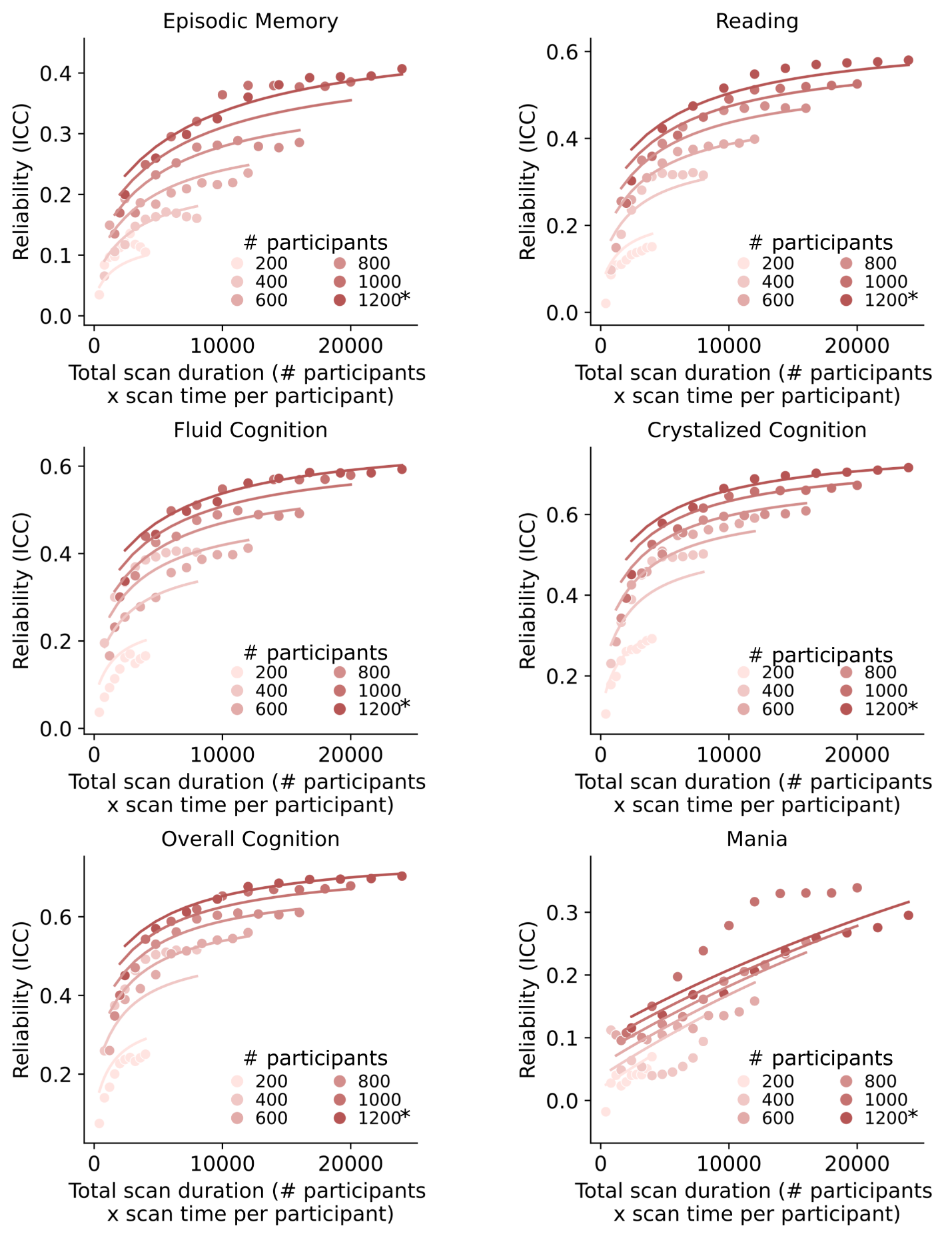

**Supplementary Fig. 26.2 |** Same as Supplementary Fig. 24b except showing the scatter plots and the fit of reliability theoretical models for 6 of 17 phenotypic measures in the ABCD dataset that visually follow a logarithmic pattern for prediction accuracy. Scatter plot of split-half univariate brain-wise association analyses reliability (intra-class correlation) against total scan duration in the ABCD dataset. The curves were obtained by fitting a theoretical model to the reliabilities of the phenotype. The * in the figures indicates that all available participants were used, therefore the sample size will be close to, but not exactly the number shown.

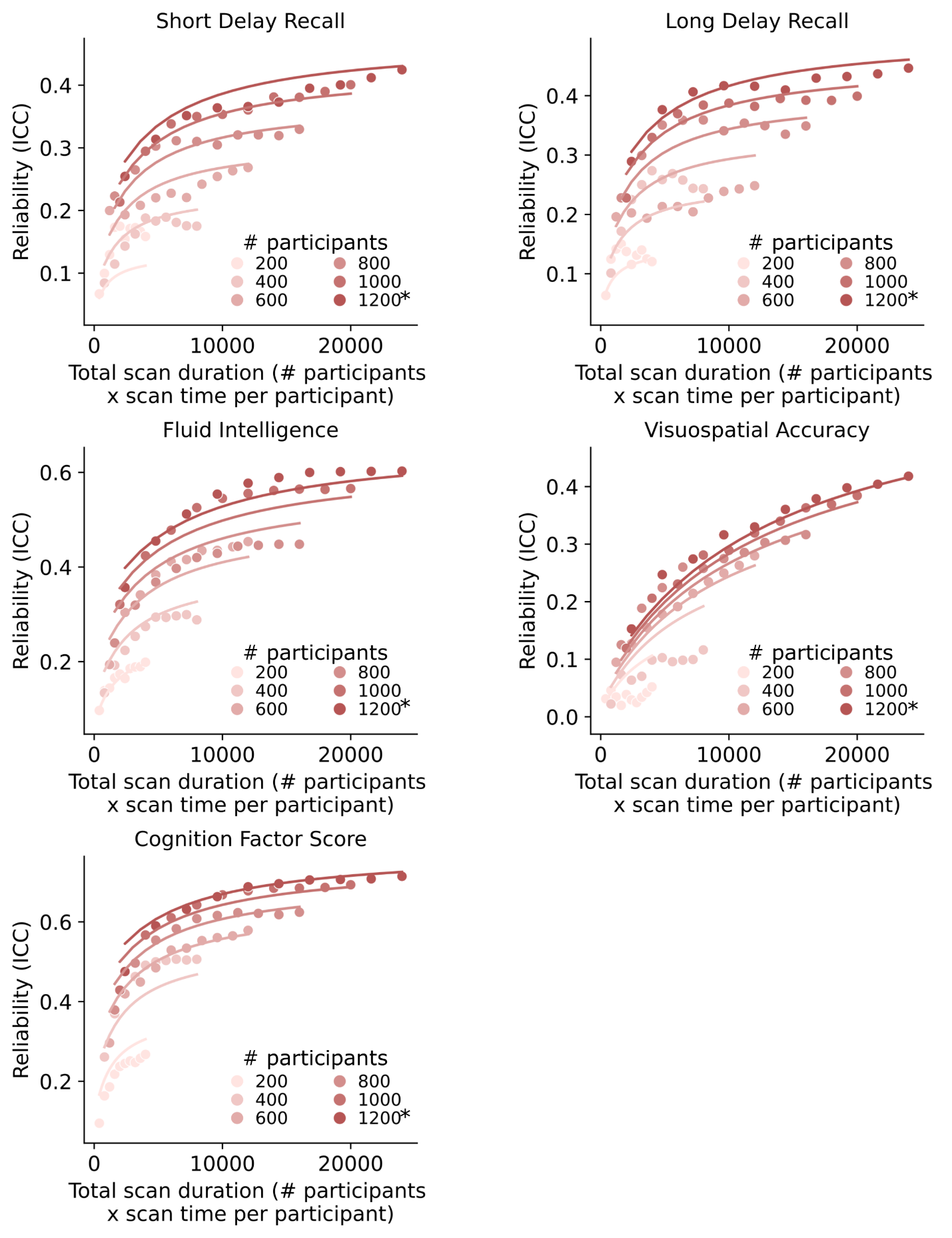

**Supplementary Fig. 26.3 |** Same as Supplementary Fig. 24b except showing the scatter plots and the fit of reliability theoretical models for 5 of 17 phenotypic measures in the ABCD dataset that visually follow a logarithmic pattern for prediction accuracy. Scatter plot of split-half univariate brain-wise association analyses reliability (intra-class correlation) against total scan duration in the ABCD dataset. The curves were obtained by fitting a theoretical model to the reliabilities of the phenotype. The * in the figures indicates that all available participants were used, therefore the sample size will be close to, but not exactly the number shown.

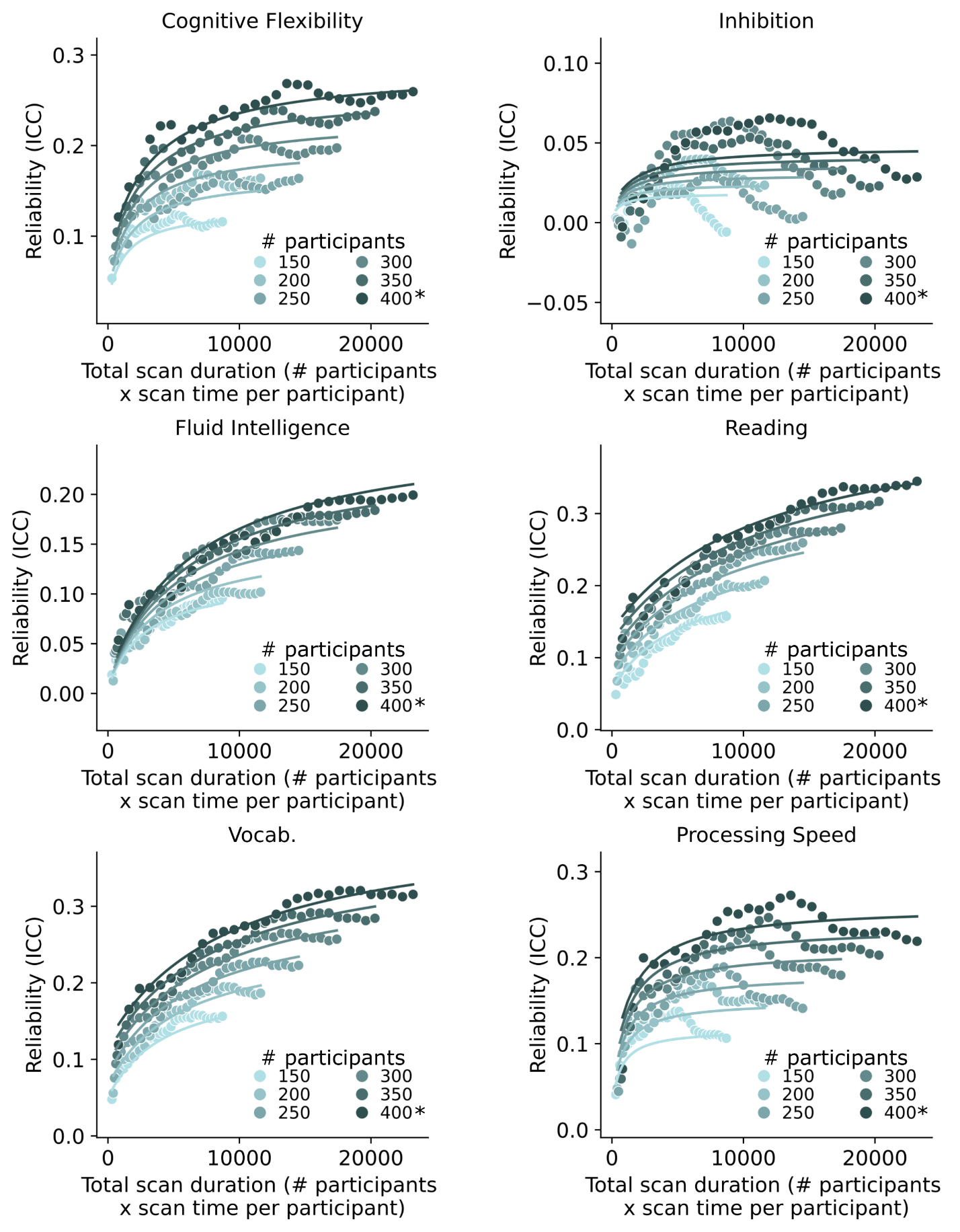

##### **Supplementary Fig. 27.1 |** Same as Supplementary Fig. 24c except showing the scatter plots and the fit of reliability theoretical models for 6 of 19 phenotypic measures in the HCP dataset that visually follow a logarithmic pattern for prediction accuracy. Scatter plot of split-half univariate brain-wise association analyses reliability (intra-class correlation) against total scan duration in the HCP dataset. The curves were obtained by fitting a theoretical model to the reliabilities of the phenotype. The * in the figures indicates that all available participants were used, therefore the sample size will be close to, but not exactly the number shown.

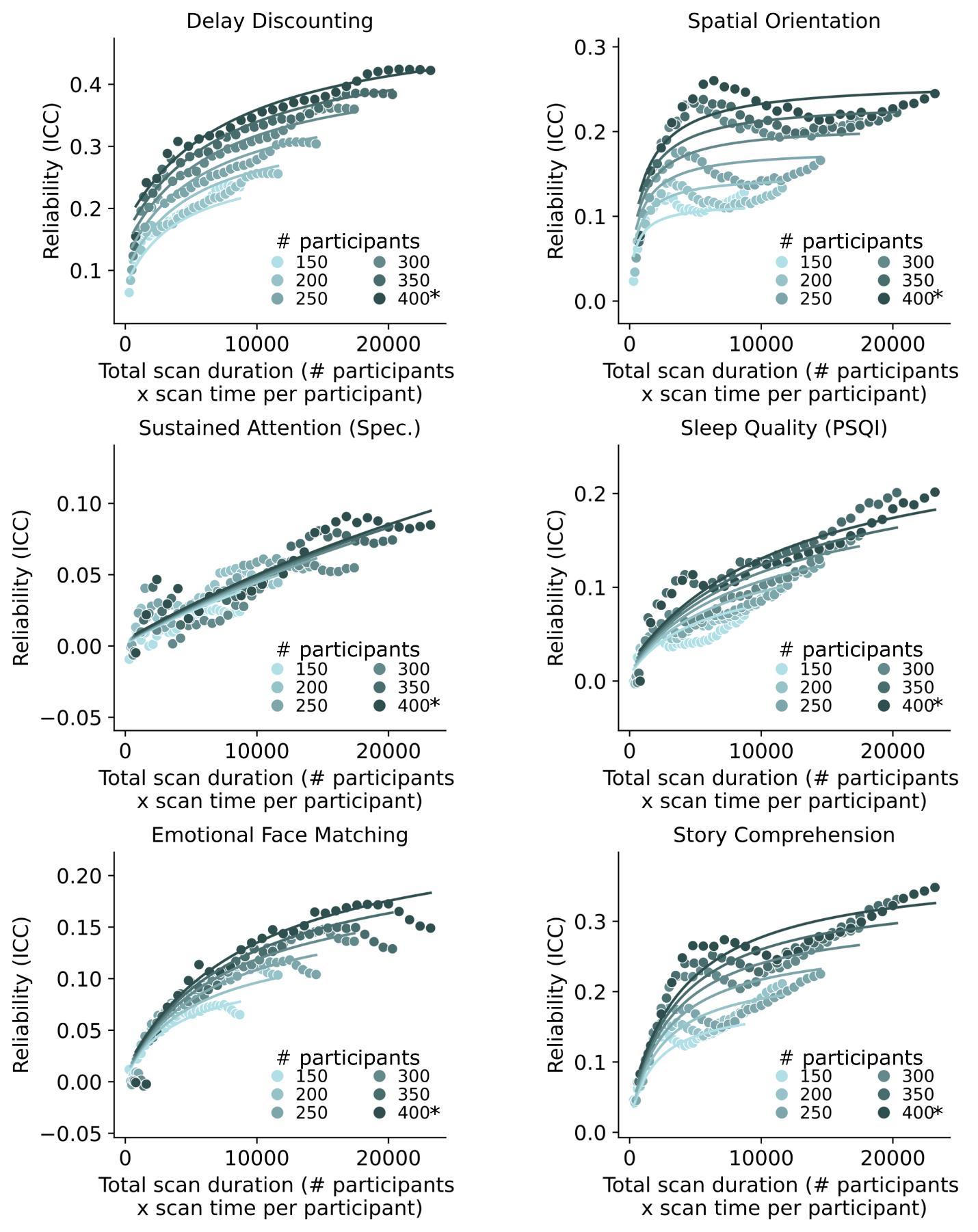
**Supplementary Fig. 27.2 |** Same as Supplementary Fig. 24c except showing the scatter plots and the fit of reliability theoretical models for 6 of 19 phenotypic measures in the HCP dataset that visually follow a logarithmic pattern for prediction accuracy. Scatter plot of split-half univariate brain-wise association analyses reliability (intra-class correlation) against total scan duration in the HCP dataset. The curves were obtained by fitting a theoretical model to the reliabilities of the phenotype. The * in the figures indicates that all available participants were used, therefore the sample size will be close to, but not exactly the number shown.

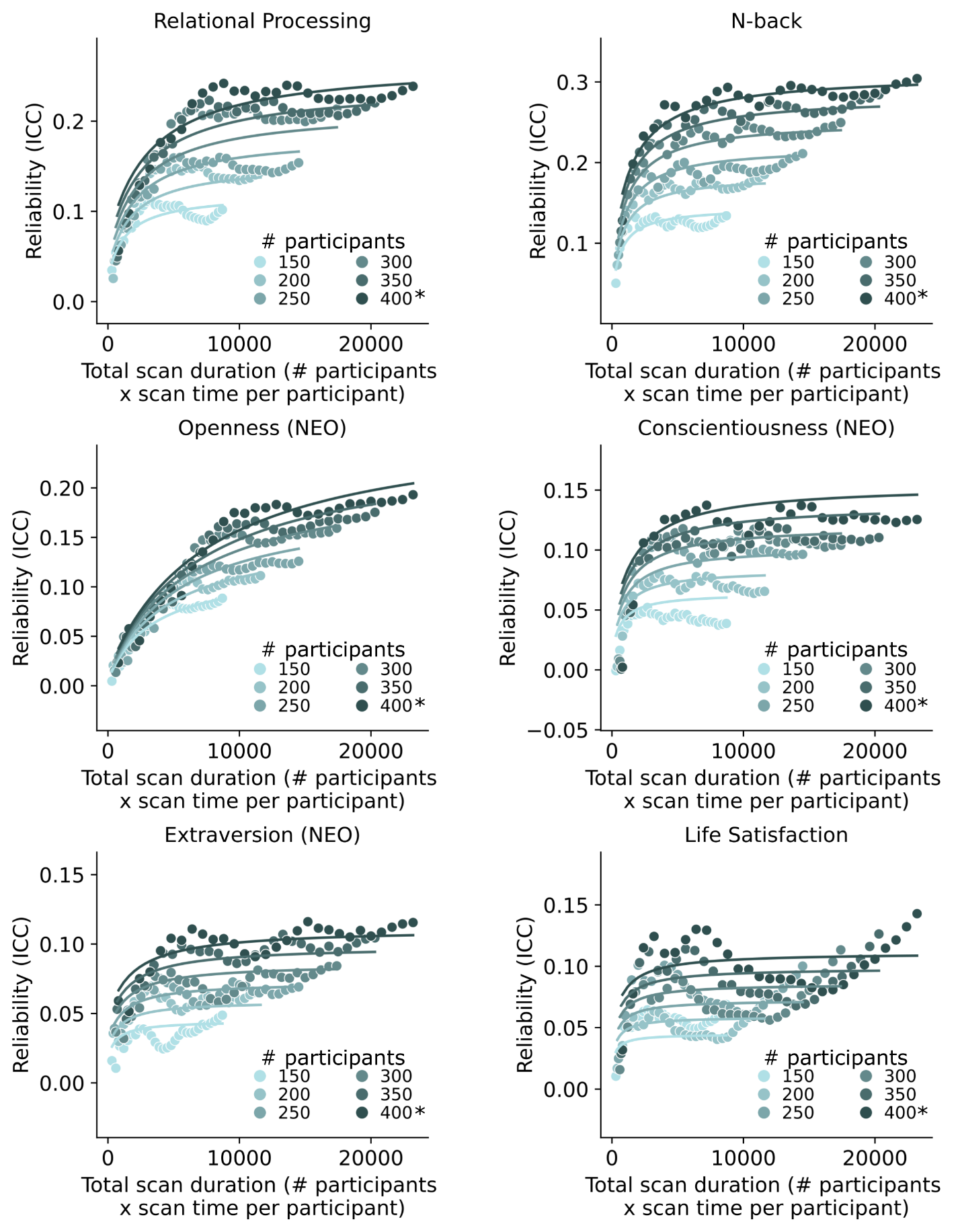

**Supplementary Fig. 27.3 |** Same as Supplementary Fig. 24c except showing the scatter plots and the fit of reliability theoretical models for 6 of 19 phenotypic measures in the HCP dataset that visually follow a logarithmic pattern for prediction accuracy. Scatter plot of split-half univariate brain-wise association analyses reliability (intra-class correlation) against total scan duration in the HCP dataset. The curves were obtained by fitting a theoretical model to the reliabilities of the phenotype. The * in the figures indicates that all available participants were used, therefore the sample size will be close to, but not exactly the number shown.

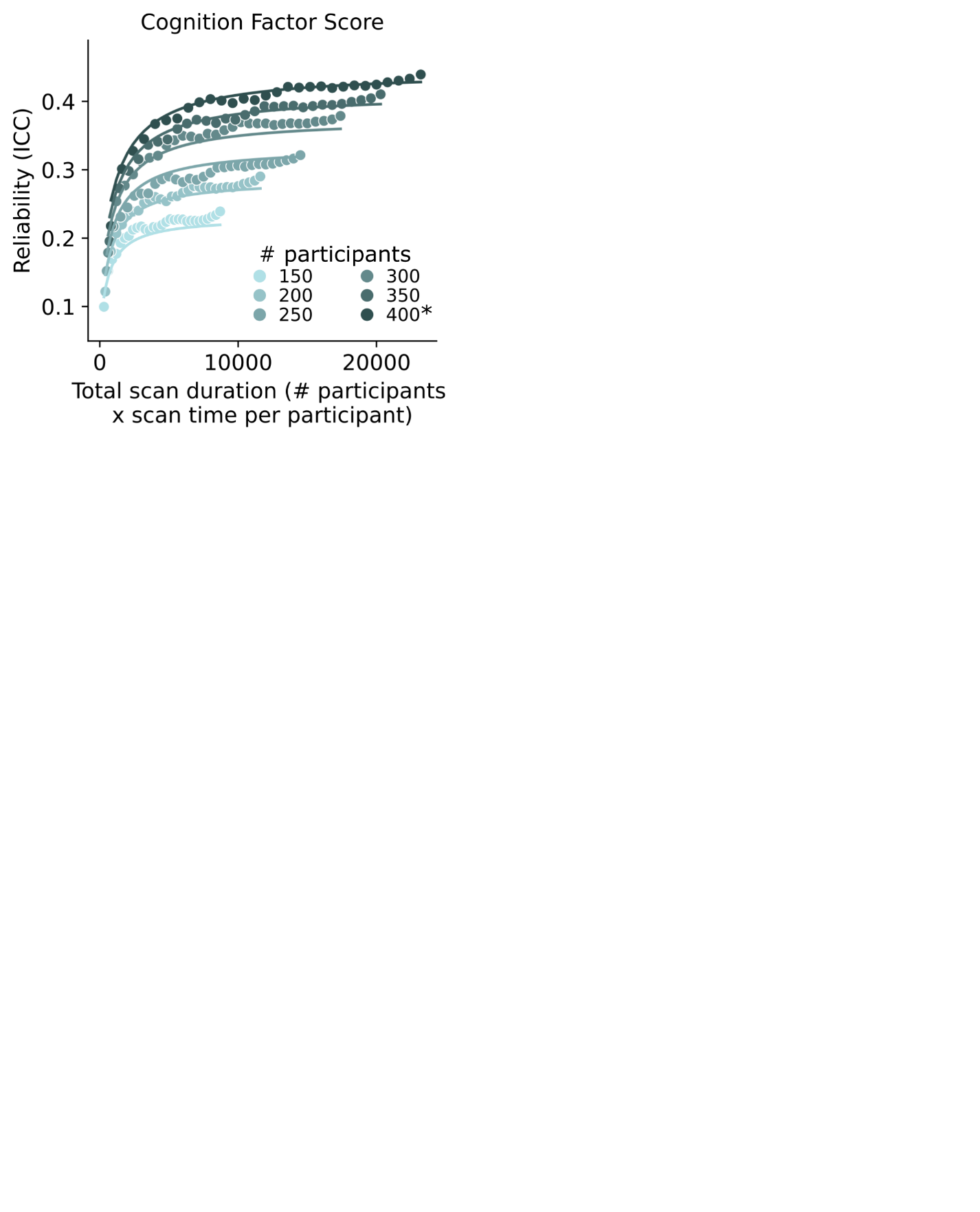

**Supplementary Fig. 27.4.** Same as Supplementary Fig. 24c except showing the scatter plots and the fit of reliability theoretical models for 1 of 19 phenotypic measures in the HCP dataset that visually follow a logarithmic pattern for prediction accuracy. Scatter plot of split-half univariate brain-wise association analyses reliability (intra-class correlation) against total scan duration in the HCP dataset. The curves were obtained by fitting a theoretical model to the reliabilities of the phenotype. The * in the figures indicates that all available participants were used, therefore the sample size will be close to, but not exactly the number shown..

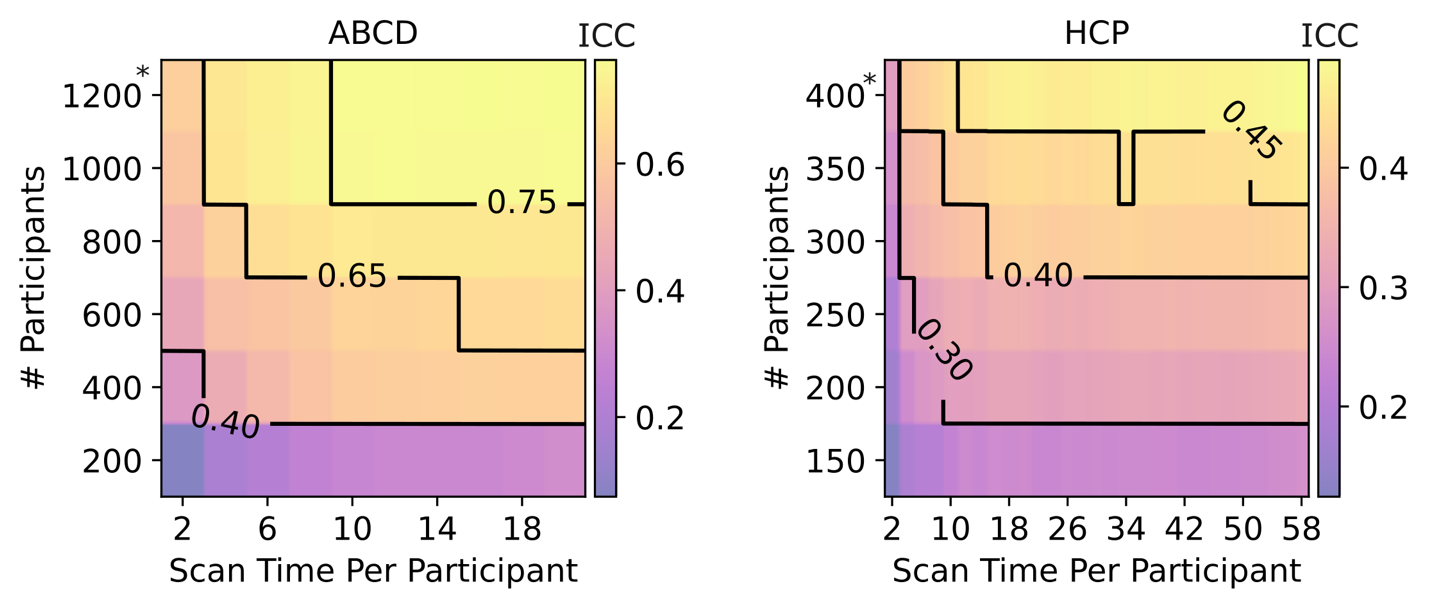

##### **Supplementary Fig. 28 |** Same as Supplementary Fig. 21b, except showing contour plot of multivariate BWAS reliability (intra-class correlation) of the cognitive factor score as a function of the scan time used to generate the functional connectivity matrix (x-axis), and the number of training participants used to train the predictive model (y-axis) in the Adolescent Brain and Cognitive Development (ABCD) and Human Connectome Project (HCP) datasets. Increasing training participants and scan time both led to increases in split-half reliability. The * in both figures indicates that all available participants were used, therefore the sample size will be close to, but not exactly the number shown.

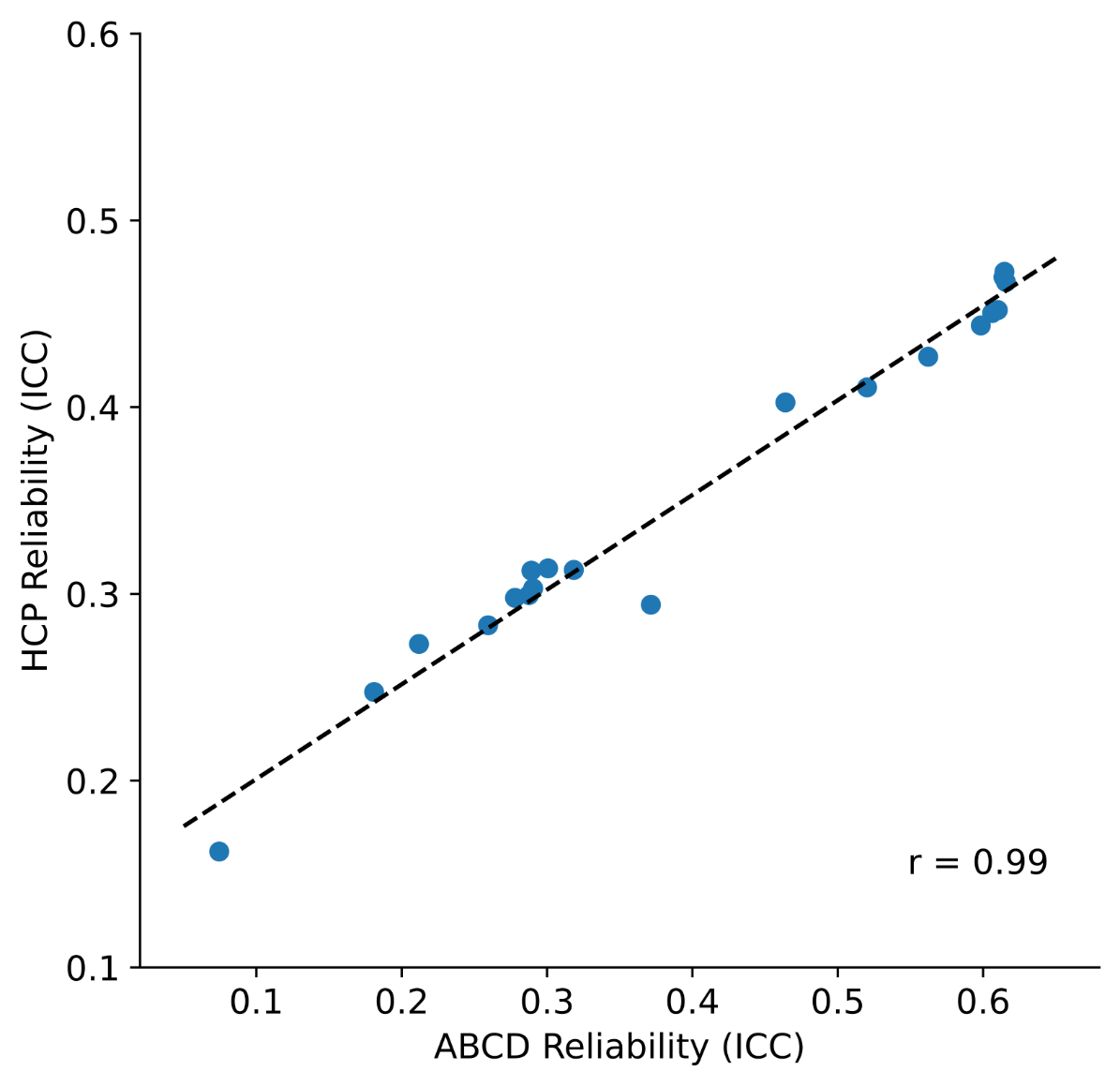

##### **Supplementary Fig. 29** | Scatter plot of the cognition factor univariate reliability (ICC) in the ABCD (x-axis) and HCP (y-axis) datasets. Each dot represents the prediction accuracy for each dataset with the same sample size and scan time per participant (extracted from Supplementary Fig. 28). Although the cognitive factor score is not comparable across datasets, we observed a strong correlation between the two datasets (r = 0.99).

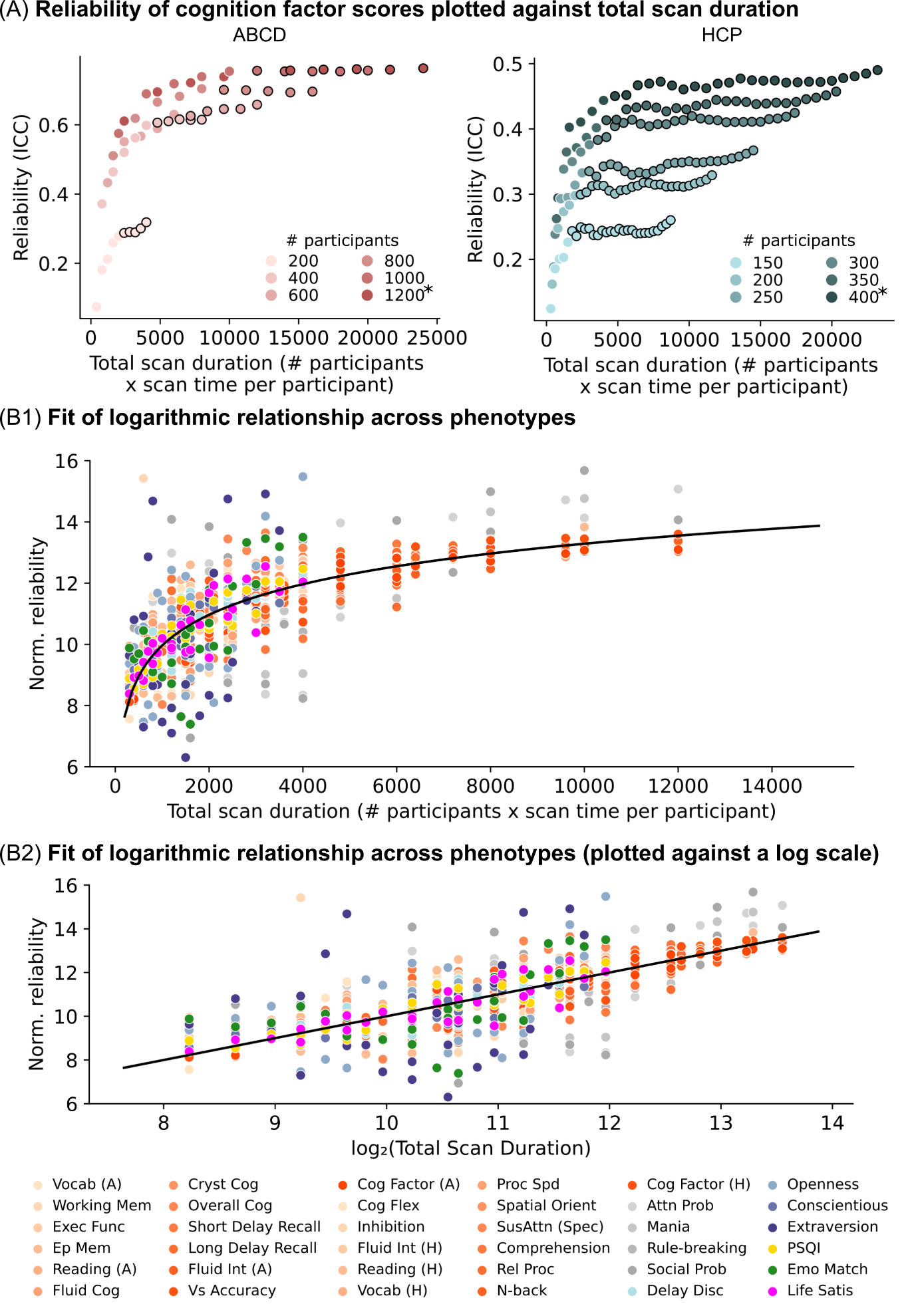

##### **Supplementary Fig. 30 | a.** Same as Supplementary Fig. 23, except showing reliability of multivariate brain-wide association (intra-class correlation) of the cognitive factor as a function of total scan duration (defined as # participants x scan time per participant). Each color represents a different number of total participants used to train the prediction algorithm. Plots were repeated for the ABCD and HCP datasets. The * indicates that all available participants were used, therefore the sample size will be close to, but not exactly the number shown. Increase in reliability observed diminishing returns (relative to sample size) when scan time per participant reached approximately 10 minutes in the HCP dataset; data points with more than 10 minutes of scan time are shown with black outlines. **b1.** Scatter plot showing normalized reliability of the cognitive factor scores and 34 other phenotypes versus total scan duration ignoring data beyond 10 minutes of scan time. Blue and red dots represent results from the ABCD and HCP datasets respectively. The logarithmic black curve suggests that total scan duration explained reliability well across phenotypic domains and datasets. **b2.** Same as panel b1, except the horizontal axis (total scan duration) is plotted on a logarithm scale. The linear black line suggests that the logarithm of total scan The linear black line suggests that the logarithm of total scan duration explained prediction performance well across phenotypic domains and datasets.
